# Nanopore microscope identifies RNA isoforms with structural colors

**DOI:** 10.1101/2021.10.16.464631

**Authors:** Filip Bošković, Ulrich Felix Keyser

## Abstract

Identifying RNA transcript isoforms requires intricate protocols that suffer from various enzymatic biases. Here we design three-dimensional molecular constructs that enable identification of transcript isoforms at the single-molecule level using solid-state nanopore microscopy. We refold target RNA into RNA identifiers (IDs) with designed sets of complementary DNA strands. Each reshaped molecule carries a unique sequence of structural (pseudo)colors. Structural colors consist of DNA structures, protein labels, native RNA structures, or a combination of all three. The sequence of structural colors of RNA IDs enables simultaneous identification and relative quantification of multiple RNA targets without prior amplification. Our **A**mplification-free **R**NA **T**arg**E**t **M**ultiplex **I**soform **S**ensing (ARTEMIS) reveals structural arrangements in native transcripts in agreement with published variants. ARTEMIS discriminates circular and linear transcript isoforms in a one step, enzyme-free reaction in a complex human transcriptome using single-molecule readout.

**One sentence summary:** Here we show enzyme-free identification and relative quantification of RNA isoforms using a nanopore microscope and structural colors.

## Main Text

Single-molecule identification of multiple transcript isoforms in parallel without preamplification is critical for understanding transcriptome diversity and gene expression networks (*1*). Identification and quantification of structural arrangements in native transcripts are both challenging, and current methods do not necessarily yield results reflecting innate transcriptome diversity (*2, 3*). Although identification of long RNA molecules is possible with existing nucleic acid detection methods (*4–6*), they lack specificity and simplicity. In addition, common approaches mainly rely on enzymatic reactions and require preamplification. These lead to inevitable biases and loss of information (*7–9*). RNA sequencing approaches require extensive and intricate adaptations to achieve the sequencing of transcript variants and to test their circularity (*2, 10,11*). These widely used techniques face amplification and reverse transcriptase biases, and detection of transcript variants is affected by short reads in RNA sequencing (*10, 11*). Recently, nanopore sequencing introduced direct RNA readout (*10*), however, access to singlemolecule information of gene expression level in combination with low-quality reads and uncertainty about the 5’end of the transcript remain major challenges (*12*). As in previously established RNA-seq methods, nanopore RNA-seq also suffers from enzymatic biases (*13,14*). Additionally, secondary structures in both RNA and complementary DNA (cDNA) contribute to biases by obstructing the binding of primers and sequencing adapters (*15*). We are in need of an enzyme-free method for targeted identification of native transcript isoforms, that avoids notoriously laborious protocols and that reveals structural arrangements on the single-molecule level.

Here, we introduce **A**mplification-free **R**NA **T**arg**E**t **M**ultiplex **I**soforms Sensing (ARTEMIS). ARTEMIS relies on the molecular design of identifiers (IDs) for native RNA targets that reshape the RNA ‘scaffolds’ into unique structures inspired by the DNA origami technique (*16*). As a first example, we introduce the concept of structural (pseudo)colors for three RNA target isoforms that are complemented with short oligonucleotides as shown in Fig. 1A. In this example, the molecular design of each isoform-specific ID is represented by a molecular pattern with two sites that have structural color ‘1’, ‘2’, or ‘3’. We create a structural color by interspacing an integer number of structural units. Each structural color has a linearly increasing molecular weight with its ‘color number’ (design details of structural colors are illustrated in Fig. S1). The basic design of the structural unit (Fig. 1B and Fig. S1) consists of a docking strand (shown in black), complementary to the specific sequence on the target RNA, and the overhang sequence (red). The imaging strand is complementary to the docking strand’s overhang and contains a structure of fixed molecular weight. For ARTEMIS, we employed monovalent streptavidin (*17*) or a DNA cuboid nanostructure (designs are depicted in Fig. S1, Fig. S2A-B, DNA cuboid oligonucleotides are listed in Table S1).

**Fig. 1.**
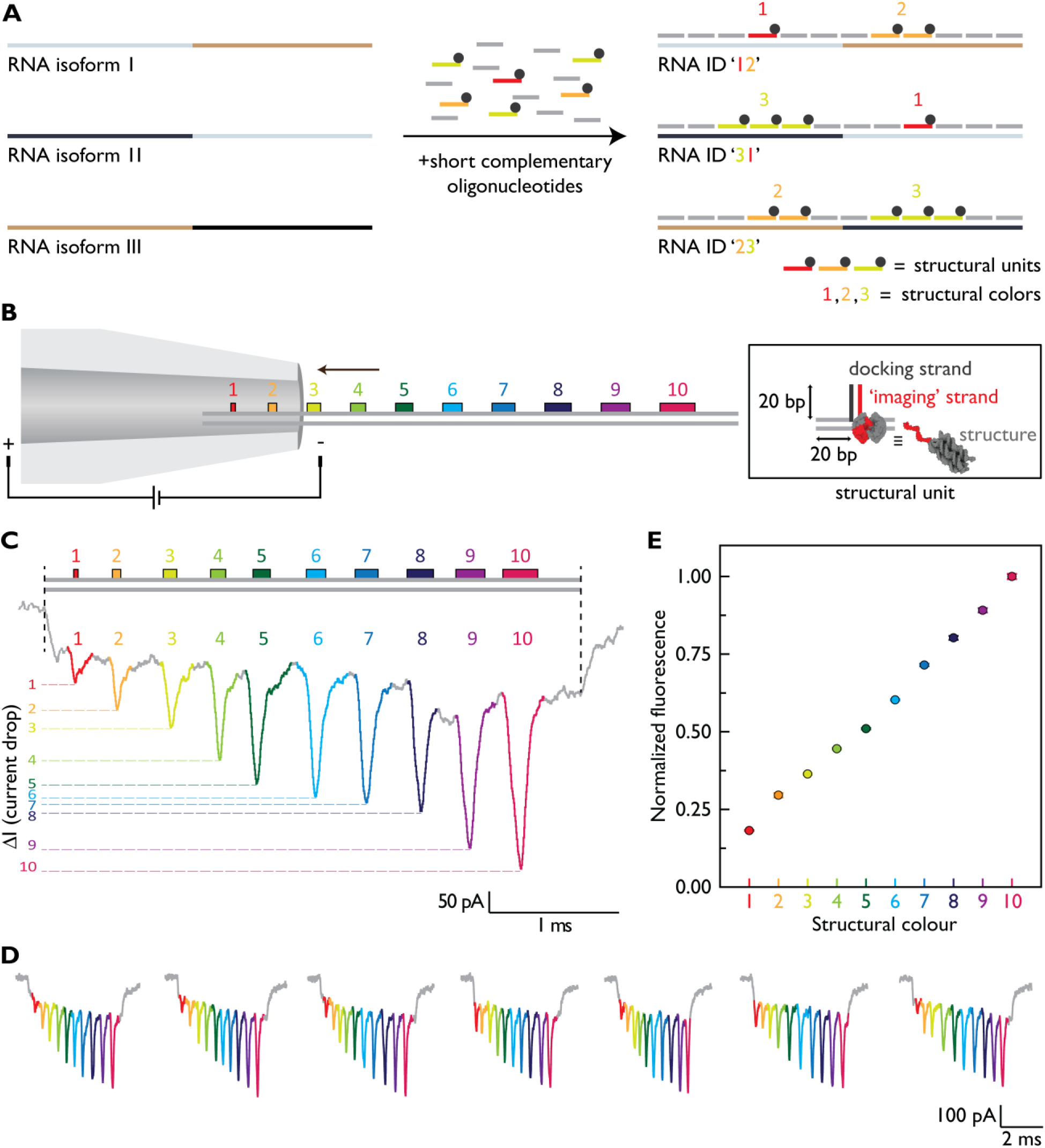
ARTEMIS identifies multiple RNA targets using structural colors and nanopore microscopy. (**A**) RNA isoform-specific identifier (ID) fabrication using structural colors (red – 1, orange – 2 and green – 3). Three RNA isoforms are labeled with an ‘exon-specific’ structural color. Each structural color is composed of an integer number of structural units. (**B**) The ID with ten different structural colors is read by passing it through the nanopore microscope. A structural color consists of an integer number of structural units (0-10) that are placed sequentially and read as one structural color. Each structural unit is composed of the part that binds to a target and has the overhang (docking strand, black) and the imaging strand (yellow) that is complementary to the overhang and has a terminal structure (monovalent streptavidin or DNA cuboid). (**C**) Nanopore microscope detects up to 10 structural colors within the same molecular ruler. An example nanopore event in which each structural color is identified by the nanopore microscope. (**D**) Singlemolecule readout of structural colors and their identity as assigned in example nanopore events. (**E**) The correct number of structural units per color is successfully verified with the fluorescently labeled (5’-fluorescein) structural units. In the plot, normalized fluorescence for 1-10 structural colors is shown. Error bars indicate a standard error of three repeats.

The sequence of structural colors in RNA ID can be conveniently read using a solid-state nanopore microscope (*18*) (Fig. 1B) as a rapid, enzyme-free, and affordable alternative for both short- and long-read sequencing. ARTEMIS with direct nanopore readout avoids the technical artifacts of RNA-seq and imperfections of motor proteins used in nanopore sequencing (*19*). RNA ID fabrication and identification do not require enzymes or preamplification, hence it offers rapid one step direct detection of native RNA targets in the whole transcriptome.

Using the well-characterized ability of nanopore microscopes to detect molecular weight (*18, 20*), we show, as in fluorescent microscopy, identification of multiple colors (Fig. 1B-E). We designed and tested structural colors with up to 10 levels as shown in the schematic in Fig.1B (4-color ID is shown in Fig. S2, Fig. S4, oligonucleotides are shown in Table S2, S3). Our ten-color palette is shown in Fig. 1B-E demonstrating the simultaneous detection of ten colors at the single-molecule level in a nanopore microscope (Fig. 1C, E). Example events and design details for the 10-color ID are shown in Fig. S3-4 (oligonucleotides are listed in Table S2, S4). Ten structural colors enable around ~10^10^ unique IDs (Fig. 1B-E), highlighting the feasibility of ARTEMIS for large-scale transcriptome profiling. We validated the fabrication of rulers with biotinylated ‘imaging strand’ using polyacrylamide gel electrophoresis (PAGE) with and without the addition of neutravidin (Fig. S4). Additionally, we verified the correct assembly of ten colors with fluorescence quenching using fluorescein (6-FAM) labeled structural units (Fig. 1E and Fig. S5A). IDs were fabricated in equimolar concentration, each containing only one color from 1 to 10 (Fig. 1E). The fluorescence output of each separate color indicates accurate fabrication of structural colors (Fig. 1E, Fig. S5B).

We next used ARTEMIS to identify various transcripts in a one pot reaction as schematized in Figure 2A. As a proof-of-concept, we created distinctive IDs in a complex nucleic acid mixture (Fig. 2A, Fig. S6) for human 18S, 28S ribosomal RNA (rRNA; oligonucleotides used for fabrication of 18S rRNA ID and 28S rRNA ID are listed in Table S5 and Table S6, respectively) and an external MS2 RNA ID control with a known concentration (oligonucleotides are listed in Table S7 and Table S8 with the design details in Fig. S7A). We successfully identified 18S and 28s rRNA in human total universal RNA (composition is listed in Table S9) and human total cervical adenocarcinoma RNA.

**Fig. 2.**
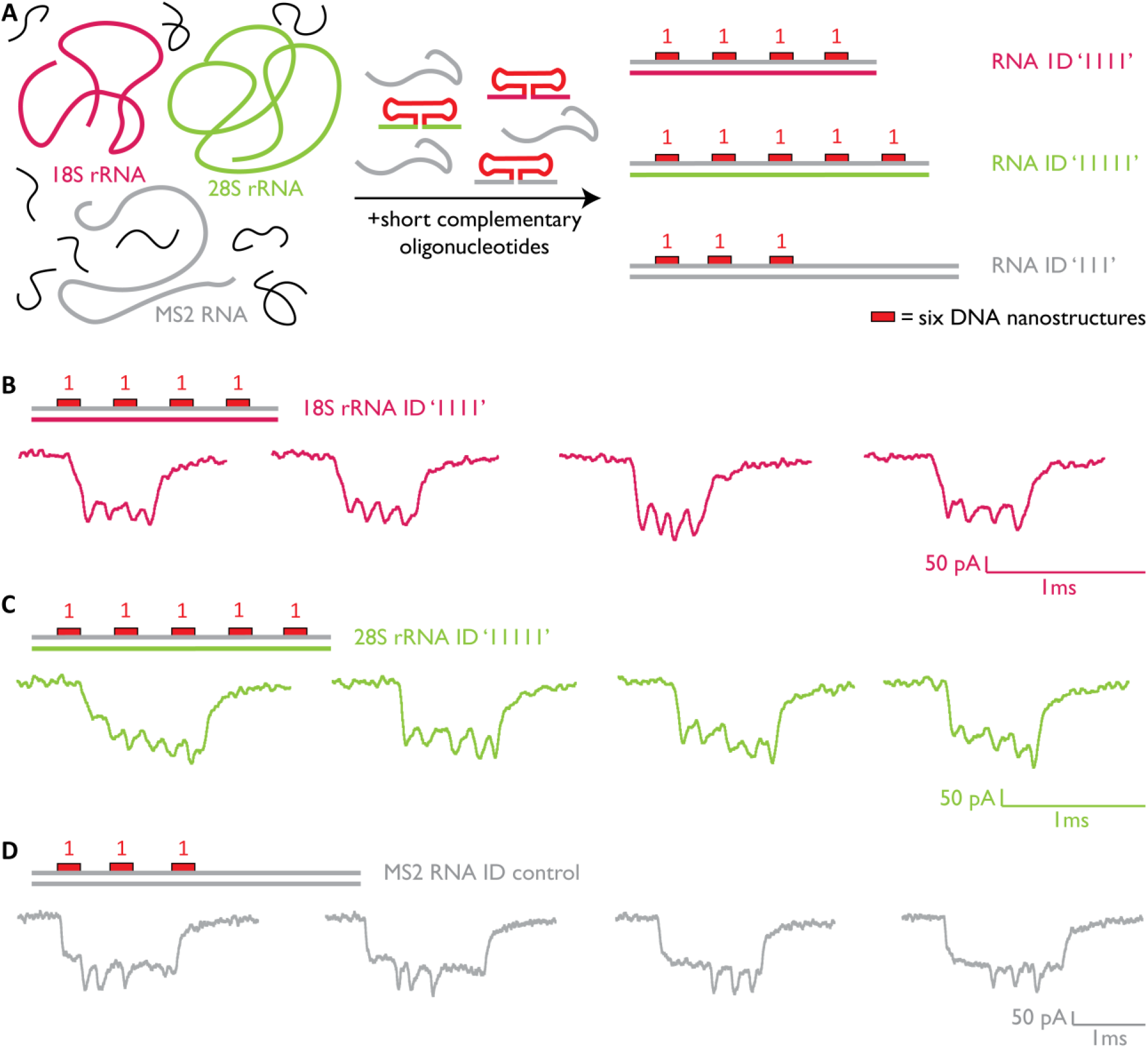
One pot reaction for multiplexed amplification-free identification of 18S and 28S rRNA. (**A**) ID fabrication and designs for multiple RNA targets in a complex mixture of human total RNA. Typical 18S rRNA ID ‘1111’ (crimson), 28S rRNA ID ‘11111’ (green), and MS2 RNA ID control (gray) as measured from a single nanopore are presented in (**B**), (**C**), and (**D**), respectively.

Each RNA ID was identified with the nanopore microscope and events for 18S rRNA ID with four sites (‘1111’), 28S rRNA with five sites (‘11111’), and external RNA ID control with three sites (‘111’) are depicted in Fig. 2B, C, and D, respectively (additional events Fig. S8). Expected velocity fluctuations during translocation (*21*) play a minor role in measuring distances between sites as we achieve correct readout and position sequencing of sites along target RNA (Fig. S7C).

Two main obstacles for general RNA analysis degradation by nucleases assisted by magnesium ions and structured regions that terminate amplification or block hybridization. The latter issue we addressed by replacing divalent with various monovalent ions. The removal of magnesium provides the added benefit to reduce RNA structure stabilization and fragmentation for RNA ID fabrication (Fig. S9). We also determined the optimum salt concentration for RNA ID fabrication in our experimental conditions (Fig. S10). The stability and purity of fabricated RNA IDs over time was assessed using nanopores and agarose gel electrophoresis (Fig. S11-12). We find that RNA IDs show no to minimal degradation with standard storage conditions, in agreement with the previous observations for RNA-DNA hybrids (*22*).

Based on the multiplexed detection, ARTEMIS discriminated transcript variants, which are a result of alternative transcript processing and structural arrangements in a premature transcript (pre-mRNA) (Fig. 3). As a proof-of-principle, we designed exons and their respective isoforms (Fig. 3). We designed a sequence of three structural colors per exon (Fig. 3A) thus creating asymmetric and isoform-specific RNA IDs (designs with example events are presented in Fig. S13 and oligonucleotides used are listed in Tables S10 and S11). ARTEMIS identified order, length, and circular isoforms (Fig. 3B, C, and D, respectively). The combination of exons results in multiple transcript isoforms with the same length but different sequences (Fig. S14). In Fig. 3B, we show three correctly identified isoforms with the same length but a different order of exons that contain either exons I and II (‘211312’), exons I and III (‘123112’), or exons II and III (‘312123’) as shown in Fig. 3B. Even more, we can clearly determine directionality of matching exons in isoforms by our asymmetric design of the three structural colors. Besides this, we demonstrate the identification of length isoforms for example of exon I (Fig. 3C, Fig. S15).

**Fig. 3.**
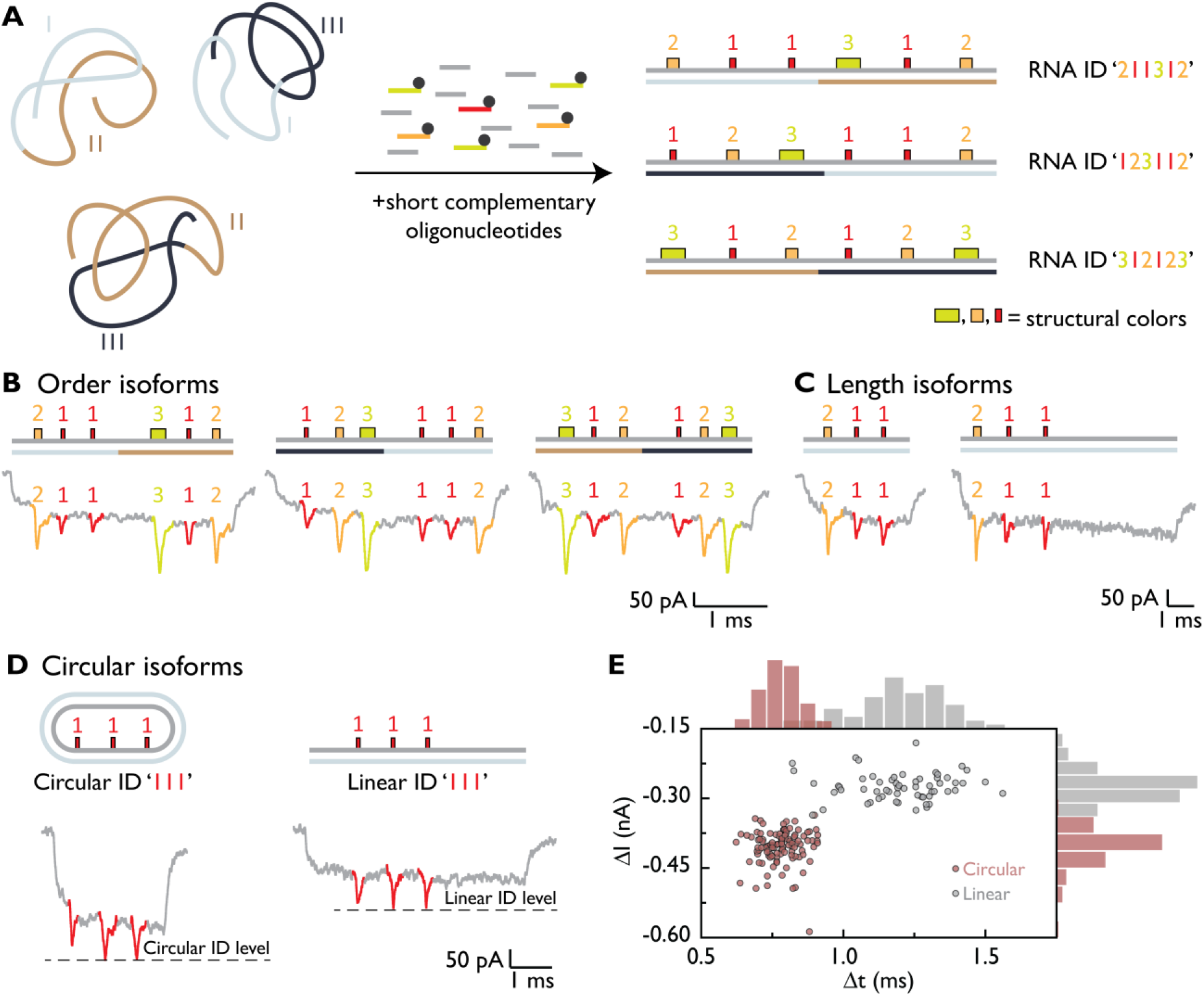
ARTEMIS discriminates engineered, alternative splicing isoforms resulting from any physical transcript arrangement. (**A**) Isoform-specific labeling is achieved by labeling each synthetic exon with an asymmetric sequence of structural colors that results in unique IDs. (**B**) Example events of the order RNA isoforms that differ in the order of structural elements i.e. synthetic exons as illustrated in (A). (**C**) RNA isoforms can differ in length and so successful discrimination of the two length isoforms is shown. (**D**) Detection of circular isoforms is shown by creating IDs on DNA scaffolds. The nanopore microscope can distinguish between circular (left) and linear configurations. (**E**) Nanopore microscope discriminates the linear and the circular populations based on the translocation time (Δt) that is ~2 times shorter for the circular isoform and event current blockage (ΔI) is ~2 times larger for the circular than for the linear ID.

Another critical feature that is hardly achievable with RNA-seq is discrimination of circular and linear isoforms (Fig. 3D-E). As a proof-of-concept, we used M13 DNA to create circular and linear IDs with the sequence of colors ‘111’ using the same oligonucleotide mixture (Fig. 3D, event examples shown in Fig. S16 while oligonucleotides are in Table S12). The scatter plot in Fig. 3E shows minimal overlap of two populations indicating circular and linear IDs ‘111’. We fixed the position of colors in the circular ID conformation by using interlock oligonucleotides (Fig. S16C-D). Circularity discrimination is also assessed by *in vitro* RNA circularization (Fig. S17) of linear MS2 RNA ID ‘111’ using T4 RNA ligase I (*23*).

Up to here, ARTEMIS has depended on the reshaping of RNA to a linear sequence of structural colors. Nevertheless, some transcripts contain strong RNA secondary structures that are challenging to remove with short oligo hybridization, or long transcripts greater than 10 kb (*15, 24*). Especially the latter transcripts would require oligos that complement the whole RNA which may be cost-prohibitive. Instead, we decided to use these parts of the RNA as alternative structural colors present in the native target. We extended ARTEMIS ID functionality by using short RNA motifs (Fig. 4A), long regions (Fig. 4B), or a combination of DNA and RNA structural colors (Fig. 4C).

**Fig. 4.**
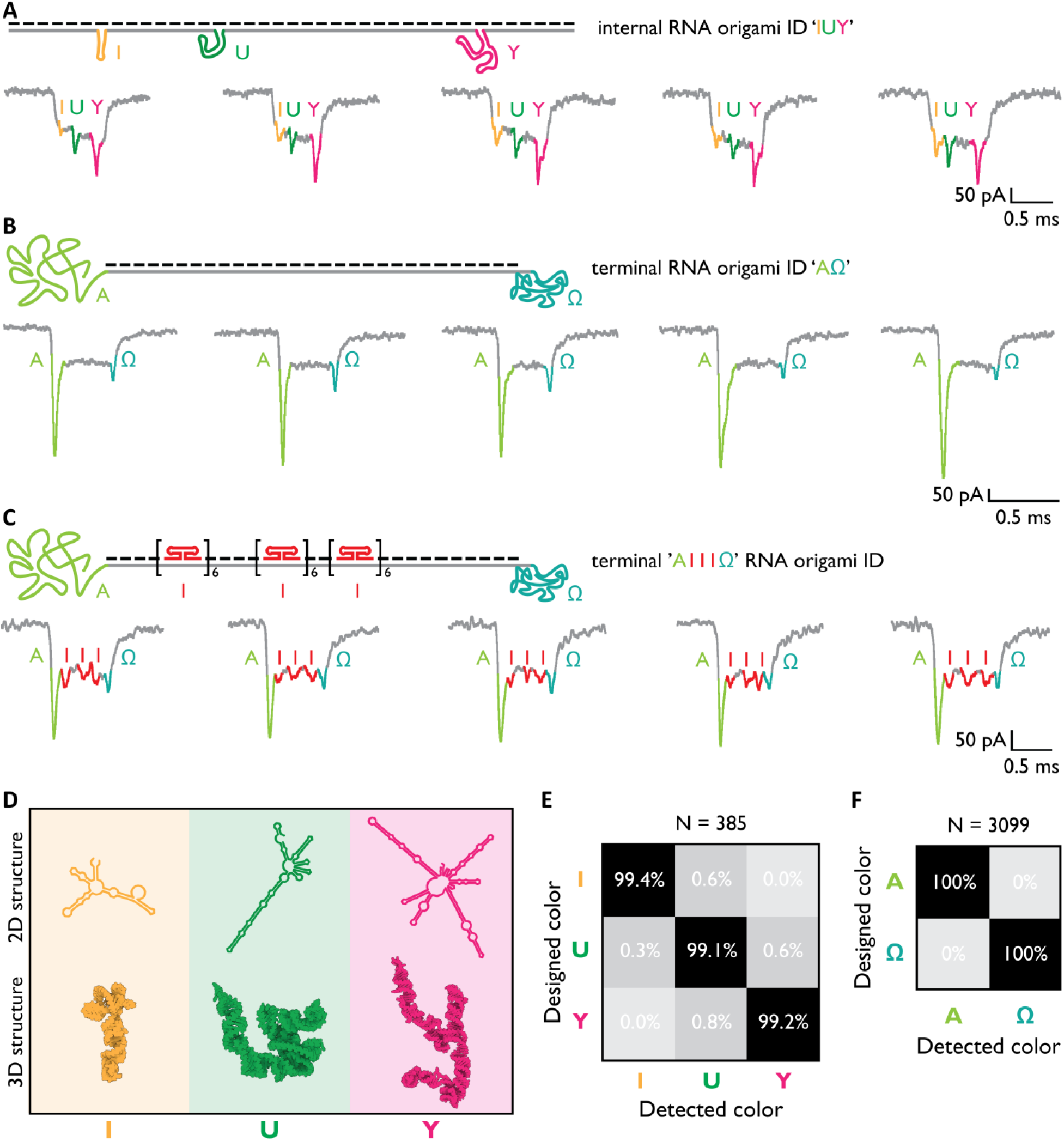
Structural colors by programmable RNA origami self-assembly. (**A**) RNA origami ID is designed to have three internal RNA origami structures at the designed locations ‘I’, ‘U’, and ‘Y’ that represent a specific structure with a unique current downward signal. (**B**) We demonstrated that ends of RNA do not need to be complemented with oligos and that can serve as terminal structural signatures. (**C**) We demonstrated that only a part of RNA is needed to assemble IDs. Here, we have self-assembled RNA terminal structures (A in green, and Ω in turquoise), RNA-DNA hybrid origami (gray black), and self-assembled DNA double-hairpins (red). (**D**) Predicted 2D and 3D structures of designed RNA origamis to match letters ‘I’, ‘U’, and ‘Y’. (**E**) Heatmap indicates correct identification of ‘I’, ‘U’, and ‘Y’ with 99.4 %, 99.1 %, and 99.2 % accuracy, respectively. Sample size is 385 nanopore events. (**F**) We show that ‘A’ and ‘Ω’ colors are identified with ~100 % accuracy. Sample size is 3099 nanopore events.

More specifically, we assembled RNA origami IDs by employing secondary structure formation in pre-designed locations (Fig. 4A; oligos are listed in Table S13). Three structural colors have been assembled by nanoscale folding of 114 nt, 190 nt, and 342 nt long single-stranded RNA to form structural colors ‘I’, ‘U’, and ‘Y’, respectively (predicted 2D and 3D structures (*25*) are shown in Fig. 4D; more details in Fig. S18). As above, each self-assembled RNA ‘origami’ has a specific current signature, that can be identified from nanopore events as shown in Fig. 4A (additional events are presented in Fig. S19). The accuracy of each structural color identification is over 99 % as displayed in the summary in Fig. 4E.

Interestingly, RNA IDs can even be realized when only the middle part of a long RNA is linearized as shown in Fig. 4B (oligonucleotides are listed in Table S14). The two terminal RNA structures are 401 nt and 1230 nt in length (‘A’ and ‘Ω’, respectively). The asymmetry in the RNA ID is directly obvious during RNA ID translocation through a nanopore. The two terminal downward signals directly correspond to the terminal RNA colors ‘A’ and ‘Ω’ (additional events are presented in Fig. S20) with accuracy in all unfolded identification of ~100 % (Fig. 4F). Finally, we designed a combination of ID ‘111’ with a terminal RNA origami on both ends as shown in Fig. 4C. The RNA ID traces in Fig. 4C show that DNA and RNA structural colors can be combined for RNA IDs. Hence, ARTEMIS can deal with secondary structures, RNA length and accurate readout is possible for any combination of structural colors. It is important to note that the addition of colors expands the number of unique RNA IDs.

Finally, we use ARTEMIS for the targeted identification of enolase (*ENO)* isoforms in commercially available human cervix adenocarcinoma total RNA (Fig. 5A-B). The *ENO* gene is known to have multiple transcript isoforms that differ in length or sequence as a result of alternative splicing of pre-mRNA (*26, 27*). We employed three colors to identify four transcript isoforms (Fig. 5A, oligonucleotides are listed in Table S15). RNA isoform ID designs and example events are illustrated in Fig. 5A. We determined the expression level of each *ENO1* transcript isoform (Fig. 5B). An internal reference ID can further improve transcript isoformlevel quantification (more details in Fig. S6) and so we chose 18S rRNA as an intersample reference (*28*). We confirmed that the nanopore event frequency is independent of the level of complementarity between target RNA and oligonucleotides (Fig. S21). The absolute concentration may be calculated from the RNA ID nanopore event frequency using a previously introduced model (*29*).

**Fig. 5.**
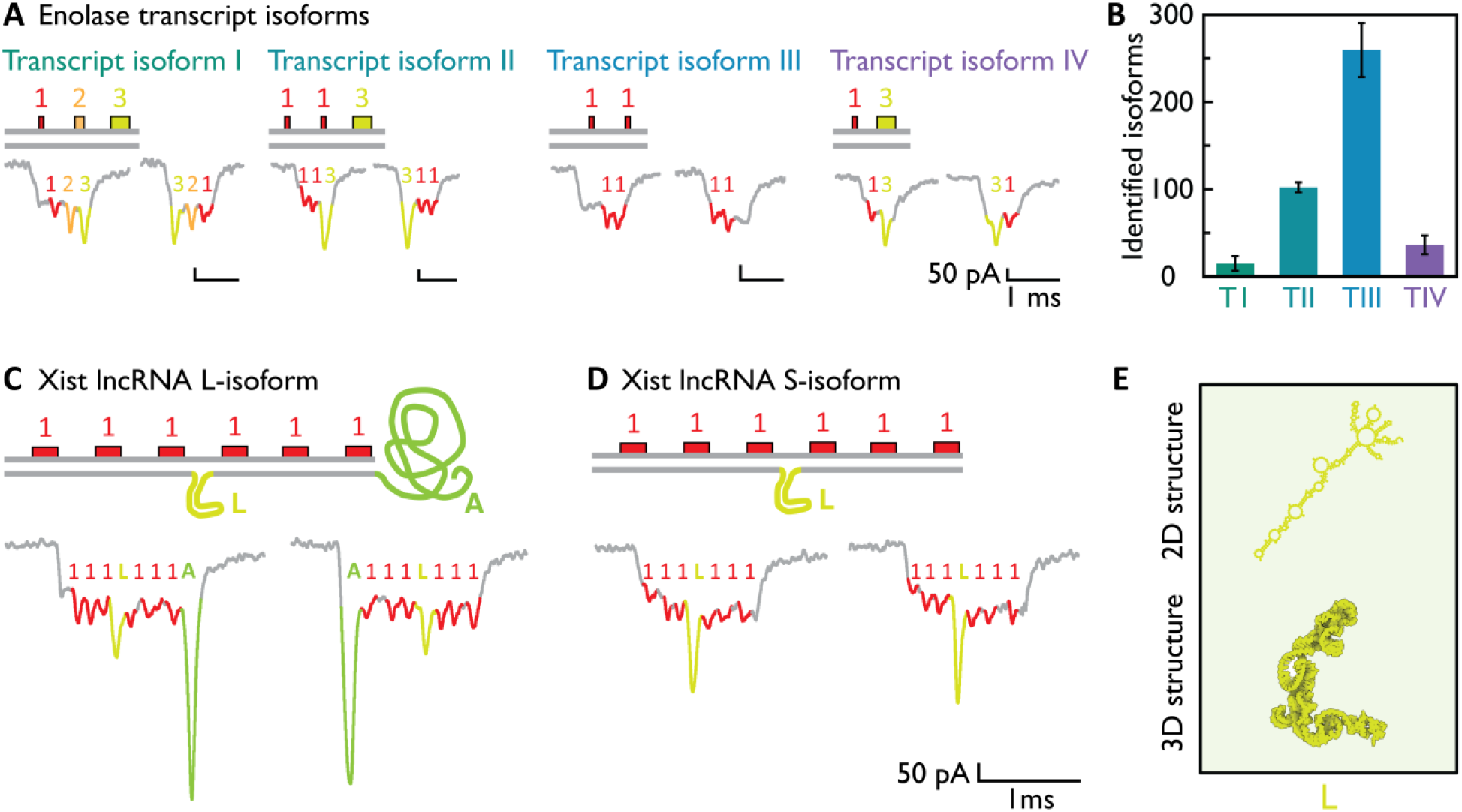
ARTEMIS discriminates alternative splicing isoforms of messenger RNA and long noncoding RNA in a complex human transcriptome mixture. (**A**) Four identified *ENO*-like transcript isoform ID designs and example nanopore events are shown. (**B**) Detected events for each *ENO1* transcript variant for three individual nanopore measurements over 20 hours with total of 39521 detected events. 18S rRNA ‘1111’ was used as internal control with 107±12 events/h. (**C**) *Xist* lncRNA L-isoform ID depicts six sites with color ‘1’, middle color ‘L’, and terminal RNA coil ‘A’. Nanopore events match their ID as depicted with colors above the downward signals. (**D**) S-isoform ID has the same design except for the terminal RNA coil that represents difference in length between these two isoforms. (**E**) Both isoforms have between third and fourth site an internal self-assembled RNA origami indicated with color ‘L’. Predicted secondary (2D) and tertiary (3D) structures of color ‘L’ are shown.

As further demonstration, we use ARTEMIS to target X-chromosome inactivation transcript long-non-coding RNA (*Xist* lncRNA). Here we use terminal RNA, internal RNA motifs, and DNA structural colors to identify length isoforms in the native transcriptome (Fig. 5C-E). We targeted part of *Xist* RNA to fabricate ID ‘111111’ (design of *Xist* lncRNA ID is schematized in Fig. S22, and oligonucleotides used for its fabrication are enumerated in Table S16). The part of the sequence that differs among long (L-isoform) and short (S-isoform) isoforms is left unpaired (Fig. 5C). The expected ID readout should depict the six sites with a structural color ‘1’, the terminal unpaired RNA coil ‘A’, and an internal self-assembled RNA origami color ‘L’ as predicted from the sequence (Fig. 5C-E). We show typical examples of *Xist* lncRNA isoform IDs that match the predicted design and previously identified *Xist* lncRNA isoforms (Fig. S22) (*27, 30*).

In this study, we introduced ARTEMIS, an approach that reshapes an RNA target into a sequence of structural colors that we call an ID, using sub-nanometer precision of DNA nanotechnology (*31, 32*). ARTEMIS omits amplification and enzyme-based steps and identifies multiple native RNA transcripts and alternative splicing variants in parallel using the nanopore microscope. As an electric measurement, a nanopore microscope has a spatial resolution comparable to complex optical microscopies with higher throughput and straightforward origami assembly (*18, 31*).

Most diseases are classified by a change of a few transcripts. Thus, accurate identification of RNAs of interest has to bypass prior amplification and reverse transcription biases (*7, 11*). Our approach has the potential to identify extensive RNA diversity from a gene of interest without the need to align to reference transcriptomes. It is known that reference transcriptomes neglect unreferenced RNA diversity, and hence the extensive RNA variation is lost (*33*). ARTEMIS is complementary to RNA sequencing-based approaches as RNA IDs rely on RNA or genomic sequence information. We demonstrated multiple isoform identification by using the same oligo mix to identify order isoforms, length isoforms as well as transcript circularity. ARTEMIS features are promising for characterizing therapeutic RNA uniformity and circularity using minimal sample amounts (*2, 34*).

We employed RNA origami self-assembly (*35–37*) as an additional way to identify transcripts of interest or to overcome some challenges of RNA analysis. Transcript IDs are assembled by using stable RNA structures as structural colors that can be either within the molecule or at its ends.

We believe that ARTEMIS opens new avenues for single-molecule mapping of RNA motifs. In addition, our study demonstrates that highly abundant natural RNAs may serve as scaffolds for DNA origami assembly with a wider length range and yield (*35*).

Our multicolor palette paves a way towards targeted isoform profiling in the whole transcriptome that excludes enzymatic and amplification biases. ARTEMIS has the potential to create ~10^10^ unique RNA IDs that are readable using nanopore microscopy or imaging methods that rely on super-resolution microscopy including DNA-PAINT.

## Acknowledgments

We are thankful to Marcus Fletcher and Jinbo Zhu for the critically reading of the manuscript and useful suggestions.

## Author contributions

F.B. conceived the idea, F.B. and U.F.K. designed the study. F.B. performed experiments and analyzed the data. F.B. and U.F.K. wrote the manuscript.

## Supplementary Materials

Materials and Methods

Figures S1-S22

Tables S1-S16

References

