## Supplementary Information - materials and methods, figures and tables for "Nanopore microscope identifies RNA isoforms with structural colors"

**This PDF file includes:**

Materials and Methods
Supplementary Text
Figs. S1 to S22
Tables S1 to S16

### Materials and Methods

Materials

The commercial buffers used in this study were Tris-EDTA buffer solution 100 × concentrate (Sigma-Aldrich, catalog number T9285), 0.2 µm filtered 1M $MgCl_2$ (Invitrogen by Thermo Fisher Scientific, catalog number AM9530G), 0.2 µm filtered and autoclaved nuclease-free water (Ambion, catalog number AM9937). Lithium chloride for molecular biology ≥ 99% purity (Sigma-Aldrich, catalog number L9650), sodium chloride for molecular biology ≥ 99% purity (Sigma-Aldrich, catalog number S3014), Tris-HCl BioPerformance certified, ≥ 99% purity (Sigma-Aldrich, catalog number T5941). All buffers used in this study were filtered with 0.22 µm Millipore syringe filter units (Merck).

Glass quartz capillaries with filament (inner diameter 0.2 mm, outer diameter 0.5 mm) were purchased from Sutter Instrument Company. PDMS was purchased from Sylgard 184, Dow Corning (catalog number 101697), microscope slides clear ground 1.0 – 1.2 mm (Thermo Fisher Scientific, catalog number 1238-3118), silver wire with 1.0 mm diameter (Advent Research Materials Ltd, catalog number AG548711). Amicon 0.5 mL filter units (100 kDa cut-off) were purchased from Merck (catalog number UFC5100BK). Membrane Filter, 0.22 µm pore size membrane filters (MF-Merck Millipore™, catalog number GSWP04700).

DNA LoBind® Tubes (Eppendorf) were purchased from Thermo Fisher Scientific, and thin-walled, frosted lid, RNase-free PCR tubes (0.2 mL) were purchased from Thermo Fisher Scientific (catalog number AM12225).

RNA from bacteriophage MS2 3569 nt in length (Roche, catalog number 10165948001), total RNA from human cervical adenocarcinoma (Thermo Fisher Scientific, Invitrogen, catalog number AM7852) and human universal reference total RNA (Thermo Fisher Scientific, Invitrogen, catalog number QS0639).

Single-stranded circular m13mp18 7249 nt in length (Guild Biosciences, foundation m13).

DNA cuboid

DNA cuboid is assembled by using six oligonucleotides provided in Table S1. We mixed 1 µL of each oligonucleotide (100 µM, IDTE buffer (10 mM Tris-HCl, 0.1 mM EDTA), pH = 8.0), 10 µL of filtered 10 × TE buffer (100 mM Tris-Cl, 10 mM EDTA, pH = 8.0), 20 µL of filtered 100 mM $MgCl_2$, and 64 µL of filtered Milli-Q ultrapure water. Buffers are filtered with the MF-Millipore™ Membrane Filter, 0.22 µm pore size. The mix is vortexed and spun down before the structure assembly. All oligonucleotides were purified by desalting and ordered in IDTE buffer in 100 µM concentration. Further details of DNA cuboid assembly can be found as CP3 short DNA origami nanopore (1, 2) without additional structural changes required for the structural unit.

The mix was heated to 95 °C for 5 minutes and slowly cooled down to 25 °C over 18 h. The mix was stored at 4°C without additional purification until further use.

The DNA cuboid for fluorescence/quenching assay was assembled using the same protocol. Oligonucleotide 1M1 is replaced with the 5' labelled end of oligo 1M1 with 6-FAM (100 µM, IDTE buffer (10 mM Tris-HCl, 0.1 mM EDTA), pH = 8.0). The 6-FAM 1M1 oligonucleotide was purified with high-performance liquid chromatography (HPLC).

Design of structural unit and structural colors

The structural unit is composed of a docking strand and an imaging strand. The docking strand has two parts. The first part is a 20 nt sequence that is complementary to the specific position in a target RNA, and the second part is a 20 nt sequence that is complementary to the imaging strand. The imaging strand harbors at the 3' end a structure (Fig. S1A, left). This structure can be a protein such as monovalent streptavidin bound to biotin or DNA cuboid (Fig. S1B).

Structural colors are made by the sequential placing of an integer number of structural units. For instance, structural color two corresponds to two structural units (Fig. S1A, right). In this study, we demonstrated the fabrication of ten structural colors (eleven including structural color 0). We used streptavidin-based structural colors for data shown in Fig. 1, and DNA cuboids for data shown in Fig. 3 and Fig. 5.

For the one-color system (as shown in Fig. 2, Fig. S8), we only varied the number of structural colors but not the structural color itself. Here, we used six DNA double-hairpin structures to induce a signal in the nanopore microscope previously extensively used for this purpose. (3–5)

Fabrication of 4-color and 10-color IDs

To fabricate multicolor IDs we used the oligonucleotide mixes that contain docking strands and imaging strand (oligonucleotides used to complement the whole target are listed in Table S2 and oligonucleotides replaced for 4-color and 10-color IDs are in Table S3 and Table S4, respectively). We prepared 40 µL reaction by mixing linearized ssDNA (to 20 nM or 800 fmoles) and oligonucleotides (to 60 nM each or 2400 fmoles), in 10 mM $MgCl_2$, 1 × TE (10 mM Tris-HCl, 1 mM EDTA, pH 8.0) buffer, and nuclease-free water was added to the final reaction volume.(6) Buffers are filtered with the MF-Millipore™ Membrane Filter, 0.22 µm pore size. The mix is mixed by pipetting and spin down. The mixture was heated up to 70 °C for 30 s and after gradually cooled down (-0.5 °C/cycle, 90 cycles each 30 s) over 45 minutes to room temperature, and hold at 4 °C. Terminal oligonucleotides contain four dT nucleotides that should prevent IDs base stacking. (3, 7, 8) 4-color and 10-color designs are illustrated in Fig. S1 and S3, respectively. Biotin 'imaging' strand in oligo mix was in 1.5 × excess to docking sites. The fabrication is performed as mentioned in previous sections.

Native agarose gel electrophoresis analysis

Samples were run on a 1% (w/v) agarose gel prepared in fresh 1 × TBE buffer in autoclaved Milli-Q water for 90 minutes, at 70 V on ice. We loaded 150 ng or otherwise indicated for each RNA sample and used fresh 1 × purple loading dye without SDS (NEB). The gel was poststained in 3 × GelRed buffer (Biotium) and imaged with a GelDoc-It™(UVP).

Gel images were processed using ImageJ (Fiji) (*9*) by inverting grayscale and subsequent homogenous background subtracted with 100-150 pixels rolling ball.

Native agarose gel electrophoresis analysis of the molecular 4-color and 10-color rulers

4-color and 10-color molecular rulers were filtered using 0.5 mL 100 kDa cutoff Amicon filter units. The washing buffer used for filtration is composed of filtered 10 mM Tris-HCl (pH = 8.0), 0.5 mM $MgCl_2$. All samples were pre-mixed with 6 × purple loading dye without sodium dodecyl sulfate (SDS) purchased from NEB. 1 × loading dye components are 2.5% Ficoll®-400, 10 mM EDTA, 3.3 mM Tris-HCl, 0.02% Dye 1, 0.001% Dye 2, pH 8 at 25°C. Additionally, the samples are mixed with filtered 10 × buffer to 1 × TBE buffer (Tris-borate-EDTA). The amount of loaded nucleic acids per well was aimed to be from 80-150 ng. All comparable samples were added in the same volume to prevent a salt difference-driven shift.

As it is shown in Fig. S4 two molecular IDs with 4 colors and 10 colors (lanes 2 and 3 respectively) with biotin-imaging strand without added neutravidin have expected shift from the single-stranded form. Even more, 10-color molecular ruler runs slightly slower than a 4-color ruler as expected from design, since a 10-color ruler has 45 more structural units (forming structural colors 5, 6, 7, 8, 9, 10) with each having 23 bp and 3 nt dT linker. The 4-color and 10-color ruler samples incubated with 10 times excess of neutravidin (ThermoFisher Scientific, catalog number 31050) prior to the PAGE are shown in lanes 5 and 6, respectively. Both rulers were significantly shifted after the addition of neutravidin (lanes 5 and 6) in comparison to the rulers without neutravidin added. 1kb DNA ladder (NEB, 10 mM Tris-HCl, 1 mM EDTA, pH= 8.0 at 25°C) clearly indicates the expected running speed of the molecular rulers.

Fluorescence-quenching assay for validation of structural color assembly

We assembled 10 different molecular rulers where each has only one structural color from 1 to 10. In this case, as a structure we employed a 5’ 6-FAM labeled DNA cuboid.

Firstly, 20 µL of a molecular ruler mix (20 nM) after assembly was mixed with 15 µL of 6-FAM labeled DNA cuboid (1 µM), filtered 4 µL of 1M NaCl, and 2 µL 100 mM $MgCl_2$ for 2 h at room temperature. After the incubation of the DNA cuboid with a molecular ruler, we added 1 µL the complementary strand with a 3’ Iowa Black fluorescent quencher (100 µM) and incubated it for 1-2 h (Fig. S5A). Iowa Black® quencher is known to quench well 6-FAM(10) since it has broad absorbance spectra ranging from 420 to 620 nm with a peak absorbance at 531 nm (according to Integrated DNA Technologies lnc).

We vortexed and spun down the mixes after final incubation with quencher strand and diluted it with 38 µL of the filtered washing buffer (10 mM Tris-HCl (pH = 8.0), 0.5 mM $MgCl_2$). The spectra are recorded with the Cary Eclipse fluorescence spectrophotometer with Peltier thermostat multicell holder and temperature controller (Agilent) using a glass quartz cuvette. Fluorescent intensity was recorded at the excitation wavelength of 495 nm (absorbance max) and emission spectra are obtained in a range from 500 to 650 nm with the emission max at 520 nm (Fig. S5B). All measurements were recorded at room temperature (20 °C). The measurements for each sample were repeated three times and error bars are presented as a standard error (Fig. 1D).

MS2 RNA ID fabrication using a part and the whole RNA sequence

We assembled MS2 RNA ID using MS2 RNA (Roche through Sigma-Aldrich, catalog number 10165948001). Short complementary oligonucleotides 32-48 nt in length were annealed to the part of MS2 RNA (Table S7) to fabricate MS2 RNA ID ‘111’ partially complementary ID (MS2 RNA ID ‘111’p) as illustrated in Fig. S7A, while for the fully complementary ID (MS2 RNA ID ‘111’f) additional short complementary oligonucleotides were added (Table S8). The six interspaced DNA double-hairpin protrusions (labeled as ‘1’) are used to induce a current signal detectable with a nanopore microscope. The distance between color positions were designed to be $\Delta t_1$ = 374 nt (~127 nm) and $\Delta t_2$ = 488 nt (~166 nm). These two distances were successfully discriminated as shown in nanopore events and by the positional analysis (Fig. S7B-C). We also show the dependence of event frequency on the concentration of 8 kb DNA (Fig. S7D).

RNA ID fabrication in a complex mixture of human total RNA

We prepared 40 µL reaction by mixing human total RNA (to 12.5 ng/µL) and oligonucleotides specific for all RNA targets (to 60 nM each), in 10 mM $MgCl_2$ (or 100 mM LiCl), 1 × TE (10 mM Tris-HCl, 1 mM EDTA, pH 8.0) buffer, and nuclease-free water was added to the final reaction volume. Buffers are filtered with the MF-Millipore™ Membrane Filter, 0.22 µm pore size. The reaction is mixed by pipetting and spun down. The mixture was heated up to 70 °C for 30 s and after gradually cooled down (-0.5 °C/cycle, 90 cycles each 30 s) over 45 minutes to room temperature, and hold at 4 °C.

Two samples were used for studying RNA identification in a complex mixture i.e. background of total RNA. The first one was human universal reference RNA (Invitrogen, catalog number QS0639) that represents a pool of total RNAs from ten different human cell lines/tissues (as listed in Table S9) that were DNase-treated. Another total RNA originates from cervical adenocarcinoma (HeLa-S3; Invitrogen, catalog number AM7852). Both total RNAs were diluted in nuclease-free water (ThermoFisher) to the final concentration of 100 ng/µL, aliquoted, and stored at -20 °C for short-term use or -80 °C for long-term storage.

For data shown in Fig. 2, we verified that oligonucleotide mixes for 18S rRNA, 28S rRNA, and *Xist* lncRNA with M13 and MS2 controls can be assembled in a single-pot reaction. For Fig. 5 we verified the assembly of oligonucleotide mixes for *ENO1* (oligonucleotides listed in Table S15) and *Xist* lncRNA (oligonucleotides listed in Table S16).

RNA ID temperature storage conditions

We assembled MS2 RNA ID ‘111’p and stored it at 4 °C or -20 °C for 1, 4, and 8 days (Fig. S11). IDs were run on 1% (w/v) agarose gel (SigmaAldrich, BioReagent for molecular biology, low EEO; catalog number A9539) prepared in 1 × TBE buffer, and cooked in the microwave oven for three minutes and after boiling were stirred and returned. The gel was cooled down under running water, poured, and cast for 1 h at room temperature (20 °C). The samples were run on the gel in 1 × TBE buffer for 3 h, 70 V, in the ice water bath. The gel was post-stained in 3 × GelRed® in water (Biotium, catalog number 41001) for 10 minutes. The gel was imaged with a UVP GelDoc-It™ imaging system. Gel images were post-processed with an image processing package Fiji (ImageJ). All gel display colors were inverted, and the contrast and brightness were adjusted. The subtract background function was equally applied to the whole gel image using a rolling ball radius of 100-150 pixels (light background).

All samples have shown a similar band running on the gel without visible difference also in the nanopore events (Fig. S11A-B).

Temperature and salt type effects on the RNA ID fabrication

We assembled M13 ID ‘11111’ and MS2 ID ‘111’p using either 10 mM $MgCl_2$ or 100 mM of monovalent salts (LiCl, NaCl, or KCl) with two temperature regimes (starting at 70 °C or 85 °C and gradually cooling to room temperature) as shown in Fig. S9. Nanopore events for both M13 and MS2 and both temperature regimes look as designed. The agarose gel prepared as previously described confirms the correct ID fabrication. However, in the condition with $MgCl_2$ at 85 °C the gel indicates that almost all RNAs are fragmented and in nanopore events, only a few events were detectable for a 2 h measurement time due to the magnesium fragmentation. (*11*) Also, M13 IDs assembled with magnesium show significant aggregation that is even more prominent at 85 °C (Fig. S9E, lanes 2 and 6). This indicates that magnesium can be omitted from the ID fabrication step. Hence, eliminating magnesium fragmentation (*11*) and nuclease-degradation of RNA that relies on magnesium ions. (*12*)

Salt concentration effects on the RNA ID fabrication

We assembled MS2 ID '111'f using either $MgCl_2$ or LiCl at various concentrations (at 70 °C temperature regime). For magnesium we used 2.5 mM, 5 mM, or 10 mM $MgCl_2$, and for lithium we used 25 mM, 50 mM, or 100 mM LiCl (Fig. S10). It can be observed that RNA IDs are assembled under all magnesium concentrations while at 25 mM LiCl RNA ID was not fabricated. The difference in band intensity might be due to the variable amount of recovered RNA IDs after the Amicon filtration.

Enrichment of RNA IDs from a complex sample

We established an RNA ID enrichment protocol that depletes background <100 kDa single-stranded nucleic acids (Fig. S12) to further decrease background in nanopore measurements.

The enrichment of RNA IDs after fabrication was performed by employing Amicon 0.5 mL filters with 100 kDa cut-off using filtered washing buffer (0.5 mM $MgCl_2$, 10 mM Tris-HCl pH 8.0). RNA ID (40 µL reaction) was filtered with washing buffer (460 µL) two times for 10 minutes, 9,200 × g at 3 °C. The sample is collected from the tube after enrichment removal and kept on ice until further experimental steps.

Synthetic exons fabrication

Synthetic exons that mimic exons as units that undergo alternative splicing are designed as follows. Each synthetic exon is characterized by a unique three-color site ID with 20 nt terminal overhangs (Fig. S13). We employed 3.6 kb RNA as a unit length measure and fabricated four different exons. The exon I has ID '112' (Fig. S13A) with terminal ends A and B' (each 20 nt in length). The exon II has ID '312' (Fig. S13B) with terminal ends A' and B' (A and A' i.e., B and B' are complementary end sequence pairs). The exon III has ID '321' (Fig. S13C) with terminal ends A' and B (each 20 nt in length). The exon IV (extended RNA; Fig. S13D) is designed to not carry structural colors and it only has the A' terminal end sequence. These synthetic exons are characterized by asymmetric ID designs that demonstrate not only the identification of targeted exons but also their directionality. Both are important features for accessing results of alternative processing of transcript.

The core oligonucleotides used for the fabrication of synthetic exons are listed in Table S10 (in red are shown oligonucleotides that are replaced with the structural unit docking strand). Oligonucleotides replaced from the core oligonucleotide mix for the fabrication of exon I, exon II, exon III, and exon IV are listed in Table S11.

We prepared 40 µL reaction for RNA ID fabrication by mixing RNA sample (20 nM for known target MS2 RNA concentration or 800 fmoles) and oligonucleotides (60 nM each or 2400 fmoles) where some of them contain the structural units, in 10 mM $MgCl_2$, 1 × TE (10 mM Tris-HCl, 1 mM EDTA, pH 8.0) buffer, and nuclease-free water was added to the final reaction volume. Buffers are filtered with the MF-Millipore™ Membrane Filter, 0.22 µm pore size. The reaction was mixed by pipetting and spin down. The mixture was heated up to 70 °C for 30 s and after gradually cooled down (-0.5 °C/cycle, 90 cycles each 30 s) over 45 minutes to room temperature, and hold at 4 °C.

The removal of short oligonucleotides after RNA ID fabrication was performed with Amicon 0.5 mL filters with 100 kDa cut-off using filtered washing buffer (0.5 mM $MgCl_2$, 10 mM Tris-HCl pH 8.0). Synthetic exon mix (40 µL reaction) was filtered with 460 µL washing buffer (460 µL) two times for 10 minutes, 9,200 × g at 3 °C. The sample is collected by reversing the filter after transfer in a new tube and spun down for 2 minutes, 1,000 × g at 3 °C. The concentrations of the synthetic exons are estimated from a NanoDrop spectrophotometer.

Synthetic isoforms fabrication

Synthetic isoforms were assembled by linking synthetic exons. We fabricated four isoforms of which three are order isoforms (same length but different synthetic exon IDs) and one length isoform that has one synthetic exon and extended RNA. The three order isoforms were fabricated using exon I and exon II (RNA isoform ID ‘211312’; Fig. S14A), exon I and exon III (RNA isoform ID '123112'; Fig. S14B), and exon II and exon II (RNA isoform ID '312123'; Fig. S14C). The length isoform is fabricated with exon I and extended RNA (RNA isoform ID '211' extended; Fig. S15).

We mixed 10 µL of each exon (= 20 µL), 2 µL 100 mM $MgCl_2$, 4 µL 1 M NaCl, 14 µL of DNA cuboid (1 µM). The mixtures were incubated at room temperature (20 °C) overnight. After incubation excess DNA cuboid was filtered using afore introduced Amicon 0.5 mL filters (100 kDa cutoff). Buffers are filtered with the MF-Millipore™ Membrane Filter, 0.22 µm pore size.

Circular and linear IDs fabrication

To verify if ARTEMIS can discriminate these two conformations we used circular single-stranded m13mp18 (Guild BioSciences). The linear version was made by annealing a 39 nt oligonucleotide (5' – TCTAGAGGATCCCCGGGTACCGAGCTCGAATTCGTAATC – 3', IDT, IDTE buffer, pH 8.0) to circular form and then subsequent restriction digestion.

Firstly, 40 µL of m13mp18 DNA (100 nM) was mixed with 2 µL oligonucleotide (100µM), 8 µL 10 × Cutsmart buffer (New England Biolabs), and 28 µL of filtered Milli-Q water. This mixture is heated to 65 °C for 30 seconds and gradually cooled down to 25°C over 40 minutes.

After oligonucleotide annealing 1 µL of BamHI-HF (100.000 units/mL, NEB, catalog number R3136T) and 1 µL of EcoRI-HF (100.000 units/mL, NEB, catalog number R3101T) were added, mixed by pipetting, and incubated at 37 °C for 1 hour. The linear form is purified with Macherey-Nagel™ NucleoSpin™ Gel and PCR Clean-up Kit (Macherey-Nagel™, catalog number 740609.50). We mixed by pipetting 400 µL (5 × 40 µL mix) of cut ss m13mp18 with 800 µL of binding buffer and separated to three columns. We followed the manufacturer's manual regarding the washing step and centrifugation conditions. Elution buffer was preheated to 70 °C to improve elution from the column. The elution step was repeated twice with 30 µL of elution buffer, after 5 minutes incubation. The concentration of linear m13mp18 is estimated from a NanoDrop spectrophotometer.

To fabricate circular and linear IDs the same oligonucleotide mixture was used (oligonucleotide listed in Table 2 and Table S12). We prepared 40 µL reaction by mixing linear or circular form (20 nM or 800 fmoles) and oligonucleotides (60 nM each or 2400 fmoles), in 10 mM $MgCl_2$, 1 × TE (10 mM Tris-HCl, 1 mM EDTA, pH 8.0) buffer, and nuclease-free water was added to the final reaction volume. Buffers are filtered with the MF-Millipore™ Membrane Filter, 0.22 µm pore size. The mix is mixed by pipetting and spin down. The mixture was heated up to 70 °C for 30 s and after gradually cooled down (-0.5 °C/cycle, 90 cycles each 30 s) over 45 minutes to room temperature, and hold at 4 °C.

*In vitro* RNA circularization

To create circular RNA we ligated MS2 RNA using T4 RNA ligase 1 and PEG8000 (New England Biolabs (NEB), M0204) that should lead to single-stranded RNA circularization (13). A 20 µL reaction contained 1 × Reaction Buffer (50 mM Tris-HCl, pH 7.5, 10 mM $MgCl_2$, 1 mM DTT), MS2 RNA (150 nM), 1 µL (10 units) T4 RNA Ligase, 10 % PEG8000 and 30 µM ATP. The reaction was incubated overnight at 16 °C.

To create exclusively circular MS2 RNA ID ‘111’/MS2 a complementary oligonucleotide (1.25 µm) that should join MS2 ends was added to the RNA ID fabrication step.

Nanopore fabrication

We fabricated 10-15 nm nanopores using a laser-assisted capillary puller (P2000F, Sutter Instruments). Glass capillaries with an outer diameter of 0.5 mm and an inner diameter of 0.2 mm with filament were purchased from Sutter Instruments, USA. The nanopore diameter was determined with scanning electron microscopy (SEM) and calculated from the conductance of nanopores as previously described (3).

Nanopore measurement and data analysis

All measurements were performed in 4 M LiCl, 1 × TE, pH 9.4 using Axopatch 200B, and data were collected under a constant voltage of 600 mV. Single events in ionic current recordings were firstly isolated according to threshold parameters such as duration, current drop, and event charge deficit (ECD). From isolated events, we can discriminate conformation of nucleic acids and for analysis of linear barcodes, we used only unfolded events. For analysis of circular RNA, we included all data, and since fully folded events were present at a negligible level in control measurements their effect on data interpretation is minimal.

**Figure S1.**

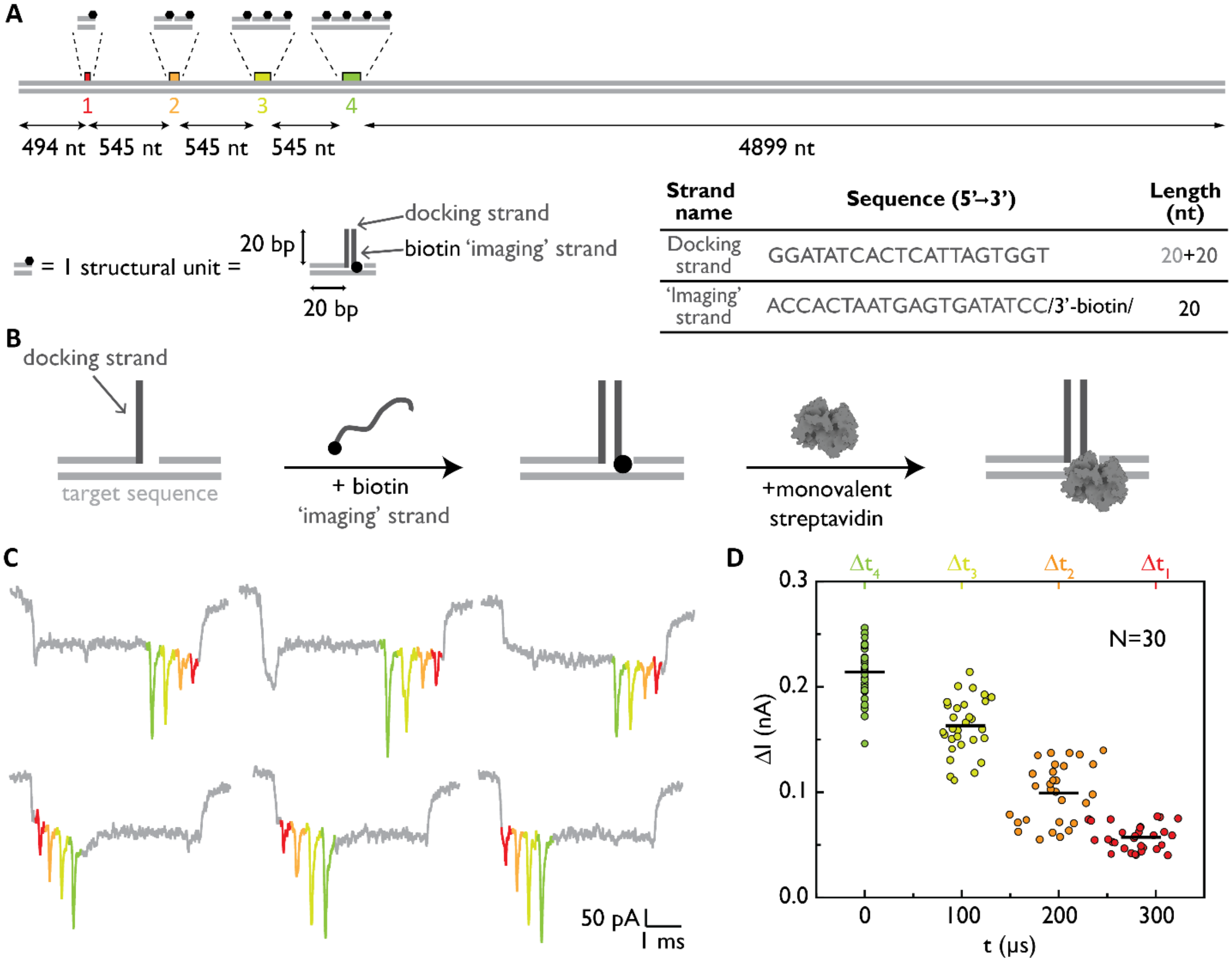


**Figure S1.** Design and analysis of 4-color ruler. (**A**) Design of 4-color ruler indicates four sites that have 1, 2, 3, or 4 structural units each having 20 nt complementary to scaffold strand (**grey**). The structural unit is composed of docking strand overhang (**dark grey**) and imaging strand with 3' biotin (**black**). Sequences of both strands are shown in the table including their length. (**B**) Protocol for the fabrication of a structural unit. In the ID fabrication step, docking and imaging strands form the duplex. Monovalent streptavidin (femtomolar affinity to biotin with inactivated three out of four biotin-binding sites) (14) is added prior to an experiment. (**C**) Example ruler events clearly indicate four downward signals corresponding to structural color from (A) 1 (red), 2 (orange), 3 (lemon yellow), and 4 (green). (**D**) Each detected structural color position is plotted by taking structural color 4 as zero time point and showing the distance from it for structural colors 3, 2, and 1. The current signal for each color is calculated as a drop from the first current drop level originating from the ruler itself. The sample size is thirty unfolded ruler events.

**Fig. S2.**

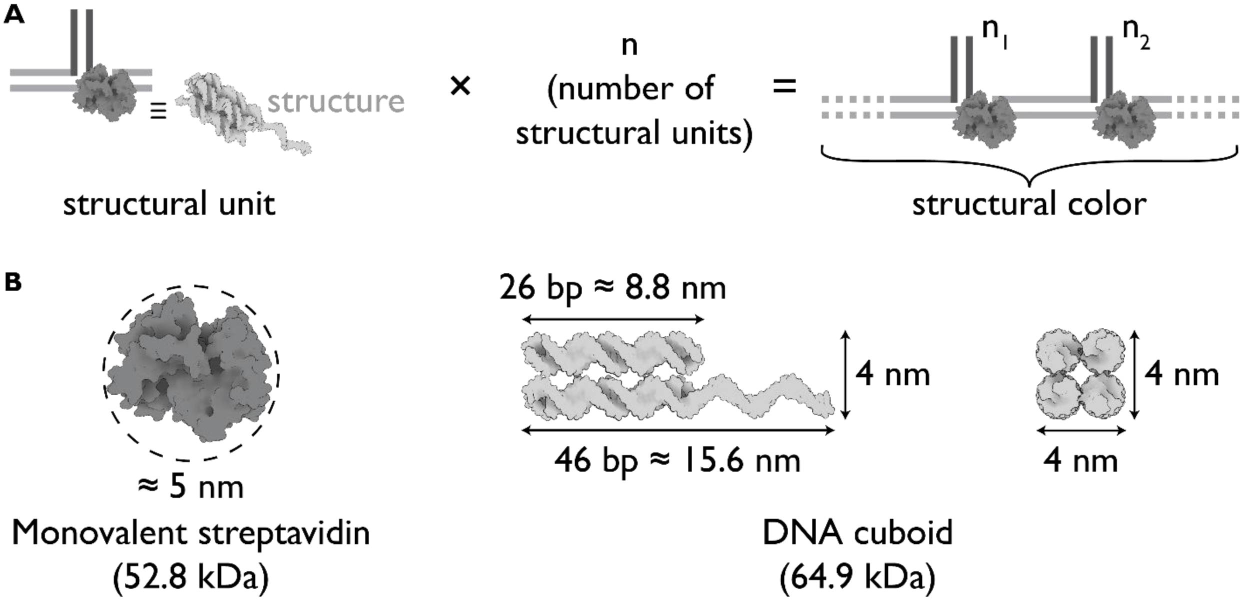


**Figure S2.** Design of the structural unit and structural color. (**A**) The key element of the structural unit is a structure required to cause a detectable signal for a nanopore microscope. Here, we demonstrated the use of protein structure (monovalent streptavidin – dark grey) and DNA-only system (DNA cuboid – light grey). Each structural color is composed of interspaced structural units, where a number of structural units correspond to the structural color. For instance, two structural units would lead to structural color '2' i.e. specific drop in the ionic current. (**B**) Physical characteristics of both monovalent streptavidin (52.8 kDa) and DNA cuboid (64.9 kDa) are delineated. Monovalent streptavidin has a diameter of 5-6 nm. DNA cuboid has the length of 15.6 nm (46 bp) with imaging strand and 8.8 nm (26 bp) without, while the width corresponds to two DNA helixes or 4 nm.

**Figure S3.**

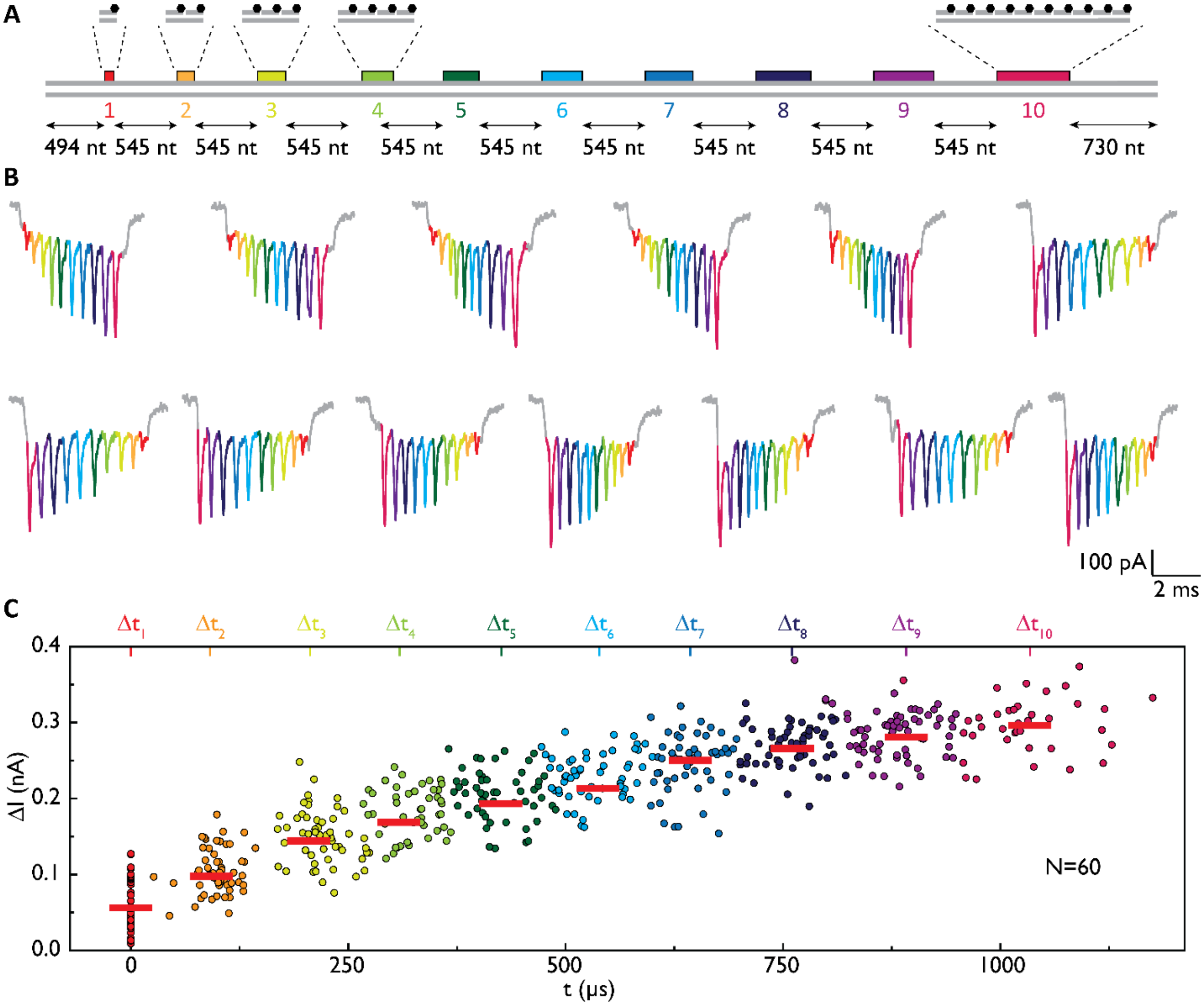


**Figure S3.** Design and additional example events of the 10-color ruler. (**A**) Design of 10-color ruler indicates ten sites that have 1, 2, 3, 4, 5, 6, 7, 8, 9, and 10 structural units. (**B**) Example ruler events indicate ten downward signals corresponding to structural colour from (A) 1 (red), 2 (orange), 3 (lemon yellow), 4 (green), 5 (dark green), 6 (light blue), 7 (sky blue), 8 (dark blue), 9 (purple) and 10 (burgundy red). (**C**) Scatter plot of current drop and normalized position for each structural color. The correct structural colors are identified as a distinctive signal on a single-event basis. The sample size is sixty unfolded ruler events.

**Figure S4.**

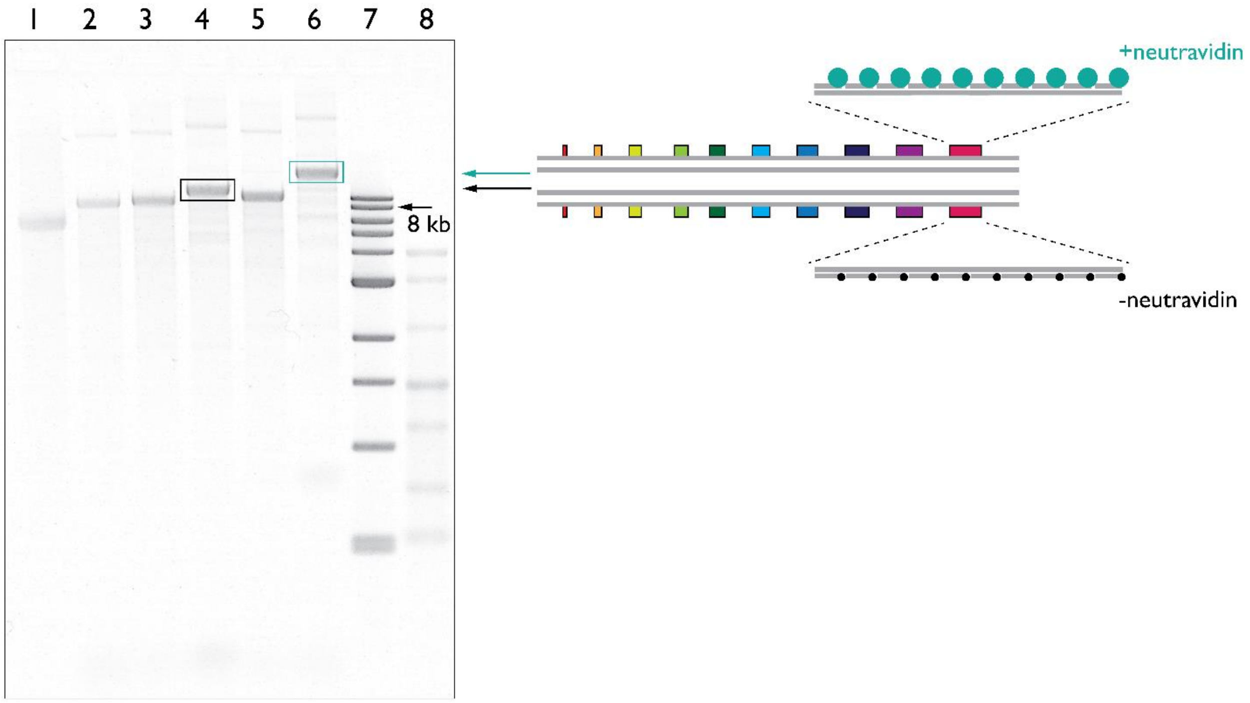


**Figure S4.** Electrophoresis analysis of fabricated 4-color and 10-color rulers on the agarose gel. Lanes: 1-linear single-stranded M13, 2 – double-stranded M13, 3 – 4-color ruler ***without neutravidin***, 4 – 10-color ruler ***without neutravidin***, 5 – 4-color ruler with neutravidin, 6 – 10-color ruler with neutravidin, 7 – 1 kb DNA ladder (NEB), 8 – single-stranded RNA ladder (NEB). Gel: 1% (w/v) agarose, 1 × TBE.

**Figure S5.**

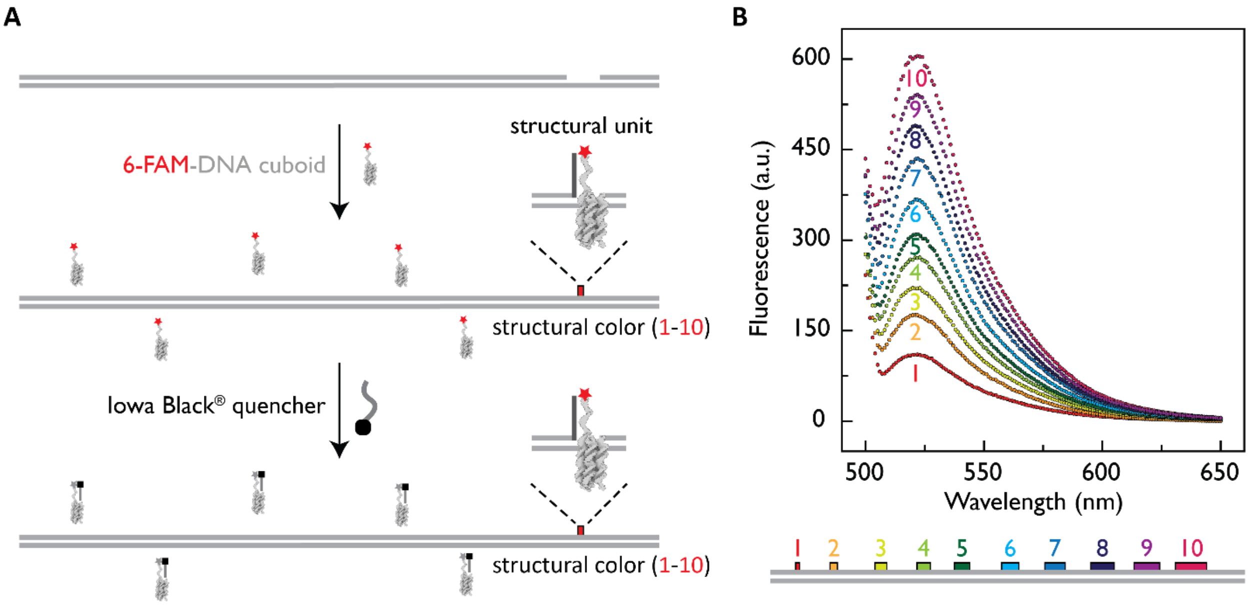


**Figure S5.** Fluorescence quenching assay for validation of expected structural color. (**A**) To a double-stranded molecular ruler, we added a specific number of structural units (1 or 2 … or 10) with 5'-fluorescein (6-FAM). The excess of 6-FAM DNA cuboid strands is quenched by binding of Iowa Black quencher. After the quenching, only 6-FAM DNA cuboids in the structural unit would emit a fluorescence signal. (**B**) Using the equimolar concentration of molecular IDs we measured fluorescence after background quenching indicating corresponding example measurement for each of 10 structural colors separately.

**Fig. S6.**

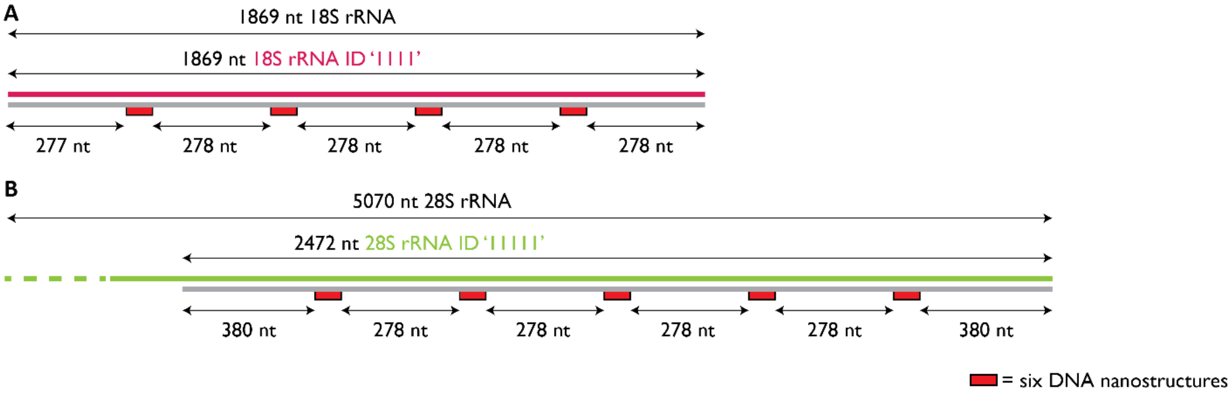


**Figure S6.** RNA ID designs for 18S rRNA and 28S rRNA. (**A**) 18S rRNA ID '1111' design. (**B**) 28S rRNA ID '11111' design.

**Figure S7.**

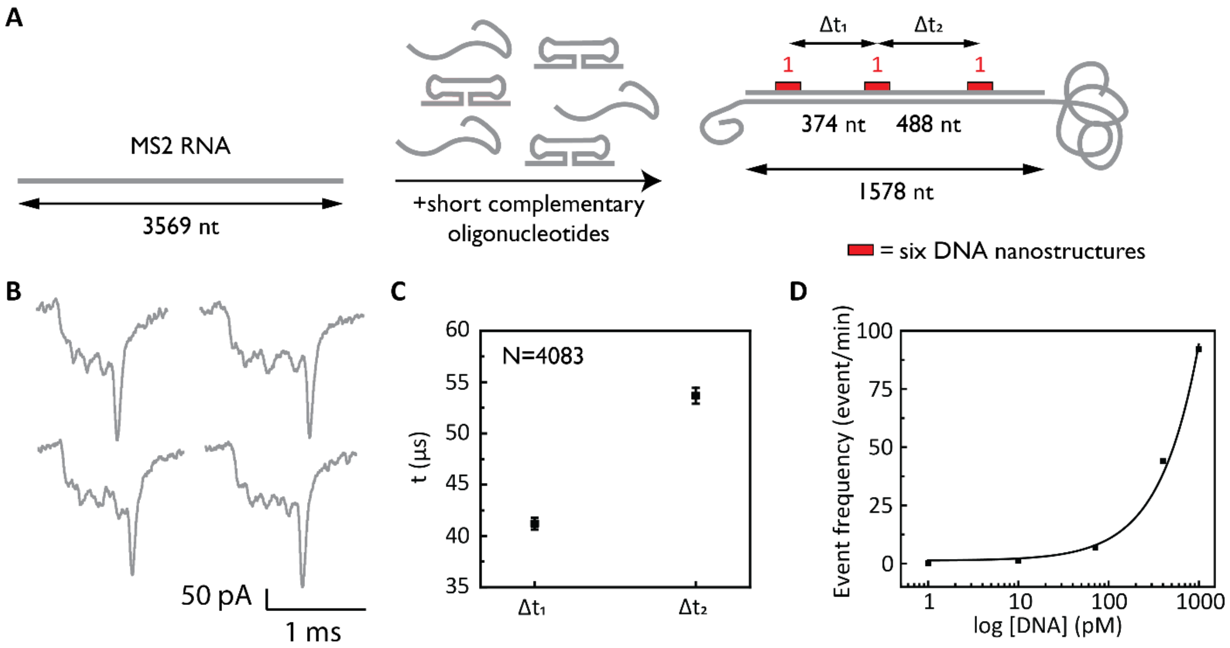


**Figure S7.** Fabrication of RNA ID using a part of long RNA ~3.6 kb. (**A**) ID '111' is designed to be in the middle of RNA using part of RNA for ID fabrication. (**B**) Detected IDs from nanopore recordings have three visible downward signals and the deep drop originating from a single-stranded coil outside of the ID region. (**C**) The time difference between the first two (374 nt) and the last two (488 nt) structural colors. (**D**) Concentration dependence of capture rate i.e. translocation frequency. The sample size is 4083 events.

**Fig. S8.**

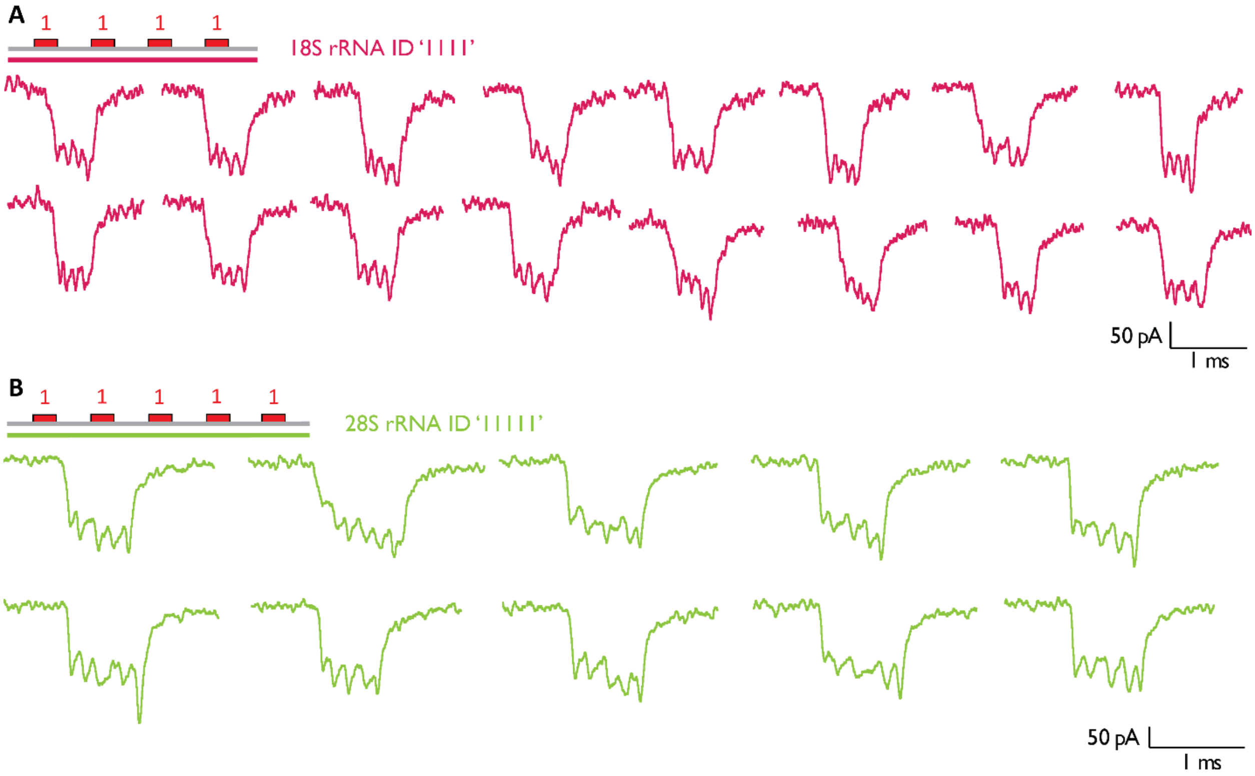


**Figure S8.** RNA ID additional event examples for 18S rRNA and 28S rRNA IDs. (**A**) 18S rRNA ID '1111' examples. (**B**) 28S rRNA ID '11111' examples.

**Fig. S9.**

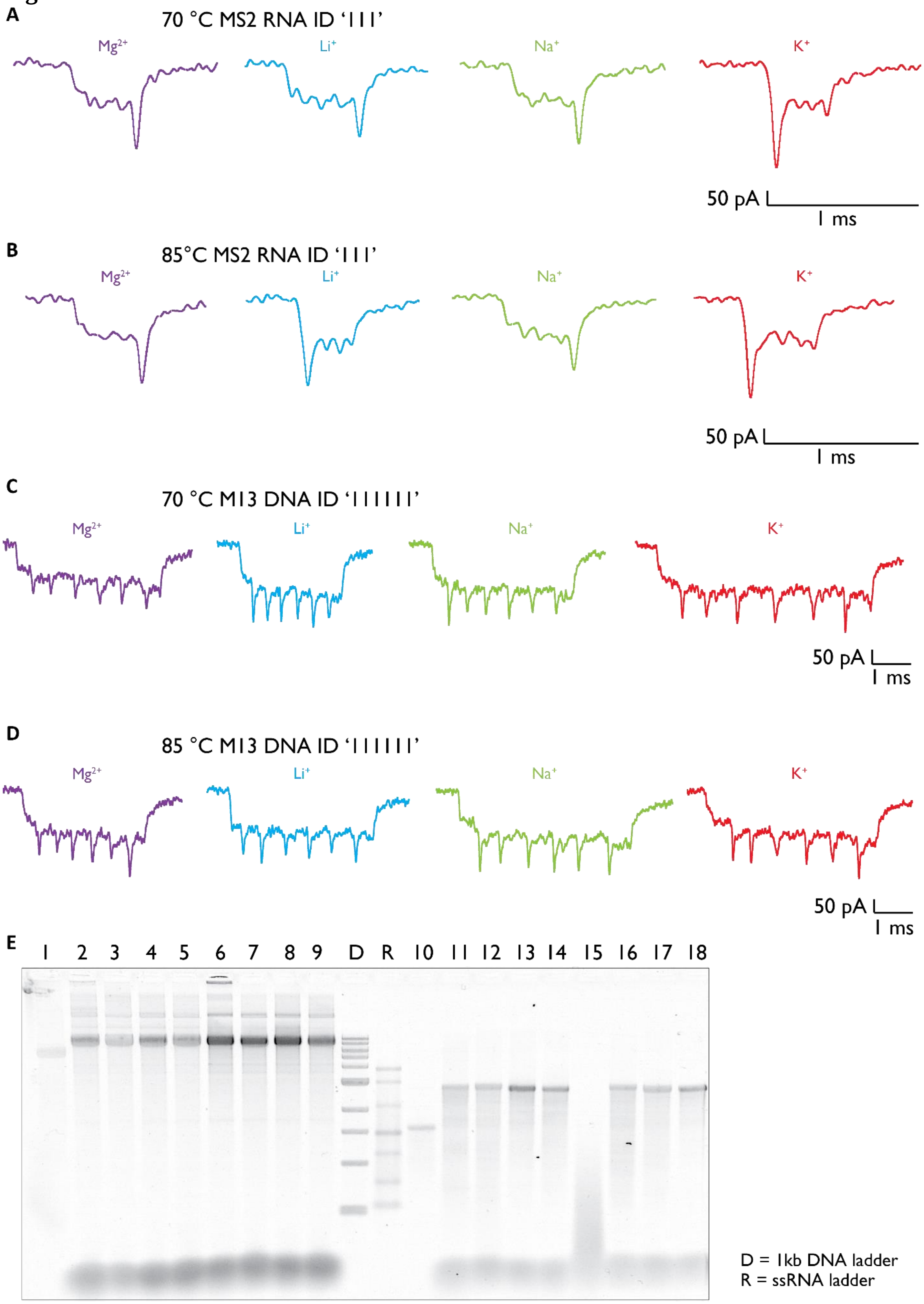
A
70 °C MS2 RNA ID '111'
Mg2+
Li+
Na+
K+
50 pA
1 ms
B
85°C MS2 RNA ID '111'
Mg2+
Li+
Na+
K+
50 pA
1 ms
C
70 °C M13 DNA ID '111111'
Mg2+
Li+
Na+
K+
50 pA
1 ms
D
85 °C M13 DNA ID '111111'
Mg2+
Li+
Na+
K+
50 pA
1 ms
E
1 2 3 4 5 6 7 8 9 D R 10 11 12 13 14 15 16 17 18
D = 1kb DNA ladder
R = ssRNA ladder

**Figure S9.** Effects of temperature and salts on ID fabrication. (**A**) Example events for MS2 RNA ID '111' using 70 °C for fabrication. (**B**) Example events for MS2 RNA ID '111' using 85 °C for fabrication. (**C**) Example events for M13 DNA ID '111111' using 70 °C for fabrication. (**D**) Example for M13 DNA ID '111111' using 85 °C for fabrication. (**E**) Agarose gel indicates ID fabrication over various conditions. 1 – ssM13 DNA, 2-4 – samples from (C), 6-9 – samples from (D), D – 1kb DNA ladder, R – ssRNA ladder, 10 – ssMS2 RNA, 11-14 – samples from (A), 15-18 – samples from (B). Gel: 1% (w/v) agarose, 1 × TBE.

**Fig. S10.**

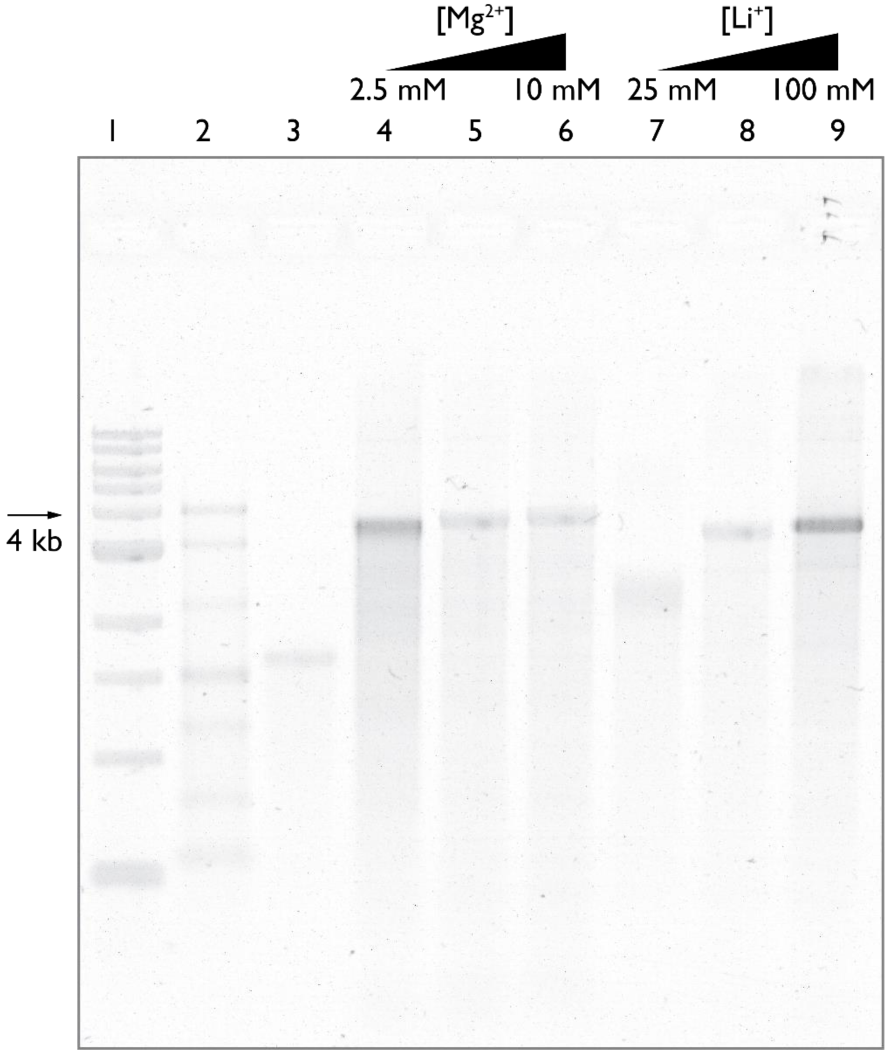


**Figure S10.** Effects of salt concentration on ID fabrication. Agarose gel indicates ID fabrication for magnesium and lithium chloride for the three concentrations. 1 – 1kb DNA ladder; 2 – ssRNA ladder; 3 – ssMS2 RNA; 4, 5, 6 – MS2 RNA ID '111' (fully complementary) for 2.5, 5, and 10 mM $MgCl_2$; 7, 8, and 9 – MS2 RNA ID '111' (fully complementary) for 25, 50, and 100 mM LiCl. Gel: 1% (w/v) agarose, 1 × TBE.

**Fig. S11.**

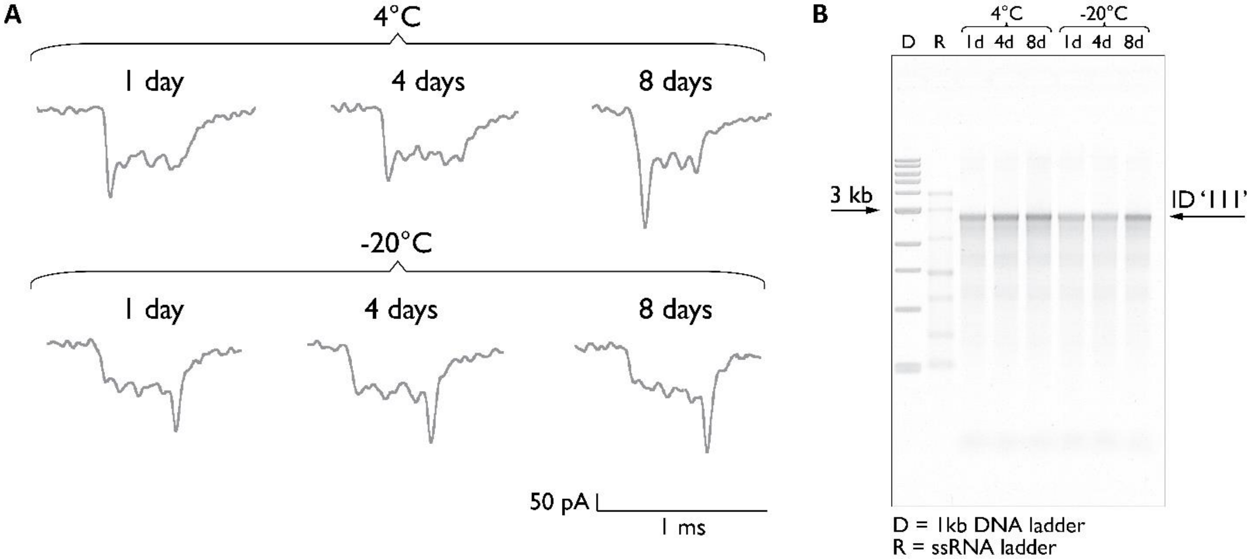


**Figure S11.** Stability of RNA IDs over time under different storage temperatures. (**A**) Example events indicate correct ID readout over 8 days for 4 °C and -20 °C. (**B**) RNA IDs have not shown significant difference over time for three times points (1 day, 4 days, and 8 days) at both 4 °C and -20 °C. Gel: 1% (w/v) agarose, 1 × TBE.

**Fig. S12.**

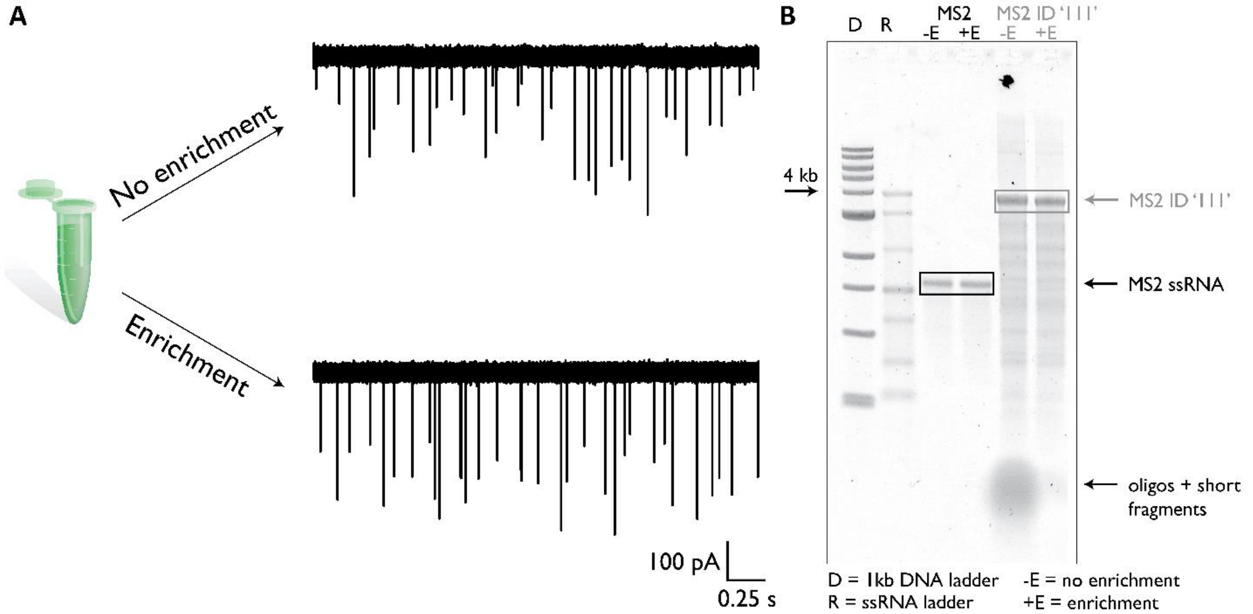


**Figure S12.** Enrichment of target RNA ID from a background of short nucleic acid fragments (<100 kDa). (**A**) Cumulative events for MS2 ID '111'p after RNA ID fabrication without and with enrichment. The ionic current trace after enrichment indicates the removal of short nucleic acid background. (**B**) Agarose gel indicates successful removal of oligos and short RNAs after enrichment. Gel: 1% (w/v) agarose, 1 × TBE.

**Fig. S13.**

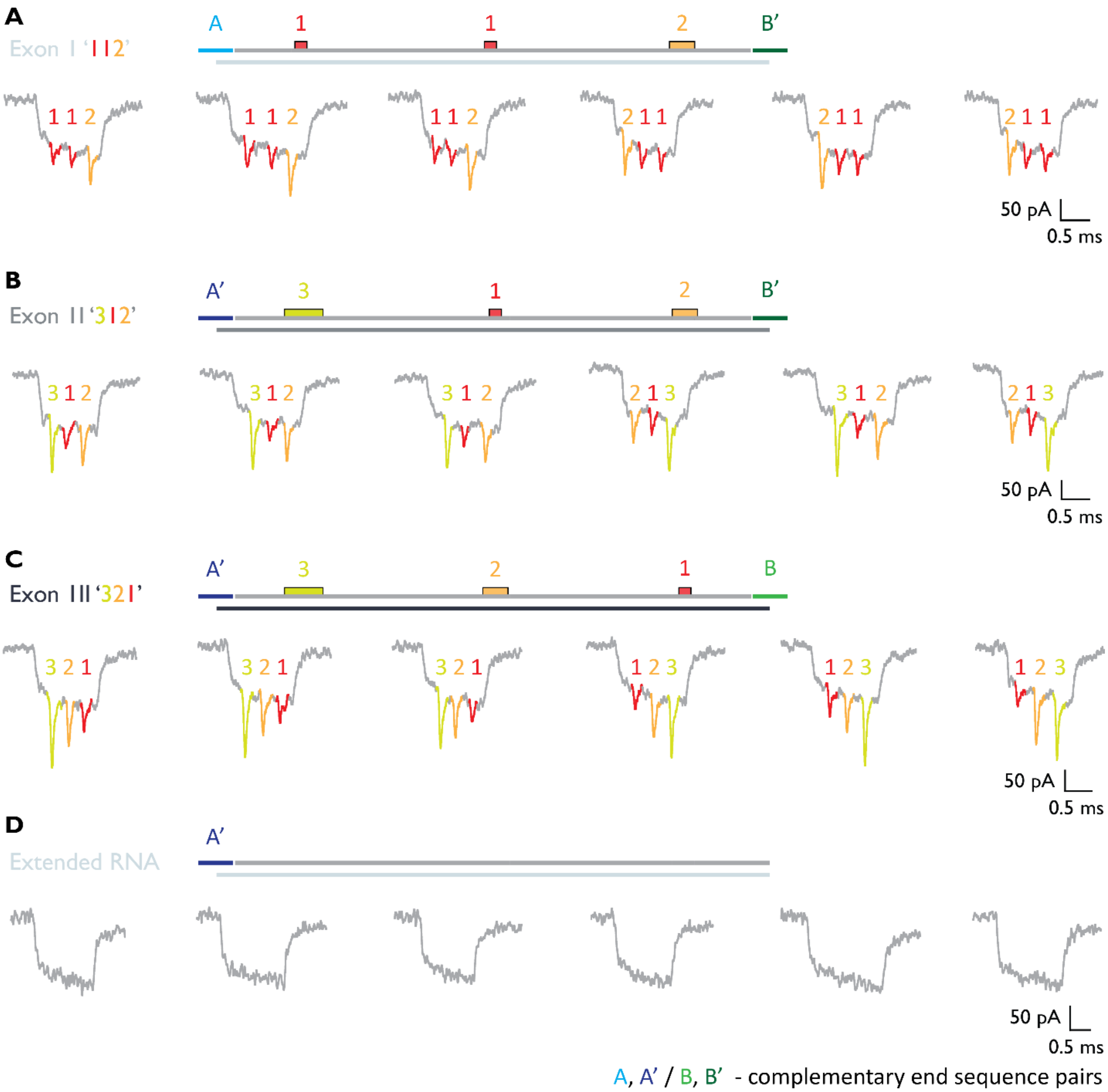


**Figure S13.** Design of exons with example nanopore events. (**A**) Design of exon I ID '112' with terminal overhangs A and B'. (**B**) Design of exon II ID '312' with terminal overhangs A' and B'. (**C**) Design of exon III ID '321' with terminal overhangs A' and B. (**D**) Design of extended RNA ID with terminal overhangs A'.

**Fig. S14.**

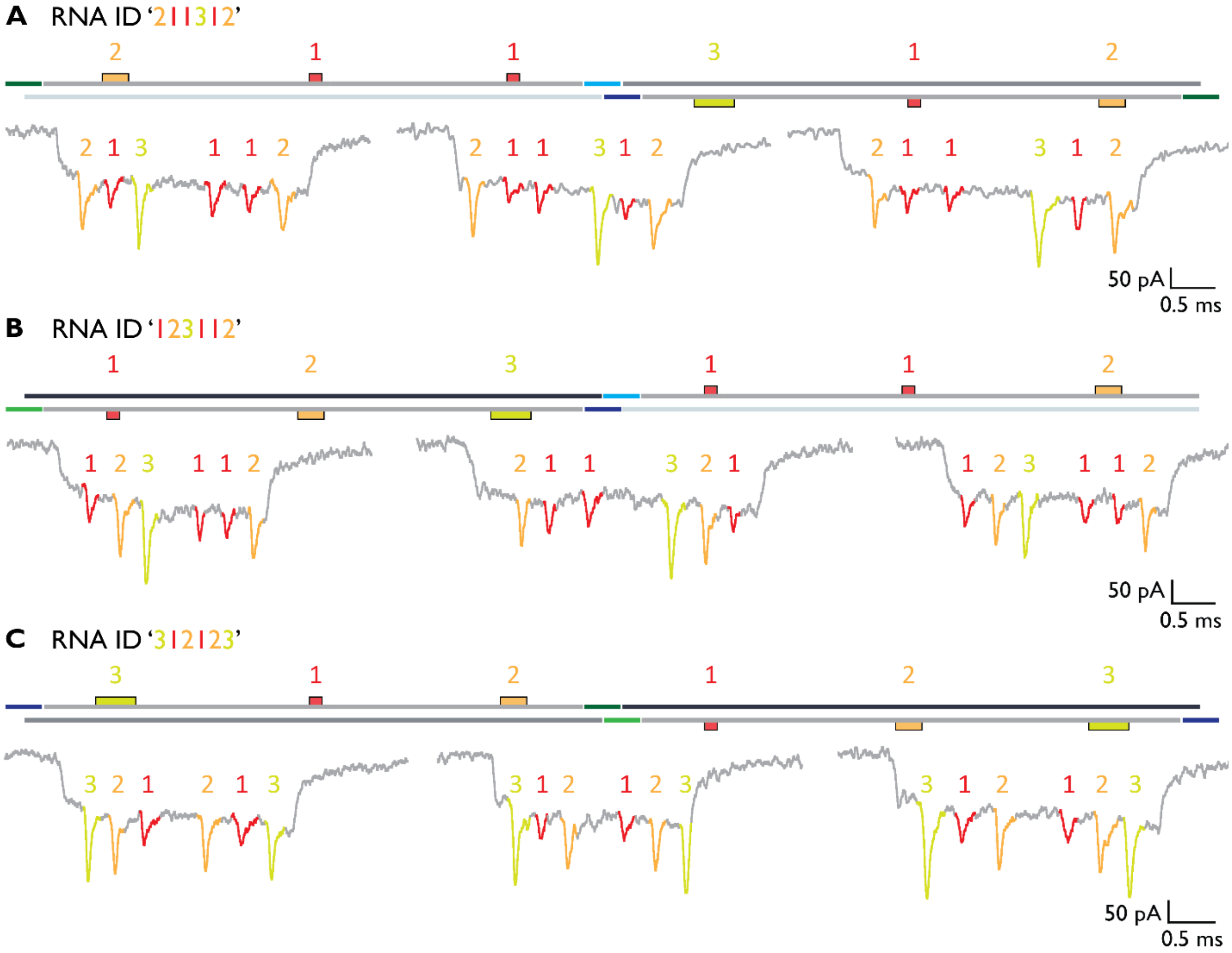


**Figure S14.** Design and example nanopore events for order isoforms. (**A**) Design of RNA ID '211312'. (**B**) Design of RNA ID '123112'. (**C**) Design of RNA ID '312123'

**Fig. S15.**

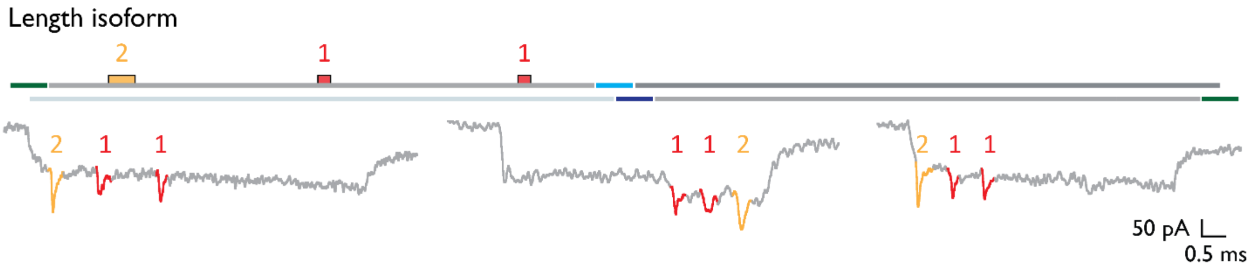


**Figure S15.** Length isoform ‘211’ with extended RNA.

**Fig. S16.**

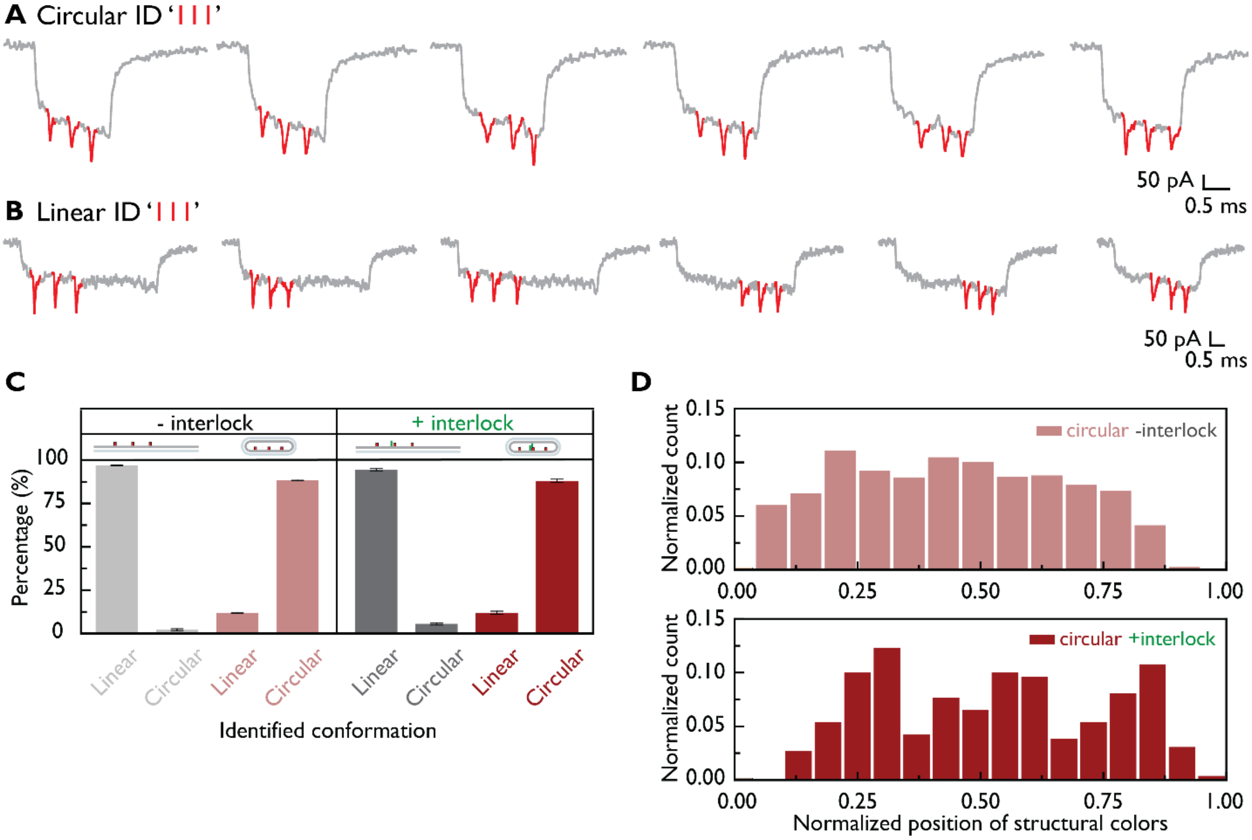


**Figure S16.** Circular and linear isoforms. (**A**) Example events for circular ID ‘111’. (**B**) Example events for linear ID ‘111’. (**C**) The percentage of identified conformation with and without the oligo interlock. As expected in linearized single-stranded conformation (with and without interlock) almost all events are identified as linear conformation. In the case of circular conformation majority of events were identified as circular conformation.

**Fig. S17.**

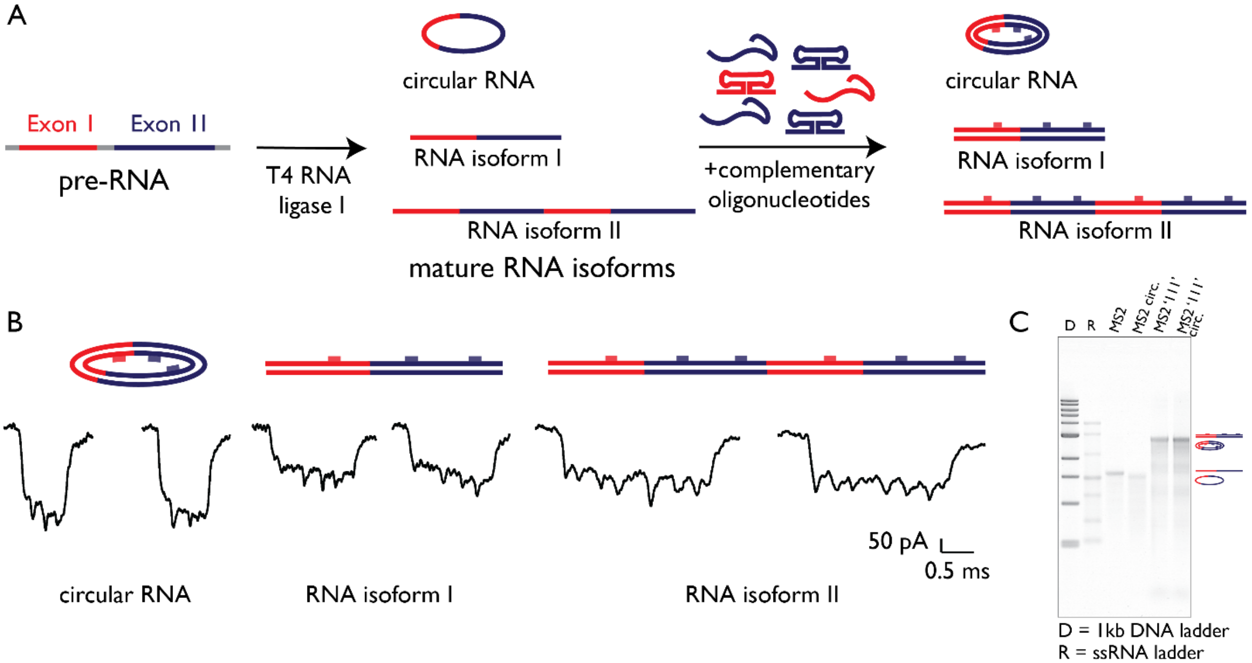


**Figure S17.** Mimicking of trans-splicing and backsplicing with T4 RNA ligase 1. (**A**) Experimental design of 'alternative splicing' with T4 RNA ligase 1. Circularization assay should promote intramolecular ligation i.e. circularization with potential intermolecular ligation. (**B**) Example events identified after ligation with nanopore measurements. circRNA conformation was fixed with two weak oligo linkers and successfully identified. Original RNA with unique RNA ID '111' was detected as well as its trans-spliced variant.

**Fig. S18.**

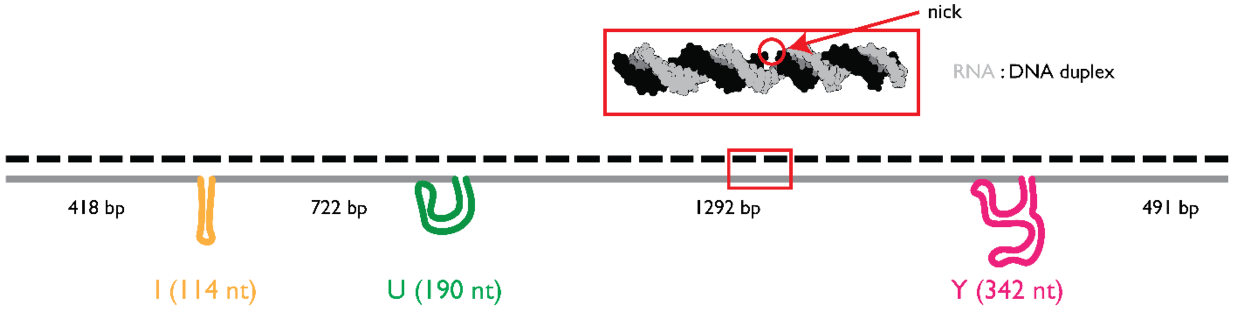


**Figure S18.** Design of internal RNA origami ID ‘IUY’ with distances between structural colors. RNA : DNA duplex structure (red rectangle) and one of nicks are circled in red.

**Fig. S19.**

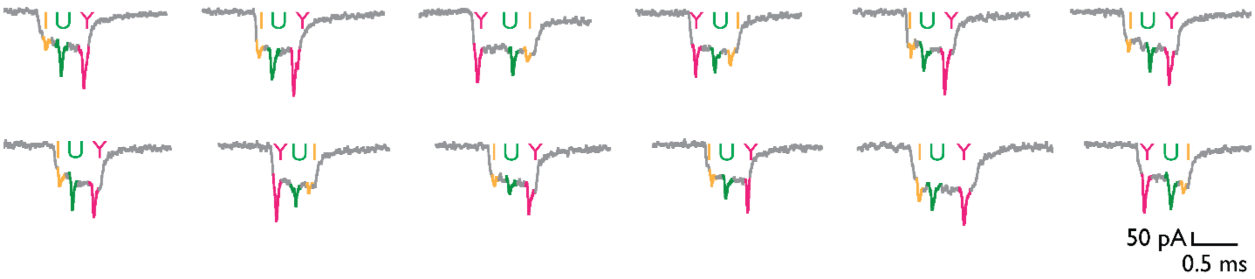


**Figure S19.** Example events for internal RNA origami ID ‘IUY’.

**Fig. S20.**

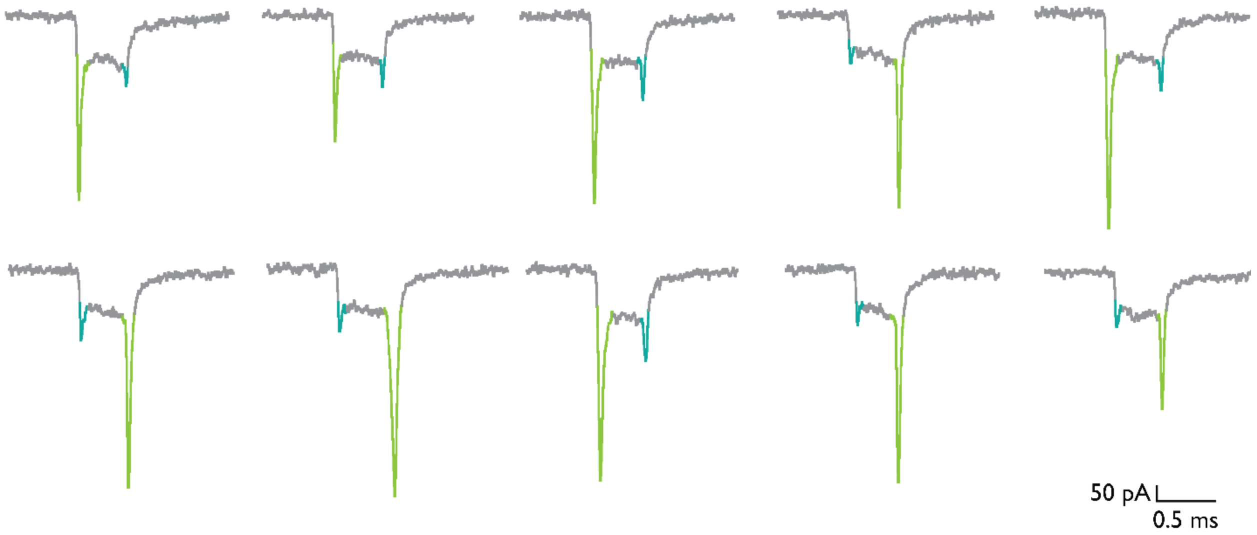


**Figure S20.** Example events for terminal RNA origami ID ‘AΩ’.

**Fig. S21.**

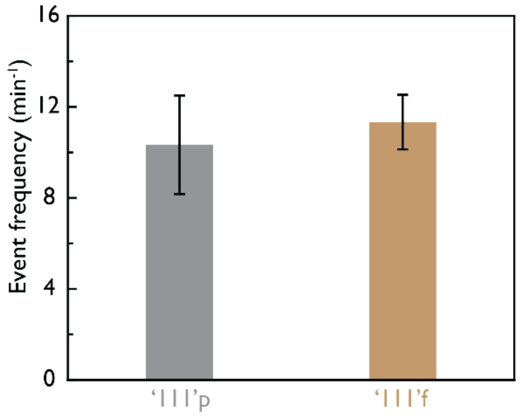


**Figure S21.** Event frequency from partially and fully complemented MS2 RNA ID '111'. Translocation frequency for three individual measurements in equimolar concentrations of both '111'p and '111'f is plotted. The event number for all three individual measurements was 6566 events. Error bars are shown as ± standard error.

**Fig. S22.**

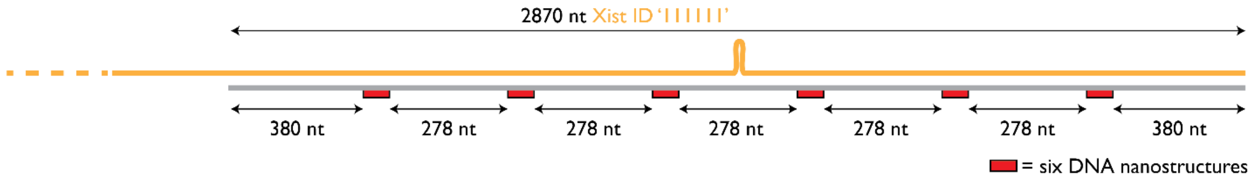


**Figure S22.** *Xist* lncRNA ID design.

**Table S1.**

**Table S1.** DNA oligonucleotides for DNA cuboid. In red is highlighted 'imaging strand' complementary to the structural unit 'docking strand'. In purple is labeled oligo with 6-fluorescein (6-FAM) used for the fluorescence-quenching assay instead of 1M1.

| Name | Sequence (5'→3') | Length (nt) |
|---|---|---|
| 2S1 | GCCGACGTGTACGGATCTGGCA | 22 |
| 2S2 | GCGACATTGCGGCCGATTCGGA | 22 |
| 1M1 | ACCACTAATGAGTGATATCCTTTTCGCATCAGGGCACATTGGCTTT | 46 |
| 1M2 | TTTTGCGGCTAAGACTTGCCAGGTTT | 26 |
| 3L1 | TTTTACACGTCGGCCCTGATGCGACCTGGCAAGTTCCGAATCGGCTTT | 48 |
| 3L2 | TTTCGCAATGTCGCCTTAGCCGCAGCCAATGTGCTGCCAGATCCGTTT | 48 |
| 1M1_FAM | /56-FAM-/ ACCACTAATGAGTGATATCCTTTTCGCATCAGGGCACATTGGCTTT | 46 |
| 1M1c_Black Iowa Quencher | GGATATCACTCATTAGTGGT/3IABkFQ/ | 20 |

**Table S2.**

**Table S2.** The core DNA oligonucleotides are complementary to linearized ssM13 (7228 nt).

| Oligonucleotide Number | Sequence (5’→’3’) | Oligonucleotide Number | Sequence (5’→’3’) |
|---|---|---|---|
| 1 | TTTTCGTAATCATGGTCATAGCTGTTTCCTGTGTGAAATTGTTATC | 96 | CTTGAGCCATTTGGGAATTAGAGCCAGCAAAATCACCA |
| 2 | CGCTCACAATTCCACACAACATACGAGCCGGAAGCATA | 97 | GTAGCACCATTACCATTAGCAAGGCCGGAAACGTCACC |
| 3 | AAGTGTAAAGCCTGGGGTGCCTAATGAGTGAGCTAACT | 98 | AATGAAACCATCGATAGCAGCACCGTAATCAGTAGCGA |
| 4 | CACATTAATTGCGTTGCGCTCACTGCCCGCTTTCCAGT | 99 | CAGAATCAAGTTTGCCTTTAGCGTCAGACTGTAGCGCG |
| 5 | CGGGAAACCTGTCGTGCCAGCTGCATTAATGAATCGGC | 100 | TTTTCATCGGCATTTTCGGTCATAGCCCCCTTATTAGC |
| 6 | CAACGCGCGGGGAGAGGCGGTTTGCGTATTGGGCGCCA | 101 | GTTTGCCATCTTTTCATAATCAAAATCACCGGAACCAG |
| 7 | GGGTGGTTTTTCTTTTCACCAGTGAGACGGGCAACAGC | 102 | AGCCACCACCGGAACCGCCTCCCTCAGAGCCGCCACCC |
| 8 | TGATTGCCCTTCACCGCCTGGCCCTGAGAGAGTTGCAG | 103 | TCAGAACCGCCACCCTCAGAGCCACCACCCTCAGAGCC |
| 9 | CAAGCGGTCCACGCTGGTTTGCCCCAGCAGGCGAAAAT | 104 | GCCACCAGAACCACCACCAGAGCCGCCGCCAGCATTGA |
| 10 | CCTGTTTGATGGTGGTTCCGAAATCGGCAAAATCCCTT | 105 | CAGGAGGTTGAGGCAGGTCAGACGATTGGCCTTGATAT |
| 11 | ATAAATCAAAAGAATAGCCCGAGATAGGGTTGAGTGTT | 106 | TCACAAACAAATAAATCCTCATTAAAGCCAGAATGGAA |
| 12 | GTTCCAGTTTGGAACAAGAGTCCACTATTAAAGAACGT | 107 | AGCGCAGTCTCTGAATTTACCGTTCCAGTAAGCGTCAT |
| 13 | GGACTCCAACGTCAAAGGGCGAAAAACCGTCTATCAGG | 108 | ACATGGCTTTTGATGATACAGGAGTGTACTGGTAATAA |
| 14 | GCGATGGCCCACTACGTGAACCATCACCCAAATCAAGT | 109 | GTTTTAACGGGGTCAGTGCCTTGAGTAACAGTGCCCGT |
| 15 | TTTTTGGGGTCGAGGTGCCGTAAAGCACTAAATCGGAA | 110 | ATAAACAGTTAATGCCCCCTGCCTATTTCGGAACCTAT |
| 16 | CCCTAAAGGGAGCCCCCGATTTAGAGCTTGACGGGGAA | 111 | TATTCTGAAACATGAAAGTATTAAGAGGCTGAGACTCC |
| 17 | AGCCGGCGAACGTGGCGAGAAAGGAAGGGAAGAAAGCG | 112 | TCAAGAGAAGGATTAGGATTAGCGGGGTTTTGCTCAGT |
| 18 | AAAGGAGCGGGCGCTAGGGCGCTGGCAAGTGTAGCGGT | 113 | ACCAGGCGGATAAGTGCCGTCGAGAGGGTTGATATAAG |
| 19 | CACGCTGCGCGTAACCACCACACCCGCCGCGCTTAATG | 114 | TATAGCCCGGAATAGGTGTATCACCGTACTCAGGAGGT |
| 20 | CGCCGCTACAGGGCGCGTACTATGGTTGCTTTGACGAG | 115 | TTAGTACCGCCACCCTCAGAACCGCCACCCTCAGAACC |
| 21 | CACGTATAACGTGCTTTCCTCGTTAGAATCAGAGCGGG | 116 | GCCACCCTCAGAGCCACCACCCTCATTTTCAGGGATAG |
| 22 | AGCTAAACAGGAGGCCGATTAAAGGGATTTTAGACAGG | 117 | CAAGCCCAATAGGAACCCATGTACCGTAACACTGAGTT |
| 23 | AACGGTACGCCAGAATCCTGAGAAGTGTTTTTATAATC | 118 | TCGTCACCAGTACAAACTACAACGCCTGTAGCATTCCA |
| 24 | AGTGAGGCCACCGAGTAAAAGAGTCTGTCCATCACGCA | 119 | CAGACAGCCCTCATAGTTAGCGTAACGATCTAAAGTTT |
| 25 | AATTAACCGTTGTAGCAATACTTCTTTGATTAGTAATA | 120 | TGTCGTCTTTCCAGACGTTAGTAAATGAATTTTCTGTA |
| 26 | ACATCACTTGCCTGAGTAGAAGAACTCAAACTATCGGC | 121 | TGGGATTTTGCTAAACAACTTTCAACAGTTTCAGCGGA |
| 27 | CTTGCTGGTAATATCCAGAACAATATTACCGCCAGCCA | 122 | GTGAGAATAGAAAGGAACAACTAAAGGAATTGCGAATA |
| 28 | TTGCAACAGGAAAAACGCTCATGGAAATACCTACATTT | 123 | ATAATTTTTTCACGTTGAAAATCTCCAAAAAAAAGGCT |
| 29 | TGACGCTCAATCGTCTGAAATGGATTATTTACATTGGC | 124 | CCAAAAGGAGCCTTTAATTGTATCGGTTTATCAGCTTG |
| 30 | AGATTCACCAGTCACACGACCAGTAATAAAAGGGACAT | 125 | CTTTCGAGGTGAATTTCTTAAACAGCTTGATACCGATA |
| 31 | TCTGGCCAACAGAGATAGAACCCTTCTGACCTGAAAGC | 126 | GTTGCGCCGACAATGACAACAACCATCGCCCACGCATA |
| 32 | GTAAGAATACGTGGCACAGACAATATTTTTGAATGGCT | 127 | ACCGATATATTCGGTCGCTGAGGCTTGCAGGGAGTTAA |
| 33 | ATTAGTCTTTAATGCGCGAACTGATAGCCCTAAAACAT | 128 | AGGCCGCTTTTGCGGGATCGTCACCCTCAGCAGCGAAA |
| 34 | CGCCATTAAAAATACCGAACGAACCACCAGCAGAAGAT | 129 | GACAGCATCGGAACGAGGGTAGCAACGGCTACAGAGGC |
| 35 | AAAACAGAGGTGAGGCGGTCAGTATTAACACCGCCTGC | 130 | TTTGAGGACTAAAGACTTTTTCATGAGGAAGTTTCCAT |
| 36 | AACAGTGCCACGCTGAGAGCCAGCAGCAAATGAAAAAT | 131 | TAAACGGGTAAAATACGTAATGCCACTACGAAGGCACC |
| 37 | CTAAAGCATCACCTTGCTGAACCTCAAATATCAAACCC | 132 | AACCTAAAACGAAAGAGGCAAAAGAATACACTAAAACA |
| 38 | TCAATCAATATCTGGTCAGTTGGCAAATCAACAGTTGA | 133 | CTCATCTTTGACCCCCAGCGATTATACCAAGCGCGAAA |

| | | | |
|---|---|---|---|
| 39 | AAGGAATTGAGGAAGGTTATCTAAAATATCTTTAGGAG | 134 | CAAAGTACAACGGAGATTTGTATCATCGCCTGATAAAT |
| 40 | CACTAACAACTAATAGATTAGAGCCGTCAATAGATAAT | 135 | TGTGTCGAAATCCGCGACCTGCTCCATGTTACTTAGCC |
| 41 | ACATTTGAGGATTTAGAAGTATTAGACTTTACAAACAA | 136 | GGAACGAGGCGCAGACGGTCAATCATAAGGGAACCGAA |
| 42 | TTCGACAACTCGTATTAAATCCTTTGCCCGAACGTTAT | 137 | CTGACCAACTTTGAAAGAGGACAGATGAACGGTGTACA |
| 43 | TAATTTTAAAAGTTTGAGTAACATTATCATTTTGCGGA | 138 | GACCAGGCGCATAGGCTGGCTGACCTTCATCAAGAGTA |
| 44 | ACAAAGAAACCACCAGAAGGAGCGGAATTATCATCATA | 139 | ATCTTGACAAGAACCGGATATTCATTACCCAAATCAAC |
| 45 | TTCCTGATTATCAGATGATGGCAATTCATCAATATAAT | 140 | GTAACAAAGCTGCTCATTCAGTGAATAAGGCTTGCCCT |
| 46 | CCTGATTGTTTGGATTATACTTCTGAATAATGGAAGGG | 141 | GACGAGAAACACCAGAACGAGTAGTAAATTGGGCTTGA |
| 47 | TTAGAACCTACCATATCAAAATTATTTGCACGTAAAAC | 142 | GATGGTTTAATTTCAACTTTAATCATTGTGAATTACCT |
| 48 | AGAAATAAAGAAATTGCGTAGATTTTCAGGTTTAACGT | 143 | TATGCGATTTTAAGAACTGGCTCATTATACCAGTCAGG |
| 49 | CAGATGAATATACAGTAACAGTACCTTTTACATCGGGA | 144 | ACGTTGGGAAGAAAAATCTACGTTAATAAAACGAACTA |
| 50 | GAAACAATAACGGATTCGCCTGATTGCTTTGAATACCA | 145 | ACGGAACAACATTATTACAGGTAGAAAGATTCATCAGT |
| 51 | AGTTACAAAATCGCGCAGAGGCGAATTATTCATTTCAA | 146 | TGAGATTTAGGAATACCACATTCAACTAATGCAGATAC |
| 52 | TTACCTGAGCAAAAGAAGATGATGAAACAAACATCAAG | 147 | ATAACGCCAAAAGGAATTACGAGGCATAGTAAGAGCAA |
| 53 | AAAACAAAATTAATTACATTTAACAATTTCATTTGAAT | 148 | CACTATCATAACCCTCGTTTACCAGACGACGATAAAAA |
| 54 | TACCTTTTTTAATGGAAACAGTACATAAATCAATATAT | 149 | CCAAAATAGCGAGAGGCTTTTGCAAAAGAAGTTTTGCC |
| 55 | GTGAGTGAATAACCTTGCTTCTGTAAATCGTCGCTATT | 150 | AGAGGGGGTAATAGTAAAATGTTTAGACTGGATAGCGT |
| 56 | AATTAATTTTCCCTTAGAATCCTTGAAAACATAGCGAT | 151 | CCAATACTGCGGAATCGTCATAAATATTCATTGAATCC |
| 57 | AGCTTAGATTAAGACGCTGAGAAGAGTCAATAGTGAAT | 152 | CCCTCAAATGCTTTAAACAGTTCAGAAAACGAGAATGA |
| 58 | TTATCAAAATCATAGGTCTGAGAGACTACCTTTTTAAC | 153 | CCATAAATCAAAAATCAGGTCTTTACCCTGACTATTAT |
| 59 | CTCCGGCTTAGGTTGGGTTATATAACTATATGTAAATG | 154 | AGTCAGAAGCAAAGCGGATTGCATCAAAAAGATTAAGA |
| 60 | CTGATGCAAATCCAATCGCAAGACAAAGAACGCGAGAA | 155 | GGAAGCCCGAAAGACTTCAAATATCGCGTTTTAATTCG |
| 61 | AACTTTTTCAAATATATTTTAGTTAATTTCATCTTCTG | 156 | AGCTTCAAAGCGAACCAGACCGGAAGCAAACTCCAACA |
| 62 | ACCTAAATTTAATGGTTTGAAATACCGACCGTGTGATA | 157 | GGTCAGGATTAGAGAGTACCTTTAATTGCTCCTTTTGA |
| 63 | AATAAGGCGTTAAATAAGAATAAACACCGGAATCATAA | 158 | TAAGAGGTCATTTTTGCGGATGGCTTAGAGCTTAATTG |
| 64 | TTACTAGAAAAAGCCTGTTTAGTATCATATGCGTTATA | 159 | CTGAATATAATGCTGTAGCTCAACATGTTTTAAATATG |
| 65 | CAAATTCTTACCAGTATAAAGCCAACGCTCAACAGTAG | 160 | CAACTAAAGTACGGTGTCTGGAAGTTTCATTCCATATA |
| 66 | GGCTTAATTGAGAATCGCCATATTTAACAACGCCAACA | 161 | ACAGTTGATTCCCAATTCTGCGAACGAGTAGATTTAGT |
| 67 | TGTAATTTAGGCAGAGGCATTTTCGAGCCAGTAATAAG | 162 | TTGACCATTAGATACATTTCGCAAATGGTCAATAACCT |
| 68 | AGAATATAAAGTACCGACAAAAGGTAAAGTAATTCTGT | 163 | GTTTAGCTATATTTTCATTTGGGGCGCGAGCTGAAAAG |
| 69 | CCAGACGACGACAATAAACAACATGTTCAGCTAATGCA | 164 | GTGGCATCAATTCTACTAATAGTAGTAGCATTAACATC |
| 70 | GAACGCGCCTGTTTATCAACAATAGATAAGTCCTGAAC | 165 | CAATAAATCATACAGGCAAGGCAAAGAATTAGCAAAAT |
| 71 | AAGAAAAATAATATCCCATCCTAATTTACGAGCATGTA | 166 | TAAGCAATAAAGCCTCAGAGCATAAAGCTAAATCGGTT |
| 72 | GAAACCAATCAATAATCGGCTGTCTTTCCTTATCATTC | 167 | GTACCAAAAACATTATGACCCTGTAATACTTTTGCGGG |
| 73 | CAAGAACGGGTATTAAACCAAGTACCGCACTCATCGAG | 168 | AGAAGCCTTTATTTCAACGCAAGGATAAAAATTTTTAG |
| 74 | AACAAGCAAGCCGTTTTTATTTTCATCGTAGGAATCAT | 169 | AACCCTCATATATTTTAAATGCAATGCCTGAGTAATGT |
| 75 | TACCGCGCCCAATAGCAAGCAAATCAGATATAGAAGGC | 170 | GTAGGTAAAGATTCAAAAGGGTGAGAAAGGCCGGAGAC |
| 76 | TTATCCGGTATTCTAAGAACGCGAGGCGTTTTAGCGAA | 171 | AGTCAAATCACCATCAATATGATATTCAACCGTTCTAG |
| 77 | CCTCCCGACTTGCGGGAGGTTTTGAAGCCTTAAATCAA | 172 | CTGATAAATTAATGCCGGAGAGGGTAGCTATTTTTGAG |
| 78 | GATTAGTTGCTATTTTGCACCCAGCTACAATTTTATCC | 173 | AGATCTACAAAGGCTATCAGGTCATTGCCTGAGAGTCT |
| 79 | TGAATCTTACCAACGCTAACGAGCGTCTTTCCAGAGCC | 174 | GGAGCAAACAAGAGAATCGATGAACGGTAATCGTAAAA |
| 80 | TAATTTGCCAGTTACAAAATAAACAGCCATATTATTTA | 175 | CTAGCATGTCAATCATATGTACCCCGGTTGATAATCAG |
| 81 | TCCCAATCCAAATAAGAAACGATTTTTTGTTTAACGTC | 176 | AAAAGCCCCAAAAACAGGAAGATTGTATAAGCAAATAT |
| 82 | AAAAATGAAAATAGCAGCCTTTACAGAGAGAATAACAT | 177 | TTAAATTGTAAACGTTAATATTTTGTTAAAATTCGCAT |

| | | | |
|---|---|---|---|
| 83 | AAAAACAGGGAAGCGCATTAGACGGGAGAATTAACTGA | 178 | TAAATTTTTGTTAAATCAGCTCATTTTTTAACCAATAG |
| 84 | ACACCCTGAACAAAGTCAGAGGGTAATTGAGCGCTAAT | 179 | GAACGCCATCAAAAATAATTCGCGTCTGGCCTTCCTGT |
| 85 | ATCAGAGAGATAACCCACAAGAATTGAGTTAAGCCCAA | 180 | AGCCAGCTTTCATCAACATTAAATGTGAGCGAGTAACA |
| 86 | TAATAAGAGCAAGAAACAATGAAATAGCAATAGCTATC | 181 | ACCCGTCGGATTCTCCGTGGGAACAAACGGCGGATTGA |
| 87 | TTACCGAAGCCCTTTTTAAGAAAAGTAAGCAGATAGCC | 182 | CCGTAATGGGATAGGTCACGTTGGTGTAGATGGGCGCA |
| 88 | GAACAAAGTTACCAGAAGGAAACCGAGGAAACGCAATA | 183 | TCGTAACCGTGCATCTGCCAGTTTGAGGGGACGACGAC |
| 89 | ATAACGGAATACCCAAAAGAACTGGCATGATTAAGACT | 184 | AGTATCGGCCTCAGGAAGATCGCACTCCAGCCAGCTTT |
| 90 | CCTTATTACGCAGTATGTTAGCAAACGTAGAAAATACA | 185 | CCGGCACCGCTTCTGGTGCCGGAAACCAGGCAAAGCGC |
| 91 | TACATAAAGGTGGCAACATATAAAAGAAACGCAAAGAC | 186 | CATTCGCCATTCAGGCTGCGCAACTGTTGGGAAGGGCG |
| 92 | ACCACGGAATAAGTTTATTTTGTCACAATCAATAGAAA | 187 | ATCGGTGCGGGCCTCTTCGCTATTACGCCAGCTGGCGA |
| 93 | ATTCATATGGTTTACCAGCGCCAAAGACAAAAGGGCGA | 188 | AAGGGGGATGTGCTGCAAGGCGATTAAGTTGGGTAACG |
| 94 | CATTCAACCGATTGAGGGAGGGAAGGTAAATATTGACG | 189 | CCAGGGTTTTCCCAGTCACGACGTTGTAAAACGACGGC |
| 95 | GAAATTATTCATTAAAGGTGAATTATCACCGTCACCGA | 190 | CAGTGCCAAGCTTGCATGCCTGCAGGTCGACTCTAGAGGATCTTTT |

**Table S3.**

**Table S3.** DNA oligonucleotides for 4-colour ruler fabrication.

| Structural color | Replaced Oligonucleotide Number | Name | Sequence (5'→'3') |
|---|---|---|---|
| 1 | 14, 15 | B-1 | GCGATGGCCCACTACGTGAACCATC TTT GGATATCACTCATTAGTGGT |
| | | B-2 | ACCCAAATCAAGTTTTTTGGGGTCG |
| | | B-3 | AGGTGCCGTAAAGCACTAAATCGGAA |
| 2 | 28, 29 | B-4 | TTGCAACAGGAAAAACGCTCATGGA TTT GGATATCACTCATTAGTGGT |
| | | B-5 | AATACCTACATTTTGACGCTCAATC TTT GGATATCACTCATTAGTGGT |
| | | B-6 | GTCTGAAATGGATTATTTACATTGGC |
| 3 | 43, 44 | B-7 | TAATTTTAAAAGTTTGAGTAACATT TTT GGATATCACTCATTAGTGGT |
| | | B-8 | ATCATTTTGCGGAACAAAGAAACCA TTT GGATATCACTCATTAGTGGT |
| | | B-9 | CCAGAAGGAGCGGAATTATCATCATA TTT GGATATCACTCATTAGTGGT |
| 4 | 58, 59, 60, 61 | B-10 | TTATCAAAATCATAGGTCTGAGAGA TTT GGATATCACTCATTAGTGGT |
| | | B-11 | CTACCTTTTTAACCTCCGGCTTAGG TTT GGATATCACTCATTAGTGGT |
| | | B-12 | TTGGGTTATATAACTATATGTAAAT TTT GGATATCACTCATTAGTGGT |
| | | B-13 | GCTGATGCAAATCCAATCGCAAGAC TTT GGATATCACTCATTAGTGGT |
| | | B-14 | AAAGAACGCGAGAAAACTTTTTCAAA |
| | | B-15 | TATATTTTAGTTAATTTCATCTTCTG |
| | 'Imaging strand' | Bio-strand HPLC | ACCACTAATGAGTGATATCC/3'-biotin/ |

**Table S4.**

**Table S4.** DNA oligonucleotides for 10-colour ruler fabrication.

| Structural color | Replaced Oligonucleotide Number | Name | Sequence (5'→'3') |
|---|---|---|---|
| 1 | 14, 15 | B-1 | GCGATGGCCCACTACGTGAACCATC TTT GGATATCACTCATTAGTGGT |
| | | B-2 | ACCCAAATCAAGTTTTTTGGGGTCG |
| | | B-3 | AGGTGCCGTAAAGCACTAAATCGGAA |
| 2 | 28, 29 | B-4 | TTGCAACAGGAAAAACGCTCATGGA TTT GGATATCACTCATTAGTGGT |
| | | B-5 | AATACCTACATTTTGACGCTCAATC TTT GGATATCACTCATTAGTGGT |
| | | B-6 | GTCTGAAATGGATTATTTACATTGGC |
| 3 | 43, 44 | B-7 | TAATTTTAAAAGTTTGAGTAACATT TTT GGATATCACTCATTAGTGGT |
| | | B-8 | ATCATTTTGCGGAACAAAGAAACCA TTT GGATATCACTCATTAGTGGT |
| | | B-9 | CCAGAAGGAGCGGAATTATCATCATA TTT GGATATCACTCATTAGTGGT |
| 4 | 58, 59, 60, 61 | B-10 | TTATCAAAATCATAGGTCTGAGAGA TTT GGATATCACTCATTAGTGGT |
| | | B-11 | CTACCTTTTTAACCTCCGGCTTAGG TTT GGATATCACTCATTAGTGGT |
| | | B-12 | TTGGGTTATATAACTATATGTAAAT TTT GGATATCACTCATTAGTGGT |
| | | B-13 | GCTGATGCAAATCCAATCGCAAGAC TTT GGATATCACTCATTAGTGGT |
| | | B-14 | AAAGAACGCGAGAAAACTTTTTCAAA |
| | | B-15 | TATATTTTAGTTAATTTCATCTTCTG |
| 5 | 74, 75, 76, 77 | B-16 | AACAAGCAAGCCGTTTTTATTTTCA TTT GGATATCACTCATTAGTGGT |
| | | B-17 | TCGTAGGAATCATTACCGCGCCCAA TTT GGATATCACTCATTAGTGGT |
| | | B-18 | TAGCAAGCAAATCAGATATAGAAGG TTT GGATATCACTCATTAGTGGT |
| | | B-19 | CTTATCCGGTATTCTAAGAACGCGA TTT GGATATCACTCATTAGTGGT |
| | | B-20 | GGCGTTTTAGCGAACCTCCCGACTT TTT GGATATCACTCATTAGTGGT |
| | | B-21 | GCGGGAGGTTTTGAAGCCTTAAATCAA |
| 6 | 90, 91, 92, 93 | B-22 | CCTTATTACGCAGTATGTTAGCAAA TTT GGATATCACTCATTAGTGGT |
| | | B-23 | CGTAGAAAATACATACATAAAGGTG TTT GGATATCACTCATTAGTGGT |
| | | B-24 | GCAACATATAAAAGAAACGCAAAGA TTT GGATATCACTCATTAGTGGT |
| | | B-25 | CACCACGGAATAAGTTTATTTTGTC TTT GGATATCACTCATTAGTGGT |
| | | B-26 | ACAATCAATAGAAAATTCATATGGT TTT GGATATCACTCATTAGTGGT |

| | | | |
|---|---|---|---|
| | | B-27 | TTACCAGCGCCAAAGACAAAAGGGCGA TTT GGATATCACTCATTAGTGGT |
| 7 | 107, 108, 109, 110, 111 | B-28 | AGCGCAGTCTCTGAATTTACCGTTC TTT GGATATCACTCATTAGTGGT |
| | | B-29 | CAGTAAGCGTCATACATGGCTTTTG TTT GGATATCACTCATTAGTGGT |
| | | B-30 | ATGATACAGGAGTGTACTGGTAATA TTT GGATATCACTCATTAGTGGT |
| | | B-31 | AGTTTTAACGGGGTCAGTGCCTTGA TTT GGATATCACTCATTAGTGGT |
| | | B-32 | GTAACAGTGCCCGTATAAACAGTTA TTT GGATATCACTCATTAGTGGT |
| | | B-33 | ATGCCCCCTGCCTATTTCGGAACCT TTT GGATATCACTCATTAGTGGT |
| | | B-34 | ATTATTCTGAAACATGAAAGTATTA TTT GGATATCACTCATTAGTGGT |
| | | B-35 | AGAGGCTGAGACTCC |
| 8 | 125, 126, 127, 128, 129, 130 | B-36 | CTTTCGAGGTGAATTTCTTAAACAG TTT GGATATCACTCATTAGTGGT |
| | | B-37 | CTTGATACCGATAGTTGCGCCGACA TTT GGATATCACTCATTAGTGGT |
| | | B-38 | ATGACAACAACCATCGCCCACGCAT TTT GGATATCACTCATTAGTGGT |
| | | B-39 | AACCGATATATTCGGTCGCTGAGGC TTT GGATATCACTCATTAGTGGT |
| | | B-40 | TTGCAGGGAGTTAAAGGCCGCTTTT TTT GGATATCACTCATTAGTGGT |
| | | B-41 | GCGGGATCGTCACCCTCAGCAGCGA TTT GGATATCACTCATTAGTGGT |
| | | B-42 | AAGACAGCATCGGAACGAGGGTAGC TTT GGATATCACTCATTAGTGGT |
| | | B-43 | AACGGCTACAGAGGCTTTGAGGACT TTT GGATATCACTCATTAGTGGT |
| | | B-44 | AAAGACTTTTTCATGAGGAAGTTTCCAT |
| 9 | 143, 144, 145, 146, 147, 148 | B-45 | TATGCGATTTTAAGAACTGGCTCAT TTT GGATATCACTCATTAGTGGT |
| | | B-46 | TATACCAGTCAGGACGTTGGGAAGA TTT GGATATCACTCATTAGTGGT |
| | | B-47 | AAAATCTACGTTAATAAAACGAACT TTT GGATATCACTCATTAGTGGT |
| | | B-48 | AACGGAACAACATTATTACAGGTAG TTT GGATATCACTCATTAGTGGT |
| | | B-49 | AAAGATTCATCAGTTGAGATTTAGG TTT GGATATCACTCATTAGTGGT |
| | | B-50 | AATACCACATTCAACTAATGCAGAT TTT GGATATCACTCATTAGTGGT |
| | | B-51 | ACATAACGCCAAAAGGAATTACGAG TTT GGATATCACTCATTAGTGGT |
| | | B-52 | GCATAGTAAGAGCAACACTATCATA TTT GGATATCACTCATTAGTGGT |
| | | B-53 | ACCCTCGTTTACCAGACGACGATAA AAA TTT GGATATCACTCATTAGTGGT |
| 10 | 162, 163, 164, 165, 166, 167, 168, 169 | B-54 | TTGACCATTAGATACATTTCGCAAA TTT GGATATCACTCATTAGTGGT |
| | | B-55 | TGGTCAATAACCTGTTTAGCTATAT TTT GGATATCACTCATTAGTGGT |

| | | |
|---|---|---|
| | B-56 | TTTCATTTGGGGCGCGAGCTGAAAA TTT GGATATCACTCATTAGTGGT |
| | B-57 | GGTGGCATCAATTCTACTAATAGTA TTT GGATATCACTCATTAGTGGT |
| | B-58 | GTAGCATTAACATCCAATAAATCAT TTT GGATATCACTCATTAGTGGT |
| | B-59 | ACAGGCAAGGCAAAGAATTAGCAAA TTT GGATATCACTCATTAGTGGT |
| | B-60 | ATTAAGCAATAAAGCCTCAGAGCAT TTT GGATATCACTCATTAGTGGT |
| | B-61 | AAAGCTAAATCGGTTGTACCAAAAA TTT GGATATCACTCATTAGTGGT |
| | B-62 | CATTATGACCCTGTAATACTTTTGC TTT GGATATCACTCATTAGTGGT |
| | B-63 | GGGAGAAGCCTTTATTTCAACGCAA TTT GGATATCACTCATTAGTGGT |
| | B-64 | GGATAAAAATTTTTAGAACCCTCATAT |
| | B-65 | ATTTTAAATGCAATGCCTGAGTAATGT |
| 'Imaging strand' | Bio-strand HPLC | ACCACTAATGAGTGATATCC/3'-biotin/ |

**Table S5.**

**Table S5.** Oligonucleotides for 18S rRNA ID '1111'. DNA oligonucleotides that form structural color are highlighted in red.

| Name | Sequence (5'→3') | Length (nt) |
|---|---|---|
| 18S_rRNA_1 | TAATGATCCTTCCGCAGGTTCACCTACGGAAACCTTGT | 38 |
| 18S_rRNA_2 | TACGACTTTTACTTCCTCTAGATAGTCAAGTTCGACCG | 38 |
| 18S_rRNA_3 | TCTTCTCAGCGCTCCGCCAGGGCCGTGGGCCGACCCCG | 38 |
| 18S_rRNA_4 | GCGGGGCCGATCCGAGGGCCTCACTAAACCATCCAATC | 38 |
| 18S_rRNA_5 | GGTAGTAGCGACGGGCGGTGTGTACAAAGGGCAGGGAC | 38 |
| 18S_rRNA_6 | TTAATCAACGCAAGCTTATGACCCGCACTTACTGGGAA | 38 |
| 18S_rRNA_7 | TTCCTCGTTCATGGGGAATAATTGC | 25 |
| 18S_rRNA_8 | AATCCCCGATCCCCATCACGAATGG | 25 |
| 18S_rRNA_9 | GGTTCAACGGTCCTCTTTTGAGGAACAAGTTTTCTTGTGTTACCCGCG | 48 |
| 18S_rRNA_10 | CCTGCCGGCGTCCTCTTTTGAGGAACAAGTTTTCTTGTTAGGGTAGGC | 48 |
| 18S_rRNA_11 | ACACGCTGAGTCCTCTTTTGAGGAACAAGTTTTCTTGTCCAGTCAGTG | 48 |
| 18S_rRNA_12 | TAGCGCGCGTTCCTCTTTTGAGGAACAAGTTTTCTTGTGCAGCCCCGG | 48 |
| 18S_rRNA_13 | ACATCTAAGGTCCTCTTTTGAGGAACAAGTTTTCTTGTGCATCACAGA | 48 |
| 18S_rRNA_14 | CCTGTTATTGTCCTCTTTTGAGGAACAAGTTTTCTTGTCTCAATCTCG | 48 |
| 18S_rRNA_15 | GGTGGCTGAACGCCACTTGTCCCTCTAAGAAGTTGGGG | 38 |
| 18S_rRNA_16 | GACGCCGACCGCTCGGGGGTCGCGTAACTAGTTAGCAT | 38 |
| 18S_rRNA_17 | GCCAGAGTCTCGTTCGTTATCGGAATTAACCAGACAAA | 38 |
| 18S_rRNA_18 | TCGCTCCACCAACTAAGAACGGCCATGCACCACCACCC | 38 |
| 18S_rRNA_19 | ACGGAATCGAGAAAGAGCTATCAATCTGTCAATCCTGT | 38 |
| 18S_rRNA_20 | CCGTGTCCGGGCCGGGTGAGGTTTCCCGTGTTGAGTCA | 38 |
| 18S_rRNA_21 | AATTAAGCCGCAGGCTCCACTCCTG | 25 |
| 18S_rRNA_22 | GTGGTGCCCTTCCGTCAATTCCTTT | 25 |
| 18S_rRNA_23 | AAGTTTCAGCTCCTCTTTTGAGGAACAAGTTTTCTTGTTTTGCAACCA | 48 |
| 18S_rRNA_24 | TACTCCCCCCTCCTCTTTTGAGGAACAAGTTTTCTTGTGGAACCCAAA | 48 |
| 18S_rRNA_25 | GACTTTGGTTTCCTCTTTTGAGGAACAAGTTTTCTTGTTCCCGGAAGC | 48 |

| | | |
|---|---|---|
| 18S_rRNA_26 | TGCCCGGCGGTCCTCTTTTGAGGAACAAGTTTTCTTGTGTCATGGGAA | 48 |
| 18S_rRNA_27 | TAACGCCGCCTCCTCTTTTGAGGAACAAGTTTTCTTGTGCATCGCCGG | 48 |
| 18S_rRNA_28 | TCGGCATCGTTCCTCTTTTGAGGAACAAGTTTTCTTGTTTATGGTCGG | 48 |
| 18S_rRNA_29 | AACTACGACGGTATCTGATCGTCTTCGAACCTCCGACT | 38 |
| 18S_rRNA_30 | TTCGTTCTTGATTAATGAAAACATTCTTGGCAAATGCT | 38 |
| 18S_rRNA_31 | TTCGCTCTGGTCCGTCTTGCGCCGGTCCAAGAATTTCA | 38 |
| 18S_rRNA_32 | CCTCTAGCGGCGCAATACGAATGCCCCCGGCCGTCCCT | 38 |
| 18S_rRNA_33 | CTTAATCATGGCCTCAGTTCCGAAAACCAACAAAATAG | 38 |
| 18S_rRNA_34 | AACCGCGGTCCTATTCCATTATTCCTAGCTGCGGTATC | 38 |
| 18S_rRNA_35 | CAGGCGGCTCGGGCCTGCTTTGAAC | 25 |
| 18S_rRNA_36 | ACTCTAATTTTTTCAAAGTAAACGC | 25 |
| 18S_rRNA_37 | TTCGGGCCCCTCCTCTTTTGAGGAACAAGTTTTCTTGTGCGGGACACT | 48 |
| 18S_rRNA_38 | CAGCTAAGAGTCCTCTTTTGAGGAACAAGTTTTCTTGTCATCGAGGGG | 48 |
| 18S_rRNA_39 | GCGCCGAGAGTCCTCTTTTGAGGAACAAGTTTTCTTGTGCAAGGGGCG | 48 |
| 18S_rRNA_40 | GGGACGGGCGTCCTCTTTTGAGGAACAAGTTTTCTTGTGTGGCTCGCC | 48 |
| 18S_rRNA_41 | TCGCGGCGGATCCTCTTTTGAGGAACAAGTTTTCTTGTCCGCCCGCCC | 48 |
| 18S_rRNA_42 | GCTCCCAAGATCCTCTTTTGAGGAACAAGTTTTCTTGTTCCAACTACG | 48 |
| 18S_rRNA_43 | AGCTTTTTAACTGCAGCAACTTTAATATACGCTATTGG | 38 |
| 18S_rRNA_44 | AGCTGGAATTACCGCGGCTGCTGGCACCAGACTTGCCC | 38 |
| 18S_rRNA_45 | TCCAATGGATCCTCGTTAAAGGATTTAAAGTGGACTCA | 38 |
| 18S_rRNA_46 | TTCCAATTACAGGGCCTCGAAAGAGTCCTGTATTGTTA | 38 |
| 18S_rRNA_47 | TTTTTCGTCACTACCTCCCCGGGTCGGGAGTGGGTAAT | 38 |
| 18S_rRNA_48 | TTGCGCGCCTGCTGCCTTCCTTGGATGTGGTAGCCGTT | 38 |
| 18S_rRNA_49 | TCTCAGGCTCCCTCTCCGGAATCGA | 25 |
| 18S_rRNA_50 | ACCCTGATTCCCCGTCACCCGTGGT | 25 |
| 18S_rRNA_51 | CACCATGGTATCCTCTTTTGAGGAACAAGTTTTCTTGTGGCACGGCGA | 48 |
| 18S_rRNA_52 | CTACCATCGATCCTCTTTTGAGGAACAAGTTTTCTTGTAAGTTGATAG | 48 |
| 18S_rRNA_53 | GGCAGACGTTTCCTCTTTTGAGGAACAAGTTTTCTTGTCGAATGGGTC | 48 |
| 18S_rRNA_54 | GTCGCCGCCATCCTCTTTTGAGGAACAAGTTTTCTTGTCGGGGGGCGT | 48 |

| | | |
|---|---|---|
| 18S_rRNA_55 | GCGATCGGCCTCCTCTTTTGAGGAACAAGTTTTCTTGTCGAGGTTATC | 48 |
| 18S_rRNA_56 | TAGAGTCACCTCCTCTTTTGAGGAACAAGTTTTCTTGTAAAGCCGCCG | 48 |
| 18S_rRNA_57 | GCGCCCGCCCCCCGGCCGGGGCCGGAGAGGGGCTGACC | 38 |
| 18S_rRNA_58 | GGGTTGGTTTTGATCTGATAAATGCACGCATCCCCCCC | 38 |
| 18S_rRNA_59 | GCGAAGGGGGTCAGCGCCCGTCGGCATGTATTAGCTCT | 38 |
| 18S_rRNA_60 | AGAATTACCACAGTTATCCAAGTGGGAGAGGAGCGAGC | 38 |
| 18S_rRNA_61 | GACCAAAGGAACCATAACTGATTTAATGAGCCATTCGC | 38 |
| 18S_rRNA_62 | AGTTTCACTGTACCGGCCGTGCGTACTTAGACATGCAT | 38 |
| 18S_rRNA_63 | GGCTTAATCTTTGAGACAAGCATAT | 25 |
| 18S_rRNA_64 | GCTACTGGCAGGATCAACCAGGTA | 25 |

**Table S6.**

**Table S6.** Oligonucleotides for 28S rRNA ID '11111'. DNA oligonucleotides that form structural color are highlighted in red.

| Name | Sequence (5'→3') | Length (nt) |
|---|---|---|
| 28S_rRNA_1 | TCGGAACGGCGCTCGCCCATCTCTCAGGACCGACTGAC | 38 |
| 28S_rRNA_2 | CCATGTTCAACTGCTGTTCACATGGAACCCTTCTCCAC | 38 |
| 28S_rRNA_3 | TTCGGCCTTCAAAGTTCTCGTTTGAATATTTGCTACTA | 38 |
| 28S_rRNA_4 | CCACCAAGATCTGCACCTGCGGCGGCTCCACCCGGGCC | 38 |
| 28S_rRNA_5 | CGCGCCCTAGGCTTCAAGGCTCACCGCAGCGGCCCTCC | 38 |
| 28S_rRNA_6 | TACTCGTCGCGGCGTAGCGTCCGCGGGGCTCCGGGGGC | 38 |
| 28S_rRNA_7 | GGGGAGCGGGGCGTGGGCGGGAGGAGGGGAGGAGGCGT | 38 |
| 28S_rRNA_8 | GGGGGGGGGGGCGGGGGAAGGACCCCACACCCCCGCCG | 38 |
| 28S_rRNA_9 | CCGCCGCCGCCGCCGCCCTCCGACGCACACCACACGCG | 38 |
| 28S_rRNA_10 | CGCGCGCGCGCGCCGCCCCCGCCGCTCCCGTCCACTCT | 38 |
| 28S_rRNA_11 | CGACTGCCGGTCCTCTTTTGAGGAACAAGTTTTCTTGTCGACGGCCGG | 48 |
| 28S_rRNA_12 | GTATGGGCCCTCCTCTTTTGAGGAACAAGTTTTCTTGTGACGCTCCAG | 48 |
| 28S_rRNA_13 | CGCCATCCATTCCTCTTTTGAGGAACAAGTTTTCTTGTTTTCAGGGCT | 48 |
| 28S_rRNA_14 | AGTTGATTCGTCCTCTTTTGAGGAACAAGTTTTCTTGTGCAGGTGAGT | 48 |
| 28S_rRNA_15 | TGTTACACACTCCTCTTTTGAGGAACAAGTTTTCTTGTTCCTTAGCGG | 48 |
| 28S_rRNA_16 | ATTCCGACTTTCCTCTTTTGAGGAACAAGTTTTCTTGTCCATGGCCAC | 48 |
| 28S_rRNA_17 | CGTCCTGCTGTCTATATCAACCAACACCTTTTCTGGGG | 38 |
| 28S_rRNA_18 | TCTGATGAGCGTCGGCATCGGGCGCCTTAACCCGGCGT | 38 |
| 28S_rRNA_19 | TCGGTTCATCCCGCAGCGCCAGTTCTGCTTACCAAAAG | 38 |
| 28S_rRNA_20 | TGGCCCACTAGGCACTCGCATTCCACGCCCGGCTCCAC | 38 |
| 28S_rRNA_21 | GCCAGCGAGCCGGGCTTCTTACCCATTTAAAGTTTGAG | 38 |
| 28S_rRNA_22 | AATAGGTTGAGATCGTTTCGGCCCCAAGACCTCTAATC | 38 |
| 28S_rRNA_23 | ATTCGCTTTACCGGATAAAACTGCG | 25 |
| 28S_rRNA_24 | TGGCGGGGGTGCGTCGGGTCTGCGA | 25 |
| 28S_rRNA_25 | GAGCGCCAGCTCCTCTTTTGAGGAACAAGTTTTCTTGTTATCCTGAGG | 48 |
| 28S_rRNA_26 | GAAACTTCGGTCCTCTTTTGAGGAACAAGTTTTCTTGTAGGGAACCAG | 48 |

| | | |
|---|---|---|
| 28S_rRNA_27 | CTACTAGATGTCCTCTTTTGAGGAACAAGTTTTCTTGTGTTCGATTAG | 48 |
| 28S_rRNA_28 | TCTTTCGCCCTCCTCTTTTGAGGAACAAGTTTTCTTGTCTATACCCAG | 48 |
| 28S_rRNA_29 | GTCGGACGACTCCTCTTTTGAGGAACAAGTTTTCTTGTCGATTTGCAC | 48 |
| 28S_rRNA_30 | GTCAGGACCGTCCTCTTTTGAGGAACAAGTTTTCTTGTCTACGGACCT | 48 |
| 28S_rRNA_31 | CCACCAGAGTTTCCTCTGGCTTCGCCCTGCCCAGGCAT | 38 |
| 28S_rRNA_32 | AGTTCACCATCTTTCGGGTCCTAACACGTGCGCTCGTG | 38 |
| 28S_rRNA_33 | CTCCACCTCCCCGGCGCGGCGGGCGAGACGGGCCGGTG | 38 |
| 28S_rRNA_34 | GTGCGCCCTCGGCGGACTGGAGAGGCCTCGGGATCCCA | 38 |
| 28S_rRNA_35 | CCTCGGCCGGCGAGCGCGCCGGCCTTCACCTTCATTGC | 38 |
| 28S_rRNA_36 | GCCACGGCGGCTTTCGTGCGAGCCCCCGACTCGCGCAC | 38 |
| 28S_rRNA_37 | GTGTTAGACTCCTTGGTCCGTGTTT | 25 |
| 28S_rRNA_38 | CAAGACGGGTCGGGTGGGTAGCCGA | 25 |
| 28S_rRNA_39 | CGTCGCCGCCTCCTCTTTTGAGGAACAAGTTTTCTTGTGACCCCGTGC | 48 |
| 28S_rRNA_40 | GCTCGCTCCGTCCTCTTTTGAGGAACAAGTTTTCTTGTCCGTCCCCCT | 48 |
| 28S_rRNA_41 | CTTCGGGGGATCCTCTTTTGAGGAACAAGTTTTCTTGTCGCGCGCGTG | 48 |
| 28S_rRNA_42 | GCCCCGAGAGTCCTCTTTTGAGGAACAAGTTTTCTTGTAACCTCCCCC | 48 |
| 28S_rRNA_43 | GGGCCCGACGTCCTCTTTTGAGGAACAAGTTTTCTTGTGCGCGACCCG | 48 |
| 28S_rRNA_44 | CCCGGGGCGCTCCTCTTTTGAGGAACAAGTTTTCTTGTACTGGGGACA | 48 |
| 28S_rRNA_45 | GTCCGCCCCGCCCCCCGACCCGCGCGCGGCACCCCCCC | 38 |
| 28S_rRNA_46 | CGTCGCCGGGGCGGGGGCGCGGGGAGGAGGGGTGGGAG | 38 |
| 28S_rRNA_47 | AGCGGTCGCGCCGTGGGAGGGGTGGCCCGGCCCCCCCA | 38 |
| 28S_rRNA_48 | CGAGGAGACGCCGGCGCGCCCCCGCGGGGGAGACCCCC | 38 |
| 28S_rRNA_49 | CTCGCGGGGGATTCCCCGCGGGGGTGGGCGCCGGGAGG | 38 |
| 28S_rRNA_50 | GGGGAGAGCGCGGCGACGGGTCTCGCTCCCTCGGCCCC | 38 |
| 28S_rRNA_51 | GGGATTCGGCGAGTGCTGCTGCCGG | 25 |
| 28S_rRNA_52 | GGGGGCTGTAACACTCGGGGGGGGT | 25 |
| 28S_rRNA_53 | TTCGGTCCCGTCCTCTTTTGAGGAACAAGTTTTCTTGTCCGCCGCCGC | 48 |
| 28S_rRNA_54 | CGCCGCCGCCTCCTCTTTTGAGGAACAAGTTTTCTTGTACCGCCGCCG | 48 |
| 28S_rRNA_55 | CCGCCGCCGCTCCTCTTTTGAGGAACAAGTTTTCTTGTCCCGACCCGC | 48 |

| | | |
|---|---|---|
| 28S_rRNA_56 | GCGCCCTCCCTCCTCTTTTGAGGAACAAGTTTTCTTGTGAGGGAGGAC | 48 |
| 28S_rRNA_57 | GCGGGGCCGGTCCTCTTTTGAGGAACAAGTTTTCTTGTGGGGCGGAGA | 48 |
| 28S_rRNA_58 | CGGGGGAGGATCCTCTTTTGAGGAACAAGTTTTCTTGTGGAGGACGGA | 48 |
| 28S_rRNA_59 | CGGACGGACGGACGGGGCCCCCCGAGCCACCTTCCCCG | 38 |
| 28S_rRNA_60 | CCGGGCCTTCCCAGCCGTCCCGGAGCCGGTCGCGGCGC | 38 |
| 28S_rRNA_61 | ACCGCCGCGGTGGAAATGCGCCCGGCGGCGGCCGGTCG | 38 |
| 28S_rRNA_62 | CCGGTCGGGGGACGGTCCCCCGCCGACCCCACCCCCGG | 38 |
| 28S_rRNA_63 | CCCCGCCCGCCCACCCCCGCACCCGCCGGAGCCCGCCC | 38 |
| 28S_rRNA_64 | CCTCCGGGGAGGAGGAGGAGGGGCGGCGGGGGAAGGGA | 38 |
| 28S_rRNA_65 | GGGCGGGTGGAGGGGTCGGGAGGAA | 25 |
| 28S_rRNA_66 | CGGGGGGCGGGAAAGATCCGCCGGG | 25 |
| 28S_rRNA_67 | CCGCCGACACTCCTCTTTTGAGGAACAAGTTTTCTTGTGGCCGGACCC | 48 |
| 28S_rRNA_68 | GCCGCCGGGTTCCTCTTTTGAGGAACAAGTTTTCTTGTTGAATCCTCC | 48 |
| 28S_rRNA_69 | GGGCGGACTGTCCTCTTTTGAGGAACAAGTTTTCTTGTCGCGGACCCC | 48 |
| 28S_rRNA_70 | ACCCGTTTACTCCTCTTTTGAGGAACAAGTTTTCTTGTCTCTTAACGG | 48 |
| 28S_rRNA_71 | TTTCACGCCCTCCTCTTTTGAGGAACAAGTTTTCTTGTTCTTGAACTC | 48 |
| 28S_rRNA_72 | TCTCTTCAAATCCTCTTTTGAGGAACAAGTTTTCTTGTGTTCTTTTCA | 48 |
| 28S_rRNA_73 | ACTTTCCCTTACGGTACTTGTTGACTATCGGTCTCGTG | 38 |
| 28S_rRNA_74 | CCGGTATTTAGCCTTAGATGGAGTTTACCACCCGCTTT | 38 |
| 28S_rRNA_75 | GGGCTGCATTCCCAAGCAACCCGACTCCGGGAAGACCC | 38 |
| 28S_rRNA_76 | GGGCCCGGCGCGCCGGGGGCCGCTACCGGCCTCACACC | 38 |
| 28S_rRNA_77 | GTCCACGGGCTGGGCCTCGATCAGAAGGACTTGGGCCC | 38 |
| 28S_rRNA_78 | CCCACGAGCGGCGCCGGGGAGCGGGTCTTCCGTACGCC | 38 |
| 28S_rRNA_79 | ACATGTCCCGCGCCCCGCCGCGGGGCGGGGATTCGGCG | 38 |
| 28S_rRNA_80 | CTGGGCTCTTCCCTGTTCACTCGCCGTTACTGAGGGAA | 38 |
| 28S_rRNA_81 | TCCTGGTTAGTTTCTTCTCCTCCGCTGACTAATATGCT | 38 |
| 28S_rRNA_82 | TAAATTCAGCGGGTCGCCACGTCTGATCTGAGGTCGCG | 38 |

**Table S7.**

**Table S7.** Oligonucleotides for fabrication of partially complementary MS2 RNA ID ‘111’. DNA oligonucleotides that form structural color are highlighted in red.

| Name | Sequence (5’→3’) | Length (nt) |
|---|---|---|
| M_1 | CACTCCGTTCCCTACAACGAGCCTAAATTCATATGACT | 38 |
| M_2 | CGTTATAGCGGACCGCGTGTCTGATCCACGGCGCACAT | 38 |
| M_3 | TGGTCTCGGACCAATAGAGCCGCTCTCAGAGCGCGGGG | 38 |
| M_4 | GGTAACGGTTGCTTGTTCAGCGAACTTCTTGTAAGGCG | 38 |
| M_5 | CTGCATCCTGCAACTTGTGCCCCATAGGAGCACCGTTG | 38 |
| M_6 | GAGAACGTGCATTGCCCAAACAACGACGATCGGTAGCC | 38 |
| M_7 | AGAGAGGAGGTTGCCAATAAGGCTACGGATGCTGGTTT | 38 |
| M_8 | GTAAAACATCCGGATCCCATGACAAGGATTTGTCATGT | 38 |
| M_9 | AAGAAACCTTCTCTATTTATCTGACCGCGATCACCATT | 38 |
| M_10 | CGCCTCCCGTTCCTCTTTTGAGGAACAAGTTTTCTTGTAGCTTAGCGA | 48 |
| M_11 | TAGCTAAGGTTCCTCTTTTGAGGAACAAGTTTTCTTGTACGACGGGTC | 48 |
| M_12 | GCCTCGTCATTCCTCTTTTGAGGAACAAGTTTTCTTGTTACCAGAACC | 48 |
| M_13 | TAAGGTCGGATCCTCTTTTGAGGAACAAGTTTTCTTGTTGCTTTGTGA | 48 |
| M_14 | GCAATTCGTCTCCTCTTTTGAGGAACAAGTTTTCTTGTCCTTAAGTAA | 48 |
| M_15 | GCAATTGCTGTCCTCTTTTGAGGAACAAGTTTTCTTGTTAAAGTCGTC | 48 |
| M_16 | ACTGTGCGGATCACCGCTTCCAGTAGCGACAG | 32 |
| M_17 | AAGCAATTGATTGGTAAATTTCGAGAGAAAGATCGCGA | 38 |
| M_18 | GGAAGATCAATACATAAAGAGTTGAACTTCTTTGTTGT | 38 |
| M_19 | CTTCGACATGGGTAATCCTCATGTTTGAATGGCCGGCG | 38 |
| M_20 | TCTATTAGTAGATGCCGGAGTTTGCTGCGATTGCTGAG | 38 |
| M_21 | GGAATCGGGTTTCCATCTTTTAGGAGACCTTGCATTGC | 38 |
| M_22 | CTTAACAATAAGCTCGCAGTCGGAATTCGTAGCGAAAA | 38 |
| M_23 | TTGGAATGGTTAGTTCCATATTTAAGTACGAACGCCAT | 38 |
| M_24 | GCGGCTACAGGAAGCTCTACACCACCAACAGTCTGGGT | 38 |
| M_25 | TGCCACTTTAGGCACCTCGACTTTGATGGTGTATTTGC | 38 |

| | | |
|---|---|---|
| M_26 | GATTCTGCGCAGAGCTCTGACGAACGCTACAGGTTACT | 38 |
| M_27 | TTGTAAGCCTGTGAACGCGAGTTAGAGCTGATCCATTC | 38 |
| M_28 | AGCGACCCCGTTAGCGAAGTTGCTTGGGGCGACAGTCA | 38 |
| M_29 | CGTCGCCAGTTCCTCTTTTGAGGAACAAGTTTTCTTGTTCCGCCATTG | 48 |
| M_30 | TCGACGAGAATCCTCTTTTGAGGAACAAGTTTTCTTGTCGAACTGAGT | 48 |
| M_31 | AAAGTTAGAATCCTCTTTTGAGGAACAAGTTTTCTTGTGCCATGCTTC | 48 |
| M_32 | AAACTCCGGTTCCTCTTTTGAGGAACAAGTTTTCTTGTTGAGGGCTCT | 48 |
| M_33 | ATCTAGAGAGTCCTCTTTTGAGGAACAAGTTTTCTTGTCCGTTGCCTG | 48 |
| M_34 | ATTAATGCTATCCTCTTTTGAGGAACAAGTTTTCTTGTACGCATCTAA | 48 |
| M_35 | GGTATGGACCATCGAGAAAGGAGACTTTACGT | 32 |
| M_36 | ACGCGCCAGTTGTTGGCCATACGGATTGTACCCCTCGA | 38 |
| M_37 | TGCATGGCTGAGATTTGGGCCTTAGCAGTGCCCTGTCT | 38 |
| M_38 | CTCCACAGTCCACCCGTAGGGAGCGTCAACGCTTATGA | 38 |
| M_39 | TGGACTCACCCGTTATTACGTCAGTAACTGTTCCTGAC | 38 |
| M_40 | ATGTAGGAGCATCCCACGGGGGCCGTAAGGCCCTCGAG | 38 |
| M_41 | CATGTTACCTACAGGTAGGAGCCAGTCGACAACGAATG | 38 |
| M_42 | AGAAAGGCACCTTTTCCCACACTATACCTAGTGGGTTC | 38 |
| M_43 | AAGATACCTAGAGACGACAACCATGCCAAACGTGCATC | 38 |
| M_44 | GTTTATGTAAAACCATATCACGATACGTCGCGATATGT | 38 |
| M_45 | TGCACGTTGTTCCTCTTTTGAGGAACAAGTTTTCTTGTCTGGAAGTTT | 48 |
| M_46 | GCAGCTGGATTCCTCTTTTGAGGAACAAGTTTTCTTGTACGACAGACG | 48 |
| M_47 | GCCATCTAACTCCTCTTTTGAGGAACAAGTTTTCTTGTTTGATGTTAG | 48 |
| M_48 | TACCGACCTGTCCTCTTTTGAGGAACAAGTTTTCTTGTACGTACGGCT | 48 |
| M_49 | CTCATAGGAATCCTCTTTTGAGGAACAAGTTTTCTTGTGAAACTCTTG | 48 |
| M_50 | AAGGTGAACCTCCTCTTTTGAGGAACAAGTTTTCTTGTTTCGTAAGCA | 48 |
| M_51 | TCTCATATGCACCCTGGATATCACTCATTAGT | 32 |
| M_52 | GGTAACCAACCGAACTGCAACTCCAACCACCTGCCGGC | 38 |
| M_53 | CACGTGTTTTGATCGAAACTTTCGATCTTCGTTTAGGG | 38 |
| M_54 | CAAGGTAGCGGAGCGCCTGGCGCCAATTACCGCGACGA | 38 |

| | | |
|---|---|---|
| M_55 | GCGGCAGTGTACGCCTTCACGAGCGCAATGGTTTGCGT | 38 |
| M_56 | CGCGAGTTGTGAGGCTGTCGACCTGGCCTCTGCTAAAG | 38 |
| M_57 | CAACACCAAGGTTAAAATTACCCTGGGTGACCTTTTGC | 38 |
| M_58 | AGGACTTCGGTCGACGCCCGGTTCGCAACGTTCTGCGG | 38 |
| M_59 | CACTTCGATGTAAGTCAAGTTTTGGCTTACAGGGAAGA | 38 |
| M_60 | GGCTGTAGCAGGAGCGTGCGTCGAGGGAGAAGCCGAAA | 38 |

**Table S8.**

**Table S8.** Oligonucleotides for the MS2 RNA fully complementary ID ‘111’.

| Name | Sequence (5’→3’) | Length (nt) |
|---|---|---|
| M-1 | CCGGCTTTCTCCTCGTACGGGCGACCCCACGATGACCCACTTCGCTTGTAG | 51 |
| M-2 | GCACCTTGATCTATCGATGTGACACTTAACGCCCCCCGTGAATACGGAGA | 50 |
| M-3 | GGGGTAGTGCCACTGTTTCGTTTTGGCCCCAGTCGAGTTAAAACGACCGG | 50 |
| M-4 | GAGTCCAGTTCGAACGATATTTTAAAGAGAATGAGTTATCTTCAGTCTCA | 50 |
| M-5 | CCGTCCGCGTAAACGCGAACGGAGGGGACGAAGGTCTCGTTCTCCCTATC | 50 |
| M-6 | AAGGGTACTAAAAGCTCGCACAGGTCAAACCTCCTAGGAATGGAATTCCG | 50 |
| M-7 | GCTACCTACAGCGATAGCCATGGTAGCGTCTCGCTAAAGACATTAAAAAT | 50 |
| M-8 | GGCATTAGCTCGACAGGAAGTTGAGCAGGACCCCGAAAGGGGTCCCACCC | 50 |
| M61 | TGGGTGGTAACTAGCCAAGCAGCTAGTTACCAAATCGGGAGAATCCCGGG | 50 |
| M62 | TCCTCTCTTTAGGGGGAGGTCCCTGGGCCGAAGCCCGCCCACCTTTCGGT | 50 |
| M63 | GGAGCCGGACCGCTTTCGCACCCGTGCTCTTTCGAGCACACCCACCCCGT | 50 |
| M64 | TTACGGGGGTCCCTCGGTCAGCTACCGAGGAGAGCTCGCTGGCCCACACT | 50 |
| M65 | CCTGAGGGAATGTGGGAACCGGCGTTAGCCACTCCGAAGTGCGTATAACG | 50 |
| M66 | CGCACGCCGGCGGACTTCATGCTGTCGGTGATTTCACCTCCAGTATGGAA | 50 |
| M67 | CCACGCTATGTAGCGACCACTGTCGTGCTTTTCGCTGAAGAACTTGCGTT | 50 |
| M68 | CTCGAGCGATACGAGCAAGACGGAAACCCGAGGTACGGGTATCCGCGAGC | 50 |
| M69 | AGCCGCCCGTACGGAGTCTTGGTGTATACCGAGACTGCCGTAGGCGGGCT | 50 |
| M70 | GACTACGTAGTAGTCGGCAGCGAGGTCCGTCCCACCGAAGAACATCGAAG | 50 |
| M71 | GCACCTGGGAGGAGAGCCGTACCCACACCTTATAGAGGCGTGGATCTGAC | 50 |
| M72 | ATACCTCCGACAACTCCCCAACCCCGTAGCCGATTTAATATCAGCATCAG | 50 |
| M73 | GGCGAAGAGATTGTCAACAGGTTTCTTGATGTAAAACGGTTTGACATCGA | 50 |
| M74 | CACCACGGTAAAAGTGCGCGCCGCAGCTCTCGCGAAAGAGCCCGGACACG | 50 |
| M75 | AACGTTTTACGAAGATTCGGTTTAAAACCGTAGTAGGCAAGTGCCTCTAG | 50 |
| M76 | CACACGGGGTGCAATCTCACTGGGACATATAATATCGTCCCCGTAGATGC | 50 |
| M77 | CTATGGTTCCGGCGTTACCAAAATGGATTTGGGTCGCTTTGACTATTGCC | 50 |

| | | |
|---|---|---|
| M78 | CAGAATATCATGGACTCTAGCTCAAATGTGAACCCATTTCCCATTGTGGA | 50 |
| M79 | AAATAGTTCCCATCGTATCGTCTCGCCATCTACGATTCCGTAGTGTGAGC | 50 |
| M80 | GGATACGATCGAGATATGAATATAGCTCTGGTGGGAGAAAACTCCACACC | 50 |
| M81 | AGGCGATCGGAGATGGAATCGGATGCAGACGATAAGTCTATCGTCGCAAG | 50 |
| M82 | CGAACCATCTACGCTGCCCTGCTGAGCCAGACGCTGGTTGATCGATTGAT | 50 |
| M83 | CATTCAGGTCTATACCAACGGATTTGAGCCGGCGTCTGATGAAAGCACCG | 50 |
| M84 | ACCCCTTTCTGGAGGTACATATTCATATCAGGCTCCTTAC | 40 |
| M85 | AGGCAGCCCGATCTATTTTATTATTCTTCGGAACTGTAAA | 40 |

**Table S9.**

**Table S9.** Human universal total RNA contains equal quantities of DNase-treated total RNA from ten different human tissues/cell lines.

| Sample type | Origin |
|---|---|
| Adenocarcinoma | Mammary gland |
| Melanoma | Skin |
| Hepatoblastoma | Liver |
| Liposarcoma | Fat cells |
| Adenocarcinoma | Cervix |
| Histiocytic lymphoma | Macrophage and histiocyte |
| Embryonal carcinoma | Testis |
| Lymphoblastic leukemia | T lymphoblast |
| Glioblastoma | Brain |
| Plasmacytoma; myeloma | B lymphocyte |

**Table S10.**

**Table S10.** Oligonucleotides for MS2 RNA exons' ID fabrication.

| Name | Sequence (5'→3') | Length (nt) |
|---|---|---|
| M_AS_1 | TGGGTGGTAACTAGCCAAGCAGCTA | 25 |
| M_AS_2 | GTTACCAAATCGGGAGAATCCCGGGTCCTCTC | 30 |
| M_AS_3 | TTTAGGGGGAGGTCCCTGGGCCGAAGCCCGCCCACCTTTC | 40 |
| M_AS_4 | GGTGGAGCCGGACCGCTTTCGCACCCGTGCTCTTTCGAGC | 40 |
| M_AS_5 | ACACCCACCCCGTTTACGGGGGTCCCTCGGTCAGCTACCG | 40 |
| M_AS_6 | AGGAGAGCTCGCTGGCCCACACTCCTGAGGGAATGTGGGA | 40 |
| M_AS_7 | ACCGGCGTTAGCCACTCCGAAGTGCGTATAACGCGCACGC | 40 |
| M_AS_8 | CGGCGGACTTCATGCTGTCGGTGATTTCACCTCCAGTATG | 40 |
| M_AS_9 | GAACCACGCTATGTAGCGACCACTGTCGTGCTTTTCGCTG | 40 |
| M_AS_10 | AAGAACTTGCGTTCTCGAGCGATACGAGCAAGACGGAAAC | 40 |
| M_AS_11 | CCGAGGTACGGGTATCCGCGAGCAGCCGCCCGTACGGAGT | 40 |
| M_AS_12 | CTTGGTGTATACCGAGACTGCCGTAGGCGGGCTGACTACG | 40 |
| M_AS_13 | TAGTAGTCGGCAGCGAGGTCCGTCCCACCGAAGAACATCG | 40 |
| M_AS_14 | AAGGCACCTGGGAGGAGAGCCGTACCCACACCTTATAGAG | 40 |
| M_AS_15 | GCGTGGATCTGACATACCTCCGACAACTCCCCAACCCCGT | 40 |
| M_AS_16 | AGCCGATTTAATATCAGCATCAGGGCGAAGAGATTGTCAA | 40 |
| M_AS_17 | CAGGTTTCTTGATGTAAAACGGTTTGACATCGACACCACG | 40 |
| M_AS_18 | GTAAAAGTGCGCGCCGCAGCTCTCGCGAAAGAGCCCGGAC | 40 |
| M_AS_19 | ACGAACGTTTTACGAAGATTCGGTTTAAAACCGTAGTAGG | 40 |
| M_AS_20 | CAAGTGCCTCTAGCACACGGGGTGCAATCTCACTGGGACA | 40 |
| M_AS_21 | TATAATATCGTCCCCGTAGATGCCTATGGTTCCGGCGTTA | 40 |
| M_AS_22 | CCAAAATGGATTTGGGTCGCTTTGACTATTGCCCAGAATA | 40 |
| M_AS_23 | TCATGGACTCTAGCTCAAATGTGAA | 25 |
| M_AS_24 | CCCATTTCCCATTGTGGAAAATAGT | 25 |
| M_AS_25 | TCCCATCGTATCGTCTCGCCATCTA | 25 |

| | | |
|---|---|---|
| M_AS_26 | CGATTCCGTAGTGTGAGCGGATACGATCGAGATATGAATA | 40 |
| M_AS_27 | TAGCTCTGGTGGGAGAAAACTCCACACCAGGCGATCGGAG | 40 |
| M_AS_28 | ATGGAATCGGATGCAGACGATAAGTCTATCGTCGCAAGCG | 40 |
| M_AS_29 | AACCATCTACGCTGCCCTGCTGAGCCAGACGCTGGTTGAT | 40 |
| M_AS_30 | CGATTGATCATTCAGGTCTATACCAACGGATTTGAGCCGG | 40 |
| M_AS_31 | CGTCTGATGAAAGCACCGACCCCTTTCTGGAGGTACATAT | 40 |
| M_AS_32 | TCATATCAGGCTCCTTACAGGCAGCCCGATCTATTTTATT | 40 |
| M_AS_33 | ATTCTTCGGAACTGTAAACACTCCGTTCCCTACAACGAGC | 40 |
| M_AS_34 | CTAAATTCATATGACTCGTTATAGCGGACCGCGTGTCTGA | 40 |
| M_AS_35 | TCCACGGCGCACATTGGTCTCGGACCAATAGAGCCGCTCT | 40 |
| M_AS_36 | CAGAGCGCGGGGGGTAACGGTTGCTTGTTCAGCGAACTTC | 40 |
| M_AS_37 | TTGTAAGGCGCTGCATCCTGCAACTTGTGCCCCATAGGAG | 40 |
| M_AS_38 | CACCGTTGGAGAACGTGCATTGCCCAAACAACGACGATCG | 40 |
| M_AS_39 | GTAGCCAGAGAGGAGGTTGCCAATAAGGCTACGGATGCTG | 40 |
| M_AS_40 | GTTTGTAAAACATCCGGATCCCATGACAAGGATTTGTCAT | 40 |
| M_AS_41 | GTAAGAAACCTTCTCTATTTATCTGACCGCGATCACCATT | 40 |
| M_AS_42 | CGCCTCCCGTAGCTTAGCGATAGCTAAGGTACGACGGGTC | 40 |
| M_AS_43 | GCCTCGTCATTACCAGAACCTAAGGTCGGATGCTTTGTGA | 40 |
| M_AS_44 | GCAATTCGTCCCTTAAGTAAGCAATTGCTGTAAAGTCGTC | 40 |
| M_AS_45 | ACTGTGCGGATCACCGCTTCCAGTAGCGACAGAAGCAATT | 40 |
| M_AS_46 | GATTGGTAAATTTCGAGAGAAAGATCGCGAGGAAGATCAA | 40 |
| M_AS_47 | TACATAAAGAGTTGAACTTCTTTGT | 25 |
| M_AS_48 | TGTCTTCGACATGGGTAATCCTCAT | 25 |
| M_AS_49 | GTTTGAATGGCCGGCGTCTATTAGTAGATGCCGGAGTTTG | 40 |
| M_AS_50 | CTGCGATTGCTGAGGGAATCGGGTTTCCATCTTTTAGGAG | 40 |
| M_AS_51 | ACCTTGCATTGCCTTAACAATAAGCTCGCAGTCGGAATTC | 40 |
| M_AS_52 | GTAGCGAAAATTGGAATGGTTAGTTCCATATTTAAGTACG | 40 |
| M_AS_53 | AACGCCATGCGGCTACAGGAAGCTCTACACCACCAACAGT | 40 |
| M_AS_54 | CTGGGTTGCCACTTTAGGCACCTCGACTTTGATGGTGTAT | 40 |

| | | |
|---|---|---|
| M_AS_55 | TTGCGATTCTGCGCAGAGCTCTGACGAACGCTACAGGTTA | 40 |
| M_AS_56 | CTTTGTAAGCCTGTGAACGCGAGTTAGAGCTGATCCATTC | 40 |
| M_AS_57 | AGCGACCCCGTTAGCGAAGTTGCTTGGGGCGACAGTCACG | 40 |
| M_AS_58 | TCGCCAGTTCCGCCATTGTCGACGAGAACGAACTGAGTAA | 40 |
| M_AS_59 | AGTTAGAAGCCATGCTTCAAACTCCGGTTGAGGGCTCTAT | 40 |
| M_AS_60 | CTAGAGAGCCGTTGCCTGATTAATGCTAACGCATCTAAGG | 40 |
| M_AS_61 | TATGGACCATCGAGAAAGGAGACTTTACGTACGCGCCAGT | 40 |
| M_AS_62 | TGTTGGCCATACGGATTGTACCCCTCGATGCATGGCTGAG | 40 |
| M_AS_63 | ATTTGGGCCTTAGCAGTGCCCTGTCTCTCCACAGTCCACC | 40 |
| M_AS_64 | CGTAGGGAGCGTCAACGCTTATGATGGACTCACCCGTTAT | 40 |
| M_AS_65 | TACGTCAGTAACTGTTCCTGACATGTAGGAGCATCCCACG | 40 |
| M_AS_66 | GGGGCCGTAAGGCCCTCGAGCATGTTACCTACAGGTAGGA | 40 |
| M_AS_67 | GCCAGTCGACAACGAATGAGAAAGGCACCTTTTCCCACAC | 40 |
| M_AS_68 | TATACCTAGTGGGTTCAAGATACCTAGAGACGACAACCAT | 40 |
| M_AS_69 | GCCAAACGTGCATCGTTTATGTAAAACCATATCACGATAC | 40 |
| M_AS_70 | GTCGCGATATGTTGCACGTTGTCTG | 25 |
| M_AS_71 | GAAGTTTGCAGCTGGATACGACAGA | 25 |
| M_AS_72 | CGGCCATCTAACTTGATGTTAGTACCGACCTGACGTACGG | 40 |
| M_AS_73 | CTCTCATAGGAAGAAACTCTTGAAGGTGAACCTTCGTAAG | 40 |
| M_AS_74 | CATCTCATATGCACCCTGGATATCACTCATTAGTGGTAAC | 40 |
| M_AS_75 | CAACCGAACTGCAACTCCAACCACCTGCCGGCCACGTGTT | 40 |
| M_AS_76 | TTGATCGAAACTTTCGATCTTCGTTTAGGGCAAGGTAGCG | 40 |
| M_AS_77 | GAGCGCCTGGCGCCAATTACCGCGACGAGCGGCAGTGTAC | 40 |
| M_AS_78 | GCCTTCACGAGCGCAATGGTTTGCGTCGCGAGTTGTGAGG | 40 |
| M_AS_79 | CTGTCGACCTGGCCTCTGCTAAAGCAACACCAAGGTTAAA | 40 |
| M_AS_80 | ATTACCCTGGGTGACCTTTTGCAGGACTTCGGTCGACGCC | 40 |
| M_AS_81 | CGGTTCGCAACGTTCTGCGGCACTTCGATGTAAGTCAAGT | 40 |
| M_AS_82 | TTTGGCTTACAGGGAAGAGGCTGTAGCAGGAGCGTGCGTC | 40 |
| M_AS_83 | GAGGGAGAAGCCGAAACCGGCTTTCTCCTCGTACGGGCGA | 40 |

| | | |
|---|---|---|
| M_AS_84 | CCCCACGATGACCCACTTCGCTTGTAGGCACCTTGATCTA | 40 |
| M_AS_85 | TCGATGTGACACTTAACGCCCCCCGTGAATACGGAGAGGG | 40 |
| M_AS_86 | GTAGTGCCACTGTTTCGTTTTGGCCCCAGTCGAGTTAAAA | 40 |
| M_AS_87 | CGACCGGGAGTCCAGTTCGAACGATATTTTAAAGAGAATG | 40 |
| M_AS_88 | AGTTATCTTCAGTCTCACCGTCCGCGTAAACGCGAACGGA | 40 |
| M_AS_89 | GGGGACGAAGGTCTCGTTCTCCCTATCAAGGGTACTAAAA | 40 |
| M_AS_90 | GCTCGCACAGGTCAAACCTCCTAGGAATGGAATTCCGGCT | 40 |
| M_AS_91 | ACCTACAGCGATAGCCATGGTAGCGTCTCGCTAAAGACAT | 40 |
| M_AS_92 | TAAAAATGGCATTAGCTCGACAGGAAGTTGAG | 32 |
| M_AS_93 | CAGGACCCCGAAAGGGGTCCCACCC | 25 |

**Table S11.**

**Table S11.** Oligonucleotides are used for each exon type instead of oligonucleotides in Table S10.

| Exon | Sequence (5'→'3') | Length (nt) | Replaced Oligonucleotide Number |
|---|---|---|---|
| Exon I '112' | GAGATGGAATCGGATGCAGATGGGTGGTAACTAGCCAAGCAGCTA | 45 | 1 |
| | TCATGGACTCTAGCTCAAATGTGAA TTT GGATATCACTCATTAGTGGT | 48 | 23 |
| | TACATAAAGAGTTGAACTTCTTTGT TTT GGATATCACTCATTAGTGGT | 48 | 47 |
| | GTCGCGATATGTTGCACGTTGTCTG TTT GGATATCACTCATTAGTGGT | 48 | 70 |
| | GAAGTTTGCAGCTGGATACGACAGA TTT GGATATCACTCATTAGTGGT | 48 | 71 |
| | CAGGACCCCGAAAGGGGTCCCACCC CCTGATATGAATATGTACCT | 45 | 93 |
| Exon II '312' | TCTGCATCCGATTCCATCTCTGGGTGGTAACTAGCCAAGCAGCTA | 45 | 1 |
| | TCATGGACTCTAGCTCAAATGTGAA TTT GGATATCACTCATTAGTGGT | 48 | 23 |
| | CCCATTTCCCATTGTGGAAAATAGT TTT GGATATCACTCATTAGTGGT | 48 | 24 |
| | TCCCATCGTATCGTCTCGCCATCTA TTT GGATATCACTCATTAGTGGT | 48 | 25 |
| | TACATAAAGAGTTGAACTTCTTTGT TTT GGATATCACTCATTAGTGGT | 48 | 47 |
| | GTCGCGATATGTTGCACGTTGTCTG TTT GGATATCACTCATTAGTGGT | 48 | 70 |
| | GAAGTTTGCAGCTGGATACGACAGA TTT GGATATCACTCATTAGTGGT | 48 | 71 |
| | CAGGACCCCGAAAGGGGTCCCACCC CCTGATATGAATATGTACCT | 45 | 93 |
| Exon III '321' | TCTGCATCCGATTCCATCTCTGGGTGGTAACTAGCCAAGCAGCTA | 45 | 1 |
| | TCATGGACTCTAGCTCAAATGTGAA TTT GGATATCACTCATTAGTGGT | 48 | 23 |
| | CCCATTTCCCATTGTGGAAAATAGT TTT GGATATCACTCATTAGTGGT | 48 | 24 |
| | TCCCATCGTATCGTCTCGCCATCTA TTT GGATATCACTCATTAGTGGT | 48 | 25 |
| | TACATAAAGAGTTGAACTTCTTTGT TTT GGATATCACTCATTAGTGGT | 48 | 47 |
| | TGTCTTCGACATGGGTAATCCTCAT TTT GGATATCACTCATTAGTGGT | 48 | 48 |
| | GTCGCGATATGTTGCACGTTGTCTG TTT GGATATCACTCATTAGTGGT | 48 | 70 |
| | CAGGACCCCGAAAGGGGTCCCACCC AGGTACATATTCATATCAGG | 45 | 93 |
| Extended RNA | TCTGCATCCGATTCCATCTCTGGGTGGTAACTAGCCAAGCAGCTA | 45 | 1 |

**Table S12.**

**Table S12.** Oligonucleotides for the circular ID.

| Site | Sequence (5'→'3') | Length (nt) | Replaced Oligonucleotide Number |
|---|---|---|---|
| 1 | ACATCACTTGTCCTCTTTTGAGGAACAAGTTTTCTTGTCCTGAGTAGA | 48 | 26-30 |
| | AGAACTCAAATCCTCTTTTGAGGAACAAGTTTTCTTGTCTATCGGCCT | 48 | |
| | TGCTGGTAATTCCTCTTTTGAGGAACAAGTTTTCTTGTATCCAGAACA | 48 | |
| | ATATTACCGCTCCTCTTTTGAGGAACAAGTTTTCTTGTCAGCCATTGC | 48 | |
| | AACAGGAAAATCCTCTTTTGAGGAACAAGTTTTCTTGTACGCTCATGG | 48 | |
| | AAATACCTACTCCTCTTTTGAGGAACAAGTTTTCTTGTATTTTGACGC | 48 | |
| | TCAATCGTCTTCCTCTTTTGAGGAACAAGTTTTCTTGTGAAATGGATT | 48 | |
| | ATTTACATTGTCCTCTTTTGAGGAACAAGTTTTCTTGTGCAGATTCAC | 48 | |
| | CAGTCACACGACCAGTAATAAAAGGGACAT | 30 | |
| 2 | TTACCTGAGCAAAAGAAGATGATGAAACAAACATCAAGAAAACA | 44 | 52-57 |
| | AAATTAATTATCCTCTTTTGAGGAACAAGTTTTCTTGTCATTTAACAA | 48 | |
| | TTTCATTTGATCCTCTTTTGAGGAACAAGTTTTCTTGTATTACCTTTT | 48 | |
| | TTAATGGAAATCCTCTTTTGAGGAACAAGTTTTCTTGTCAGTACATAA | 48 | |
| | ATCAATATATTCCTCTTTTGAGGAACAAGTTTTCTTGTGTGAGTGAAT | 48 | |
| | AACCTTGCTTTCCTCTTTTGAGGAACAAGTTTTCTTGTCTGTAAATCG | 48 | |
| | TCGCTATTAATCCTCTTTTGAGGAACAAGTTTTCTTGTTTAATTTTCC | 48 | |
| | CTTAGAATCCTCCTCTTTTGAGGAACAAGTTTTCTTGTTTGAAAACAT | 48 | |
| | AGCGATAGCTTCCTCTTTTGAGGAACAAGTTTTCTTGTTAGATTAAGA | 48 | |
| | CGCTGAGAAGAGTCAATAGTGAAT | 24 | |
| 3 | TGAATCTTACCAACGCTAACGAGCGTCTTTCCAGAGCCTAATTTGCCAGT | 50 | 79-85 |
| | TACAAAATAATCCTCTTTTGAGGAACAAGTTTTCTTGTACAGCCATAT | 48 | |
| | TATTTATCCCTCCTCTTTTGAGGAACAAGTTTTCTTGTAATCCAAATA | 48 | |
| | AGAAACGATTTCCTCTTTTGAGGAACAAGTTTTCTTGTTTTTGTTTAA | 48 | |
| | CGTCAAAAATTCCTCTTTTGAGGAACAAGTTTTCTTGTGAAAATAGCA | 48 | |
| | GCCTTTACAGTCCTCTTTTGAGGAACAAGTTTTCTTGTAGAGAATAAC | 48 | |

| | |
|---|---|
| ATAAAAACAGTCCTCTTTTGAGGAACAAGTTTTCTTGTGGAAGCGCAT | 48 |
| TAGACGGGAGTCCTCTTTTGAGGAACAAGTTTTCTTGTAATTAACTGA | 48 |
| ACACCCTGAATCCTCTTTTGAGGAACAAGTTTTCTTGTCAAAGTCAGA | 48 |
| GGGTAATTGAGCGCTAATATCAGAGAGATAACCCACAAGAATTGAGTTAAGCCCAA | 56 |

**Table S13.**

**Table S13.** Oligonucleotides for fabrication of internal RNA origami ID.

| Name | Sequence (5'→3') | Length (nt) |
|---|---|---|
| internal M1 | TGGGTGGTAACTAGCCAAGCAGCTAGTTACCAAATCGG | 38 |
| internal M2 | GAGAATCCCGGGTCCTCTCTTTAGGGGGAGGTCCCTGG | 38 |
| internal M3 | GCCGAAGCCCGCCCACCTTTCGGTGGAGCCGGACCGCT | 38 |
| internal M4 | TTCGCACCCGTGCTCTTTCGAGCACACCCACCCCGTTT | 38 |
| internal M5 | ACGGGGGTCCCTCGGTCAGCTACCGAGGAGAGCTCGCT | 38 |
| internal M6 | GGCCCACACTCCTGAGGGAATGTGGGAACCGGCGTTAG | 38 |
| internal M7 | CCACTCCGAAGTGCGTATAACGCGCACGCCGGCGGACT | 38 |
| internal M8 | TCATGCTGTCGGTGATTTCACCTCCAGTATGGAACCAC | 38 |
| internal M9 | GCTATGTAGCGACCACTGTCGTGCTTTTCGCTGAAGAA | 38 |
| internal M10 | CTTGCGTTCTCGAGCGATACGAGCA | 25 |
| internal M11 | AGACGGAAACCCGAGGTACGGGTATC | 26 |
| internal M12 | CGCGAGCAGCCGCCCGTACGGAGTCTAGAGGCGTGGATCTGACATACCTC | 50 |
| internal M13 | CGACAACTCCCCAACCCCGTAGCCGA | 26 |
| internal M14 | TTTAATATCAGCATCAGGGCGAAGA | 25 |
| internal M15 | GATTGTCAACAGGTTTCTTGATGTAAAACGGTTTGACA | 38 |
| internal M16 | TCGACACCACGGTAAAAGTGCGCGCCGCAGCTCTCGCG | 38 |
| internal M17 | AAAGAGCCCGGACACGAACGTTTTACGAAGATTCGGTT | 38 |
| internal M18 | TAAAACCGTAGTAGGCAAGTGCCTCTAGCACACGGGGT | 38 |
| internal M19 | GCAATCTCACTGGGACATATAATATCGTCCCCGTAGAT | 38 |
| internal M20 | GCCTATGGTTCCGGCGTTACCAAAATGGATTTGGGTCG | 38 |
| internal M21 | CTTTGACTATTGCCCAGAATATCATGGACTCTAGCTCA | 38 |
| internal M22 | AATGTGAACCCATTTCCCATTGTGGAAAATAGTTCCCA | 38 |
| internal M23 | TCGTATCGTCTCGCCATCTACGATTCCGTAGTGTGAGC | 38 |
| internal M24 | GGATACGATCGAGATATGAATATAGCTCTGGTGGGAGA | 38 |
| internal M25 | AAACTCCACACCAGGCGATCGGAGATGGAATCGGATGC | 38 |

| | | |
|---|---|---|
| internal M26 | AGACGATAAGTCTATCGTCGCAAGCGAACCATCTACGC | 38 |
| internal M27 | TGCCCTGCTGAGCCAGACGCTGGTTGATCGATTGATCA | 38 |
| internal M28 | TTCAGGTCTATACCAACGGATTTGAGCCGGCGTCTGAT | 38 |
| internal M29 | GAAAGCACCGACCCCTTTCTGGAGGTACATATTCATAT | 38 |
| internal M30 | CAGGCTCCTTACAGGCAGCCCGATC | 25 |
| internal M31 | TATTTTATTATTCTTCGGAACTGTAA | 26 |
| internal M32 | ACACTCCGTTCCCTACAACGAGCCTAAATTCATATGACTCGTTATAGCGG | 50 |
| internal M33 | ACCGCGTGTCTGAACGGATGCTGGT | 25 |
| internal M34 | TTGTAAAACATCCGGATCCCATGACA | 26 |
| internal M35 | AGGATTTGTCATGTAAGAAACCTTCTCTATTTATCTGA | 38 |
| internal M36 | CCGCGATCACCATTCGCCTCCCGTAGCTTAGCGATAGC | 38 |
| internal M37 | TAAGGTACGACGGGTCGCCTCGTCATTACCAGAACCTA | 38 |
| internal M38 | AGGTCGGATGCTTTGTGAGCAATTCGTCCCTTAAGTAA | 38 |
| internal M39 | GCAATTGCTGTAAAGTCGTCACTGTGCGGATCACCGCT | 38 |
| internal M40 | TCCAGTAGCGACAGAAGCAATTGATTGGTAAATTTCGA | 38 |
| internal M41 | GAGAAAGATCGCGAGGAAGATCAATACATAAAGAGTTG | 38 |
| internal M42 | AACTTCTTTGTTGTCTTCGACATGGGTAATCCTCATGT | 38 |
| internal M43 | TTGAATGGCCGGCGTCTATTAGTAGATGCCGGAGTTTG | 38 |
| internal M44 | CTGCGATTGCTGAGGGAATCGGGTTTCCATCTTTTAGG | 38 |
| internal M45 | AGACCTTGCATTGCCTTAACAATAAGCTCGCAGTCGGA | 38 |
| internal M46 | ATTCGTAGCGAAAATTGGAATGGTTAGTTCCATATTTA | 38 |
| internal M47 | AGTACGAACGCCATGCGGCTACAGGAAGCTCTACACCA | 38 |
| internal M48 | CCAACAGTCTGGGTTGCCACTTTAGGCACCTCGACTTT | 38 |
| internal M49 | GATGGTGTATTTGCGATTCTGCGCAGAGCTCTGACGAA | 38 |
| internal M50 | CGCTACAGGTTACTTTGTAAGCCTGTGAACGCGAGTTA | 38 |
| internal M51 | GAGCTGATCCATTCAGCGACCCCGTTAGCGAAGTTGCT | 38 |
| internal M52 | TGGGGCGACAGTCACGTCGCCAGTTCCGCCATTGTCGA | 38 |
| internal M53 | CGAGAACGAACTGAGTAAAGTTAGAAGCCATGCTTCAA | 38 |
| internal M54 | ACTCCGGTTGAGGGCTCTATCTAGAGAGCCGTTGCCTG | 38 |

| | | |
|---|---|---|
| internal M55 | ATTAATGCTAACGCATCTAAGGTATGGACCATCGAGAA | 38 |
| internal M56 | AGGAGACTTTACGTACGCGCCAGTTGTTGGCCATACGG | 38 |
| internal M57 | ATTGTACCCCTCGATGCATGGCTGAGATTTGGGCCTTA | 38 |
| internal M58 | GCAGTGCCCTGTCTCTCCACAGTCCACCCGTAGGGAGC | 38 |
| internal M59 | GTCAACGCTTATGATGGACTCACCCGTTATTACGTCAG | 38 |
| internal M60 | TAACTGTTCCTGACATGTAGGAGCATCCCACGGGGGCC | 38 |
| internal M61 | GTAAGGCCCTCGAGCATGTTACCTACAGGTAGGAGCCA | 38 |
| internal M62 | GTCGACAACGAATGAGAAAGGCACCTTTTCCCACACTA | 38 |
| internal M63 | TACCTAGTGGGTTCAAGATACCTAGAGACGACAACCAT | 38 |
| internal M64 | GCCAAACGTGCATCGTTTATGTAAAACCATATCACGAT | 38 |
| internal M65 | ACGTCGCGATATGTTGCACGTTGTC | 25 |
| internal M66 | TGGAAGTTTGCAGCTGGATACGACAG | 26 |
| internal M67 | ACGGCCATCTAACTTGATGTTAGTAGCAACGTTCTGCGGCACTTCGATGT | 50 |
| internal M68 | AAGTCAAGTTTTGGCTTACAGGGAA | 25 |
| internal M69 | GAGGCTGTAGCAGGAGCGTGCGTCGA | 26 |
| internal M70 | GGGAGAAGCCGAAACCGGCTTTCTCCTCGTACGGGCGA | 38 |
| internal M71 | CCCCACGATGACCCACTTCGCTTGTAGGCACCTTGATC | 38 |
| internal M72 | TATCGATGTGACACTTAACGCCCCCCGTGAATACGGAG | 38 |
| internal M73 | AGGGGTAGTGCCACTGTTTCGTTTTGGCCCCAGTCGAG | 38 |
| internal M74 | TTAAAACGACCGGGAGTCCAGTTCGAACGATATTTTAA | 38 |
| internal M75 | AGAGAATGAGTTATCTTCAGTCTCACCGTCCGCGTAAA | 38 |
| internal M76 | CGCGAACGGAGGGGACGAAGGTCTCGTTCTCCCTATCA | 38 |
| internal M77 | AGGGTACTAAAAGCTCGCACAGGTCAAACCTCCTAGGA | 38 |
| internal M78 | ATGGAATTCCGGCTACCTACAGCGATAGCCATGGTAGC | 38 |
| internal M79 | GTCTCGCTAAAGACATTAAAAATGGCATTAGCTCGACA | 38 |
| internal M80 | GGAAGTTGAGCAGGACCCCGAAAGGGGTCCCACCC | 35 |

**Table S14.**

**Table S14.** Oligonucleotides for fabrication of terminal RNA origami ID.

| Name | Sequence (5'→3') | Length (nt) |
|---|---|---|
| terminal M1 | CACTCCGTTCCCTACAACGAGCCTAAATTCATATGACT | 38 |
| terminal M2 | CGTTATAGCGGACCGCGTGTCTGATCCACGGCGCACAT | 38 |
| terminal M3 | TGGTCTCGGACCAATAGAGCCGCTCTCAGAGCGCGGGG | 38 |
| terminal M4 | GGTAACGGTTGCTTGTTCAGCGAACTTCTTGTAAGGCG | 38 |
| terminal M5 | CTGCATCCTGCAACTTGTGCCCCATAGGAGCACCGTTG | 38 |
| terminal M6 | GAGAACGTGCATTGCCCAAACAACGACGATCGGTAGCC | 38 |
| terminal M7 | AGAGAGGAGGTTGCCAATAAGGCTACGGATGCTGGTTT | 38 |
| terminal M8 | GTAAAACATCCGGATCCCATGACAAGGATTTGTCATGT | 38 |
| terminal M9 | AAGAAACCTTCTCTATTTATCTGACCGCGATCACCATT | 38 |
| terminal M10 | CGCCTCCCGTAGCTTAGCGATAGCTAAGGTACGACGGGTC | 40 |
| terminal M11 | GCCTCGTCATTACCAGAACCTAAGGTCGGATGCTTTGTGA | 40 |
| terminal M12 | GCAATTCGTCCCTTAAGTAAGCAATTGCTGTAAAGTCGTC | 40 |
| terminal M13 | ACTGTGCGGATCACCGCTTCCAGTAGCGACAG | 32 |
| terminal M14 | AAGCAATTGATTGGTAAATTTCGAGAGAAAGATCGCGA | 38 |
| terminal M15 | GGAAGATCAATACATAAAGAGTTGAACTTCTTTGTTGT | 38 |
| terminal M16 | CTTCGACATGGGTAATCCTCATGTTTGAATGGCCGGCG | 38 |
| terminal M17 | TCTATTAGTAGATGCCGGAGTTTGCTGCGATTGCTGAG | 38 |
| terminal M18 | GGAATCGGGTTTCCATCTTTTAGGAGACCTTGCATTGC | 38 |
| terminal M19 | CTTAACAATAAGCTCGCAGTCGGAATTCGTAGCGAAAA | 38 |
| terminal M20 | TTGGAATGGTTAGTTCCATATTTAAGTACGAACGCCAT | 38 |
| terminal M21 | GCGGCTACAGGAAGCTCTACACCACCAACAGTCTGGGT | 38 |
| terminal M22 | TGCCACTTTAGGCACCTCGACTTTGATGGTGTATTTGC | 38 |
| terminal M23 | GATTCTGCGCAGAGCTCTGACGAACGCTACAGGTTACT | 38 |
| terminal M24 | TTGTAAGCCTGTGAACGCGAGTTAGAGCTGATCCATTC | 38 |
| terminal M25 | AGCGACCCCGTTAGCGAAGTTGCTTGGGGCGACAGTCA | 38 |
| terminal M26 | CGTCGCCAGTTCCGCCATTGTCGACGAGAACGAACTGAGT | 40 |

| | | |
|---|---|---|
| terminal M27 | AAAGTTAGAAGCCATGCTTCAAACTCCGGTTGAGGGCTCT | 40 |
| terminal M28 | ATCTAGAGAGCCGTTGCCTGATTAATGCTAACGCATCTAA | 40 |
| terminal M29 | GGTATGGACCATCGAGAAAGGAGACTTTACGT | 32 |
| terminal M30 | ACGCGCCAGTTGTTGGCCATACGGATTGTACCCCTCGA | 38 |
| terminal M31 | TGCATGGCTGAGATTTGGGCCTTAGCAGTGCCCTGTCT | 38 |
| terminal M32 | CTCCACAGTCCACCCGTAGGGAGCGTCAACGCTTATGA | 38 |
| terminal M33 | TGGACTCACCCGTTATTACGTCAGTAACTGTTCCTGAC | 38 |
| terminal M34 | ATGTAGGAGCATCCCACGGGGGCCGTAAGGCCCTCGAG | 38 |
| terminal M35 | CATGTTACCTACAGGTAGGAGCCAGTCGACAACGAATG | 38 |
| terminal M36 | AGAAAGGCACCTTTTCCCACACTATACCTAGTGGGTTC | 38 |
| terminal M37 | AAGATACCTAGAGACGACAACCATGCCAAACGTGCATC | 38 |
| terminal M38 | GTTTATGTAAAACCATATCACGATACGTCGCGATATGT | 38 |
| terminal M39 | TGCACGTTGTCTGGAAGTTTGCAGCTGGATACGACAGACG | 40 |
| terminal M40 | GCCATCTAACTTGATGTTAGTACCGACCTGACGTACGGCT | 40 |
| terminal M41 | CTCATAGGAAGAAACTCTTGAAGGTGAACCTTCGTAAGCA | 40 |
| terminal M42 | TCTCATATGCACCCTGGATATCACTCATTAGT | 32 |
| terminal M43 | GGTAACCAACCGAACTGCAACTCCAACCACCTGCCGGC | 38 |
| terminal M44 | CACGTGTTTTGATCGAAACTTTCGATCTTCGTTTAGGG | 38 |
| terminal M45 | CAAGGTAGCGGAGCGCCTGGCGCCAATTACCGCGACGA | 38 |
| terminal M46 | GCGGCAGTGTACGCCTTCACGAGCGCAATGGTTTGCGT | 38 |
| terminal M47 | CGCGAGTTGTGAGGCTGTCGACCTGGCCTCTGCTAAAG | 38 |
| terminal M48 | CAACACCAAGGTTAAAATTACCCTGGGTGACCTTTTGC | 38 |
| terminal M49 | AGGACTTCGGTCGACGCCCGGTTCGCAACGTTCTGCGG | 38 |
| terminal M50 | CACTTCGATGTAAGTCAAGTTTTGGCTTACAGGGAAGA | 38 |
| terminal M51 | GGCTGTAGCAGGAGCGTGCGTCGAGGGAGAAGCCGAAA | 38 |

**Table S15.**

**Table S15.** Oligonucleotides for the *ENO1*.

| Name | Sequence (5’→3’) | Length (nt) |
|---|---|---|
| ENO1_1 | TATTCTCATGGGTCACTGAGGCTTTTTATTTTGAGCAC | 38 |
| ENO1_2 | AAAACCACCGGGGATCTAGCCTGTGGCCACCCCGGAGA | 38 |
| ENO1_3 | TGACACGAGGCTCACATGACTCTAGACACTTGGTGGAA | 38 |
| ENO1_4 | AGTGAGGCGAGAAAAACAATGACTTGGGCCAATTACAC | 38 |
| ENO1_5 | GACTGCAAAGCTAGAGCTGCCAACAGGGCTCCAGGGAG | 38 |
| ENO1_6 | CTTGGCTTCTGTAGAAGTTCTAAGGAAGCGGTACGAAC | 38 |
| ENO1_7 | TCCACGGCGGTGGGGCGCTAACTAGCAGGGACCCCTGC | 38 |
| ENO1_8 | AAGTGTTGGTCGGGGGCCTCGAGCTGCCTGAGCTGACA | 38 |
| ENO1_9 | CGAGGGGAGGGGTCTGTGTAGCCAACAGGTGACCGAAG | 38 |
| ENO1_10 | GGCTTGCCTGCCCACAGCTTACTTGGCCAAGGGGTTTC | 38 |
| ENO1_11 | TGAAGTTCCTGCCGGCAAACTTAGCCTTGCTGCCCAGC | 38 |
| ENO1_12 | TCCTCTTCAATTCTGAGGAGCTGGTTGTACTTGGCCAA | 38 |
| ENO1_13 | GCGCTCAGATCGGCAAGGGGCACCAGTCTTGATCTGCC | 38 |
| ENO1_14 | CAGTGCACAGCCCCACAACCAGGTCAGCGATGAAGGTA | 38 |
| ENO1_15 | TCTTCAGTCTCCCCCGAACGATGAGACACCATGA | 34 |
| ENO1_16 | CGCCCCAACCTCCTCTTTTGAGGAACAAGTTTTCTTGTATTGGCCTGG | 48 |
| ENO1_17 | GCCAGCTTGCTCCTCTTTTGAGGAACAAGTTTTCTTGTACGCCTGAAG | 48 |
| ENO1_18 | AGACTCGGTCTCCTCTTTTGAGGAACAAGTTTTCTTGTACGGAGCCAA | 48 |
| ENO1_19 | TCTGGTTGACTCCTCTTTTGAGGAACAAGTTTTCTTGTTTTGAGCAGG | 48 |
| ENO1_20 | AGGCAGTTGCTCCTCTTTTGAGGAACAAGTTTTCTTGTAGGACTTCTC | 48 |
| ENO1_21 | GTTCACGGCCTCCTCTTTTGAGGAACAAGTTTTCTTGTTTGGCGATCC | 48 |
| ENO1_22 | TCTTTGGGTTGGTCACTGTGAGATCATCCCCCACTACC | 38 |
| ENO1_23 | TGGATTCCTGCACTGGCTGTGAACTTCTGCCAAGCTCC | 38 |
| ENO1_24 | CCAGTCATCCTGGTCAAAGGGATCTTCGATAGACACCA | 38 |
| ENO1_25 | CTGGGTAGTCCTTGATGAAGGACTTGTACAGGTCAGCC | 38 |
| ENO1_26 | AGCTGGTCAGGCGAGATGTACCTGCTGGGGTCATCGGG | 38 |

| | | |
|---|---|---|
| ENO1_27 | AGACTTGAAGTCCAGGTCATACTTCCCAGACCTGAAGA | 38 |
| ENO1_28 | ACTCGGAGGCCGCTACGTCCATGCCGATGACCACCTTATCAGTGTAGCCA | 48 |
| ENO1_29 | GCTTTCCCAATCCTCTTTTGAGGAACAAGTTTTCTTGTTAGCAGTCTT | 48 |
| ENO1_30 | CAGCAGCTCCTCCTCTTTTGAGGAACAAGTTTTCTTGTAGGCCTTCTT | 48 |
| ENO1_31 | TATTCTCCAGTCCTCTTTTGAGGAACAAGTTTTCTTGTGATGTTGGGA | 48 |
| ENO1_32 | GCAAACCCGCTCCTCTTTTGAGGAACAAGTTTTCTTGTCTTCATCCCC | 48 |
| ENO1_33 | CACATTGGTGTCCTCTTTTGAGGAACAAGTTTTCTTGTGCATCTTTCC | 48 |
| ENO1_34 | CATATTTCTCTCCTCTTTTGAGGAACAAGTTTTCTTGTCTTGATGACA | 48 |
| ENO1_35 | TTCTTCAGGTTGTGGTAAACCTCTGCTCCAATGCGCAT | 38 |
| ENO1_36 | GGCTTCCCTGAAGTTTGCTGCACCGACTGGGAGGATCA | 38 |
| ENO1_37 | TGAACTCCTGCATGGCCAGCTTGTTGCCAGCATGAGAA | 38 |
| ENO1_38 | CCGCCATTGATGACATTGAACGCCGGGACTGGCAGGAT | 38 |
| ENO1_39 | GACTTCAGAGTTGCCAGCCAAGTCAGCGATGTGGCGGT | 38 |
| ENO1_40 | ACAGGGGGACCCCCTTCTCAACGGCACCAGCTTTGCAG | 38 |
| ENO1_41 | ACGGCAAGGGACACCCCCAGAATGGCGTTCGCACCAAACTTAGATTTATT | 48 |
| ENO1_42 | TTCTGTTCCATCCTCTTTTGAGGAACAAGTTTTCTTGTTCCATCTCGA | 48 |
| ENO1_43 | TCATCAGTTTTCCTCTTTTGAGGAACAAGTTTTCTTGTGTCAATCTTC | 48 |
| ENO1_44 | TCTTGTTCTGTCCTCTTTTGAGGAACAAGTTTTCTTGTTGACGTTCAG | 48 |
| ENO1_45 | TTTCTTGCTATCCTCTTTTGAGGAACAAGTTTTCTTGTACCAGGGCAG | 48 |
| ENO1_46 | GCGCAATAGTTCCTCTTTTGAGGAACAAGTTTTCTTGTTTTATTGATG | 48 |
| ENO1_47 | TGCTCAACAGTCCTCTTTTGAGGAACAAGTTTTCTTGTCCTTTGAGAC | 48 |
| ENO1_48 | ACCTGGAAGGAACAATCAAATCAAATTACTGACATTAA | 38 |
| ENO1_49 | CTTTAAAACAGGCTTGAGCAGCATTGTTAGAGAGAACC | 38 |
| ENO1_50 | ACTGTTGAGGCCGACGCGGAAGCCCACCAAGGGACCTT | 38 |
| ENO1_51 | CTGTGGGACCTCTTCCCCCTCTCCCCTTTCCCCCCAGA | 38 |
| ENO1_52 | ATCGGAGGACTTTCCGGCCCAGTCTGAGTGCTCCTACC | 38 |
| ENO1_53 | TAGGACCCCAGAGGGTTAGAGCCACACAAAACAGCAGC | 38 |
| ENO1_54 | TGGCTCAGAACGATCTGACTGGAGGTTAGTTTGCAAGA | 38 |
| ENO1_55 | CCATGCACTGGAAACGCCAACTGCAGACTAAAGGCTGT | 38 |

| | | |
|---|---|---|
| ENO1_56 | TCAATCAGTTACCTGAGTGCACAGGGCTGGTGCCAGGA | 38 |
| ENO1_57 | ACTCATTAATATACTTAATGGGTCTGGAGACGCCTGCC | 38 |
| ENO1_58 | CAGCAGAGGATGAAGAAGGAAAAATGAGAGAGACTCAG | 38 |
| ENO1_59 | AAGAGAACCGGTGATTAAGGACTGCGAGTGAGAGACAG | 38 |
| ENO1_60 | TGAGGGGGCAGGAAAGCAGAGGAGAGCATTTCAGGGGC | 38 |
| ENO1_61 | CATGCGTTCAATCTTCCACAAATGCTTACTTACTGAGC | 38 |
| ENO1_62 | ACTCATTACCATCAAGGCACAGCATCATCCTGCTCCGG | 38 |
| ENO1_63 | CTAAGTCCCCACGTACGCCATTAAACAACGGTCAAATG | 38 |
| ENO1_64 | GTAACATGTTCGTGTGTGTAGAGTGCTTCTTACAGCAT | 38 |
| ENO1_65 | TAAAGCGGGACTGAACACCCAGCACCACCACTGCGACA | 38 |
| ENO1_66 | CCAGCCCAGTGTTTCACAACCTGATTTCATCATTGTCC | 38 |
| ENO1_67 | CCTCCCTGCAGCCTTTTTGGAGTTTTCCGGGTGCCCCA | 38 |
| ENO1_68 | CTGAGAATGCGTGATCTAGCCCTATGTGCTTTTCTGTA | 38 |
| ENO1_69 | ATTTGGCCACATGTTCTATCTCTAGAAAGGTCAATGTT | 38 |
| ENO1_70 | GCACACACACTAAGTCAATTTGAATA | 26 |
| ENO1_71 | CAGCCAGAGGTCCTCTTTTGAGGAACAAGTTTTCTTGTCTAGCTCTGC | 48 |
| ENO1_72 | CACAGAGCAGTCCTCTTTTGAGGAACAAGTTTTCTTGTGGCTCCTATG | 48 |
| ENO1_73 | AGTCAGTCAATCCTCTTTTGAGGAACAAGTTTTCTTGTCAGGAAAATC | 48 |
| ENO1_74 | ACACTAAATGTCCTCTTTTGAGGAACAAGTTTTCTTGTACACACCTGC | 48 |
| ENO1_75 | AGCTCACCATTCCTCTTTTGAGGAACAAGTTTTCTTGTTCAGGGCAGG | 48 |
| ENO1_76 | TCATTGGTTATCCTCTTTTGAGGAACAAGTTTTCTTGTGACCTTCCCT | 48 |
| ENO1_77 | GGATCCTTTCTAGCCCGTGAAGAGGCTGATGATCTCTGGAGGGCCCA | 47 |
| ENO1_78 | CAGAACCCTCCTGGGAGATGGGGAAACTTTTCTTTAGTTAACAGGCC | 47 |
| ENO1_79 | CTGTTCTCATGCTTGGGTTCCTGTCCTAACCGTGAAGGAGTATTTCG | 47 |
| ENO1_80 | CAGATTGGGAACAAACTGCAATAGAAGCAAGCATTGGAACTCTCGAC | 47 |
| ENO1_81 | CCAGAGCATGCAGGCTCGGGAACCCCGGAATCCACACACCAACTTTC | 47 |
| ENO1_82 | CTGTCCACAAGGTGGTGCACTGCTTCCATCAACATCATGGGTC | 43 |
| ENO1_83 | ACAGCAGGTTTCCTCTTTTGAGGAACAAGTTTTCTTGTTACCTGCCAT | 48 |
| ENO1_84 | AAACCTGCAATCCTCTTTTGAGGAACAAGTTTTCTTGTGTGCAAGTGC | 48 |

| | | |
|---|---|---|
| ENO1_85 | CACCCAGAGATCCTCTTTTGAGGAACAAGTTTTCTTGTGGACAGGACT | 48 |
| ENO1_86 | GGGCAAGGCCTCCTCTTTTGAGGAACAAGTTTTCTTGTAGGAATGGGG | 48 |
| ENO1_87 | CTGGAAGCAGTCCTCTTTTGAGGAACAAGTTTTCTTGTGGAGGCCATG | 48 |
| ENO1_88 | GGCTGTGGGTTCCTCTTTTGAGGAACAAGTTTTCTTGTTCTAAGGCTT | 48 |
| ENO1_89 | ACCCTTCCCCATATAGCGAGTCTTATCATTGTCCCGGAGCTCTAGGGCCT | 50 |
| ENO1_90 | CATAGATACCAGTTGAAGCACCACTGGGCACAGCAGCTCTGAAGAGACCT | 50 |
| ENO1_91 | TTTGAGGTGAAGAGATCAACCTCAACAGTGGGATTCCCGCGAGAGTCAAA | 50 |
| ENO1_92 | GATCTCCCTGGCATGGATCTTGAGAATAGACATGGTGAACTTCTAGCCAC | 50 |
| ENO1_93 | TGGGTCTCGTCGCCTAGGAGAGGAAGCGGAGGGTGCTGCAGACACCGAGG | 50 |
| ENO1_94 | TGAACGTAAAGCCGGCGAGATCTCCGTGCTCCGGGTACCCACAGATACT | 49 |
| ENO1_95 | GAGTGCACAGGGCTGGTGCCAGGAACTC | 28 |
| ENO1_96 | ATTAATATACTTAATGGGTCTGGAGACG | 28 |

**Table S16.**

**Table S16**. *Xist* lncRNA ID '111111' oligonucleotides. DNA oligonucleotides that form structural color are highlighted in red.

| Name | Sequence (5'→3') | Length (nt) |
|---|---|---|
| Xist_1 | TTTTTTTTTTTTTTTTCAAAGTTTTCAAAACAGTATAT | 38 |
| Xist_2 | TTTATTTTACAATAGCAACCAACTCCCCAGTTTGTTTC | 38 |
| Xist_3 | AATTGTGACATCTAGATGGCTTAAGATTACTTTCTGGT | 38 |
| Xist_4 | GGTCACCCATGCTGAACAATATTTTTCAATCTTCCAAA | 38 |
| Xist_5 | CAGCAAAGACTCAAAAGAGATTCTGCATTTCACATCAG | 38 |
| Xist_6 | TTCACAAGTTCAAGAGTCTTCCATTTATCTTAGCTTTT | 38 |
| Xist_7 | GGAATAAATTATCTTTGAGGTAGAAGGACAATGACGAA | 38 |
| Xist_8 | GCCACTTAATTCCTTGTGTCTGCATAAAAGCAGATTTA | 38 |
| Xist_9 | TTCATCACAACTTCATTTATGTGAATAAAGCAGATGAT | 38 |
| Xist_10 | GATAAAATGTTCTCTTATTCTTGTTTAATCAGTAGTGG | 38 |
| Xist_11 | TAGTGATGCCTCCTCTTTTGAGGAACAAGTTTTCTTGTAGAAACTGTG | 48 |
| Xist_12 | AAAGGAAGGCTCCTCTTTTGAGGAACAAGTTTTCTTGTTTTTAGTTAC | 48 |
| Xist_13 | TTTCTTCTTTTCCTCTTTTGAGGAACAAGTTTTCTTGTCCATTTTCCA | 48 |
| Xist_14 | ATAATCCATTTCCTCTTTTGAGGAACAAGTTTTCTTGTCCCCATCCCC | 48 |
| Xist_15 | AGCTGAAGAATCCTCTTTTGAGGAACAAGTTTTCTTGTAGGGGTGTTA | 48 |
| Xist_16 | CTGAGTCCAGTCCTCTTTTGAGGAACAAGTTTTCTTGTCTGATACCAC | 48 |
| Xist_17 | ACATTGAAAGGTAAACATTAATATTTCAAATCTGATGT | 38 |
| Xist_18 | CTAACTAAAAATGTACAGAATGAAAACTAGAAAATTTC | 38 |
| Xist_19 | AACCCCAGATTATCTTCAACCTTGCTCCCTCCACCAAT | 38 |
| Xist_20 | CATACTTTGACATTTATCTATTTCCTTCTCCACTTATG | 38 |
| Xist_21 | GATGTAATTGGCTTGCTATAGAAACTACAGTTCAGATG | 38 |
| Xist_22 | CTTTGAATGTATGAACTACAATGAACAATAAAGTCCTC | 38 |
| Xist_23 | TTCTTTTGAAGCATATTTTGGCTTC | 25 |
| Xist_24 | AGCTTTAAGATAATCTTATGACAAG | 25 |
| Xist_25 | AAGGGTCACATCCTCTTTTGAGGAACAAGTTTTCTTGTCTGATTCACT | 48 |

| | | |
|---|---|---|
| Xist_26 | TAATAAATTCTCCTCTTTTGAGGAACAAGTTTTCTTGTCATTCTTACC | 48 |
| Xist_27 | TAACACAAGGTCCTCTTTTGAGGAACAAGTTTTCTTGTTTTAGTTGAT | 48 |
| Xist_28 | AAGCACTTGGTCCTCTTTTGAGGAACAAGTTTTCTTGTACAAAAATAA | 48 |
| Xist_29 | TACTTTTCAATCCTCTTTTGAGGAACAAGTTTTCTTGTAAATGTAAAG | 48 |
| Xist_30 | CAAACTAGTGTCCTCTTTTGAGGAACAAGTTTTCTTGTAGGACAAAGG | 48 |
| Xist_31 | ATTTTGTCCTCATCTCAACAATGATCAGCTATTGGAAC | 38 |
| Xist_32 | TGCATGAAACTGAACAATTTAAACCTGGAGCTGGTAAT | 38 |
| Xist_33 | GTTCTTAAGACCAATTCAGAACAAAGGCAGGTTGCCCT | 38 |
| Xist_34 | TAAAACAGGTTTGACCTTTTCCTTCACTCTTCCTCCTG | 38 |
| Xist_35 | TCCCACCCTCTGTGAGTGATTTAAAAACGGAAAAGGTC | 38 |
| Xist_36 | AAAGCCCAGCCAGGCCTACATTTAGAGAAATTTTAAAA | 38 |
| Xist_37 | AAATTTTTCCTTTCAATTTTGGCCA | 25 |
| Xist_38 | ATTATTTCCAATTTTTATTTTATTC | 25 |
| Xist_39 | TTAAAACTTATCCTCTTTTGAGGAACAAGTTTTCTTGTAGCATGGTAA | 48 |
| Xist_40 | AATGTTAAGCTCCTCTTTTGAGGAACAAGTTTTCTTGTTGTTTTCATC | 48 |
| Xist_41 | CACTGATATTTCCTCTTTTGAGGAACAAGTTTTCTTGTATTCTACTAT | 48 |
| Xist_42 | AAAAAGCCCTTCCTCTTTTGAGGAACAAGTTTTCTTGTTCTTGAGCAG | 48 |
| Xist_43 | ATTTGATGATTCCTCTTTTGAGGAACAAGTTTTCTTGTAAAAAGGAGA | 48 |
| Xist_44 | ACATTTCTAATCCTCTTTTGAGGAACAAGTTTTCTTGTGGTATAATTA | 48 |
| Xist_45 | GAGAAGCTGTTGCATGAGAAATCATGTCTCCATCTCCA | 38 |
| Xist_46 | TTTTGCTATGCGTTATCTGAGGATTGTTTCTGAAAGAG | 38 |
| Xist_47 | ATCTATTTAGGCAGTGTATGTATGTGTCAGCATGTAAA | 38 |
| Xist_48 | AAGTAAAGAACTGAAAATAGACAAACCTTGTAAATGCA | 38 |
| Xist_49 | CTTCAAAACCAATTGTGGCTCAAGTGTAGGTGGTTCCC | 38 |
| Xist_50 | CAAGGCTGGTACCAATGAGACTGGGGTTTGGGAATTAG | 38 |
| Xist_51 | TTGGTCATCATCCCTCCTGCTGCCC | 25 |
| Xist_52 | AGCAGTGGTCAGTCATTTTTCATGA | 25 |
| Xist_53 | GGGATGGACTTCCTCTTTTGAGGAACAAGTTTTCTTGTAGGAAAATGA | 48 |
| Xist_54 | AGTGTCTATTTCCTCTTTTGAGGAACAAGTTTTCTTGTATAAGGAGAG | 48 |

| | | |
|---|---|---|
| Xist_55 | TTCCTGTAACTCCTCTTTTGAGGAACAAGTTTTCTTGTTTAAGGAGCA | 48 |
| Xist_56 | AGCTCACTACTCCTCTTTTGAGGAACAAGTTTTCTTGTAAAACCCAGC | 48 |
| Xist_57 | TACCTGCTTATCCTCTTTTGAGGAACAAGTTTTCTTGTTCGTAGTGGC | 48 |
| Xist_58 | CAGAGTGGTATCCTCTTTTGAGGAACAAGTTTTCTTGTGAAGAGATAC | 48 |
| Xist_59 | GGAGTAGGAATTAAACCACACAATGTTATTTAGGGACT | 38 |
| Xist_60 | AAGCCATGCCCCTAACAAGAAAACAAGCCAAAAGGAAA | 38 |
| Xist_61 | GTATTAGGCATTCTCTGGGAAGGCATGCATTTTTTTCC | 38 |
| Xist_62 | CATGTCTCTGGGGCCAAAAACCTTATACCAAGTACCTA | 38 |
| Xist_63 | TTGGCACCCGAATATATTTGTAGAATGAATGAATACAT | 38 |
| Xist_64 | GAAAAAAAATAAACAGTAACCTTTCTCCTATATTCTAC | 38 |
| Xist_65 | TTTCCAAGCCAATTAATAAGCAAGT | 25 |
| Xist_66 | GTCTTTTCGTCATGATTTTTTTTGT | 25 |
| Xist_67 | TTTCTGTTTATCCTCTTTTGAGGAACAAGTTTTCTTGTGGATTTAACA | 48 |
| Xist_68 | AAATGGTTGATCCTCTTTTGAGGAACAAGTTTTCTTGTGATAACAGTC | 48 |
| Xist_69 | ACTTCTGTTTTCCTCTTTTGAGGAACAAGTTTTCTTGTGATGAAGAGT | 48 |
| Xist_70 | ATCACTTCATTCCTCTTTTGAGGAACAAGTTTTCTTGTTCCATTTTTG | 48 |
| Xist_71 | TGTTTTTGTTTCCTCTTTTGAGGAACAAGTTTTCTTGTGCATCTCCAA | 48 |
| Xist_72 | GTCAGAATAATCCTCTTTTGAGGAACAAGTTTTCTTGTATGCCTTTTG | 48 |
| Xist_73 | GAGCAGATATATTTCATTTAGCATTTAGTATCATCTTC | 38 |
| Xist_74 | ATCAATACTCGTATGAACGAAAAAATAAAAAGCCCTCT | 38 |
| Xist_75 | CTTATTCCCACTCTACAACGCATGTCAAAGGTGATCTG | 38 |
| Xist_76 | TTTAGTTTTTCCTTAGTATCGAACATATCACAGCTACT | 38 |
| Xist_77 | CAATGAAGTTTCTTCTCAACTAAAGAAACACAGTCTCT | 38 |
| Xist_78 | TAGAGAATTTGTTCCTGTGTTTCCACCATAAGATAAAT | 38 |
| Xist_79 | GAGATAAAGTGCACTTTGGTTTACG | 25 |
| Xist_80 | TCTTTCAAGCACTTAAGAACAGCAC | 25 |
| Xist_81 | TTTATTCAGTTCCTCTTTTGAGGAACAAGTTTTCTTGTTTTAATAAAT | 48 |
| Xist_82 | TTTTACTAAATCCTCTTTTGAGGAACAAGTTTTCTTGTTTGGTAAATG | 48 |
| Xist_83 | CAAAAGAATCTCCTCTTTTGAGGAACAAGTTTTCTTGTTTTTTTTTAT | 48 |

| | | |
|---|---|---|
| Xist_84 | TGCTCATATGTCCTCTTTTGAGGAACAAGTTTTCTTGTTCTTCCTGTC | 48 |
| Xist_85 | TCTAAGCCAGTCCTCTTTTGAGGAACAAGTTTTCTTGTACCAATGAGC | 48 |
| Xist_86 | AAATACCTTTTCCTCTTTTGAGGAACAAGTTTTCTTGTAAAACTAGTT | 48 |
| Xist_87 | GTTACATCTTGAACCATTTAACTGTAATAAAAGCAGAA | 38 |
| Xist_88 | TGTTTAGTTAATGAATTAAAGAACAAACCCTGAGCCCT | 38 |
| Xist_89 | TTTATCAGTCTCCTGGCTTTAAACTAAGCCAATGAGGA | 38 |
| Xist_90 | AGTGATTTGGGGGATTCCTGAAACTAGGAAAAATGTCT | 38 |
| Xist_91 | CTTTATTTTGAAAGAAACTTGCATTTTTCTTTCTTTTT | 38 |
| Xist_92 | TTTTTTTTTTTTAATCTCAAAGGCAATTGAGTGGGTCT | 38 |
| Xist_93 | TCTGGGCCAGACCTATTTAATTTACGAAACATAGTACC | 38 |
| Xist_94 | TTGCAGAGAATAGGCATTGAAATATTATTTAAACAATC | 38 |
| Xist_95 | AAACCAAAGATGTTCTTCTATCTTCAGCTGTCAGTGAT | 38 |
| Xist_96 | CTAATGCCCTCATCTCTCTTATCCTCAGGACCCAGAAT | 38 |
